# SCOPE: Gaussian-regularized graph learning for enzyme catalytic residue prediction

**DOI:** 10.64898/2026.11.20.768221

**Authors:** Zixu Ran, Tong Pan, Yumeng Zhang, Xudong Guo, Yi Hao, Heyun Sun, Jiayu Wang, Xuting Sun, Zhuoqian Li, Xin Cui, Shiwen Zhou, Zhenhao Wang, Jiangning Song, Guimin Zhang, Fuyi Li

## Abstract

Accurate identification of catalytic residues is central to understanding enzyme mechanisms and engineering biocatalysts, yet annotation remains challenging because experimentally validated sites are sparse. Here, we present SCOPE, a graph learning framework for catalytic residue prediction that integrates evolutionary, physicochemical, and three-dimensional structural information at the residue level. SCOPE represents enzymes as residue-level structural graphs and incorporates a Gaussian auxiliary training objective that aggregates structure-aware representations along sequence order. Evaluations across five benchmark datasets, CATH family-disjoint retraining and the external Cata-loDB benchmark assess predictive performance and generalization. Structural analyses associate complete catalytic-residue recovery with compact active-site arrangements. We further validated SCOPE prospectively across four enzymes using site-directed mutagenesis and activity assays, con-firming multiple predicted residues as functionally important. These results support SCOPE as a framework for catalytic residue annotation and prioritizing functional sites for experimental investigation.

## Introduction

Enzymes catalyse a wide range of biochemical transformations essential to cellular function and pro-vide versatile catalysts for chemical synthesis and industrial biotechnology ^1, 2^. Their catalytic activities depend on a relatively small set of residues arranged within three-dimensional active-site environments. These residues contribute to reaction chemistry through mechanisms including general acid-base catalysis, covalent catalysis, metal coordination and transition-state stabilisation ^3^. Identifying catalytic residues therefore supports mechanistic interpretation and helps prioritise positions for functional investigation and enzyme engineering.

Catalytic residues have traditionally been characterised through experimental methods such as site-directed mutagenesis, kinetic characterisation, and high-resolution structural analysis ^4^. Although high-throughput methods have expanded the scale of functional measurements, detailed mechanistic characterisation remains experimentally demanding. Many protein sequences lack experimental characterisation, and their functional annotations therefore rely on computational inference ^5^. Curated resources such as M-CSA provide residue-level mechanistic information for experimentally characterised enzymes ^3^. Accurate and scalable computational methods for catalytic residue prediction are therefore needed to prioritise functional sites, guide experimental validation and accelerate enzyme discovery.

Computational identification of catalytic residues remains an important challenge, particularly for enzymes with incomplete functional annotation. Homology-based strategies provide an established route to functional-site identification by transferring information from annotated proteins. Catalytic residues and their local structural arrangements can remain conserved despite substantial sequence divergence, making both homologous templates and three-dimensional motif searches informative for catalytic-site annotation ^6^. These motif-based approaches explicitly exploit the spatial organisation of functional residues and therefore provide important context for structure-based prediction. More recent machine learning and deep learning approaches (**Table S1**) integrate evolutionary, sequence and structural features to generate residue-level predictions ^7, 8, 9, 10^, while graph neural networks provide a natural framework for representing three-dimensional residue relationships ^11, 12^. Building on these approaches, we investigate whether multimodal feature integration combined with a Gaussian auxiliary training objective can improve catalytic-residue prediction.

Here we introduce SCOPE (**S**moothing-based **CO**nvolutional graph approach for catalytic residue **P**rediction in **E**nzymes), a graph learning framework for catalytic residue prediction with a Gaussian auxiliary training objective. SCOPE represents each enzyme as a residue-level structural graph that integrates evolutionary profiles, protein language model embeddings, physicochemical descriptors and three-dimensional contact information. SCOPE incorporates a Gaussian auxiliary objective that aggregates structure-aware residue representations along sequence order during training. Empirical evaluations across diverse benchmark datasets demonstrate competitive predictive performance, with gains varying across datasets and metrics. Further analyses reveal that SCOPE preferentially resolves enzymes with spatially compact catalytic microenvironments, particularly hydrolase and peptidase-related enzymes, with Gaussian smoothing supporting the recognition of otherwise missed residues within these locally clustered catalytic contexts. Finally, we demonstrate the framework’s practical utility through prospective prediction and experimental validation of functionally critical residues in four low-homology enzymes. Together, these results support SCOPE as a framework for catalytic-residue annotation and prioritisation of functional sites for experimental investigation and enzyme engineering.

## Results

### SCOPE integrates structure-aware graph learning with an auxiliary Gaussian smoothing objective

The overall architecture of SCOPE is shown in **Fig. 1**. SCOPE was designed to predict catalytic residues by modelling enzymes as residue-level structural graphs that integrate sequence-derived, evolutionary and three-dimensional structural information. The framework consists of two main components: a graph construction module (**Fig. 1a**) and a feature-fusion and prediction module (**Fig. 1b**). The graph construction module encodes each enzyme as a residue-level contact graph, whereas the prediction module combines multimodal residue features within a DeepGCN framework enhanced by a Gaussian smoothing constraint. This design enables SCOPE to model catalytic residues not as isolated sequence positions, but as spatially organised functional sites embedded within local structural environments.

**Fig. 1.**
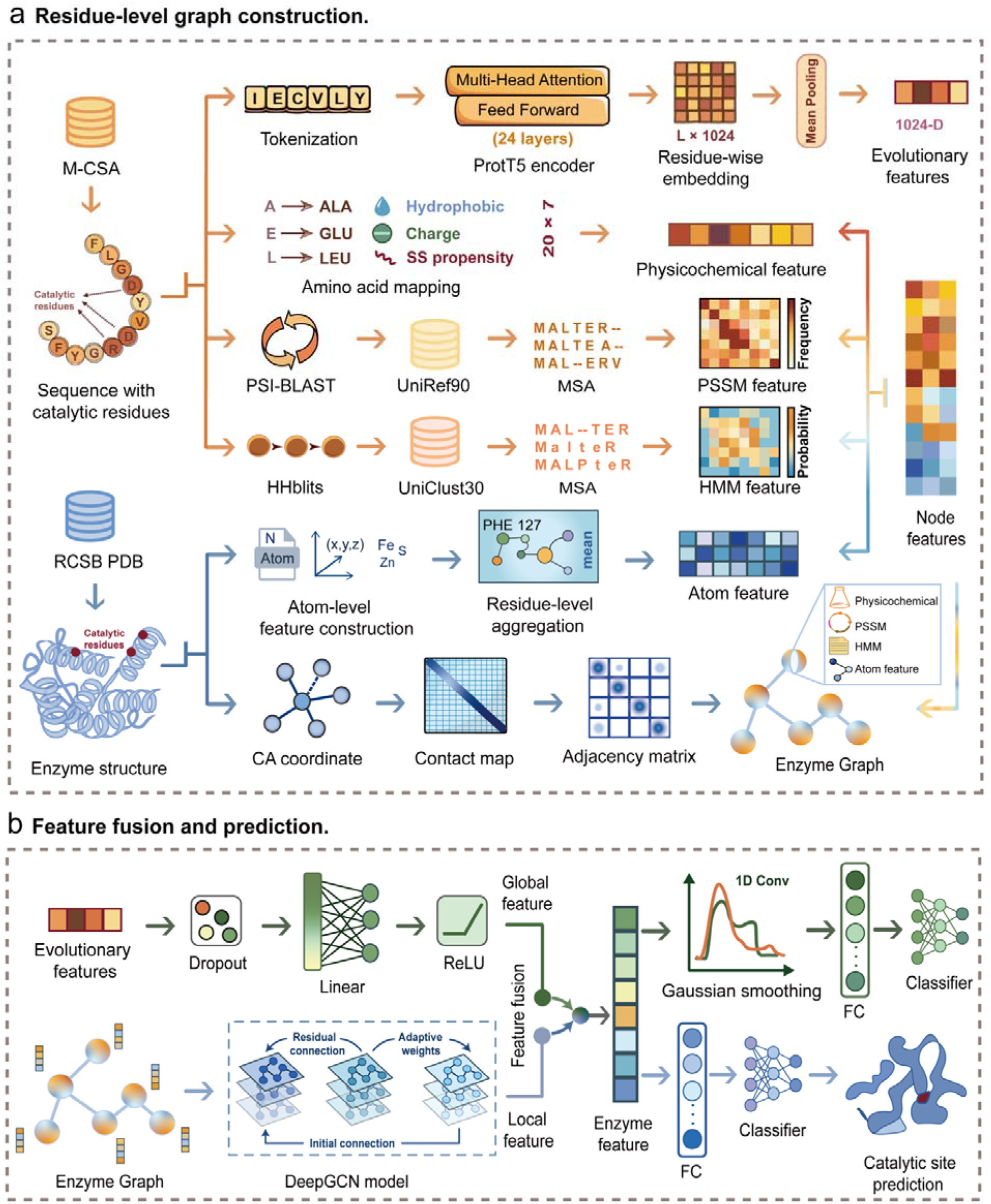
Overall framework of SCOPE for catalytic residue prediction. **a**, Residue-level graph construction. For each enzyme, sequence-derived features, including physicochemical descriptors, PSSM profiles, and HMM profiles, are combined with structure-derived atomic features to generate residue-level node representations. A contact map derived from the three-dimensional enzyme structure defines edges between residues within a predefined spatial distance threshold, producing a residue-level structural graph. **b**, Feature fusion and prediction. The residue graph is processed by a DeepGCN to learn local structure-aware representations, whereas ProtT5 embeddings provide complementary global evolutionary information. These two representations are projected, fused, and transformed into final residue embeddings, which are then mapped to catalytic-residue probabilities. A Gaussian smoothing operation is applied as an auxiliary training constraint to reinforce locally coherent catalytic signals within structurally organised residue neighbourhoods.

As shown in **Fig. 1a**, SCOPE first aligns each enzyme sequence with its corresponding three-dimensional structure and extracts residual-level features from both sequence and structural modalities. Catalytic residue annotations are obtained from the M-CSA database ^3^. To capture sequence-derived evolutionary context, SCOPE uses residue-wise embeddings generated by the pretrained protein language model ProtT5 ^13^, together with position-specific scoring matrix (PSSM) profiles generated by PSI-BLAST and hidden Markov model (HMM) profiles generated by HHblits _14._

Structural information is incorporated by retrieving the corresponding three-dimensional enzyme structures from the RCSB PDB ^15^ and extracting atom-level structural features. SCOPE then constructs a residue-residue contact map from inter-residue spatial distances. Residues are connected when their spatial distance falls below a predefined threshold (6Å), yielding a graph in which nodes represent residues and edges represent spatial proximity. PSSM profiles, HMM features, atomic structural features, and amino acid physicochemical descriptors are concatenated to form residue-level node features, while the contact map defines the graph topology. Through this representation, SCOPE embeds each enzyme within a structure-aware graph that jointly captures evolutionary signals, biochemical residue properties and three-dimensional spatial constraints.

In the feature-fusion and prediction module (**Fig. 1b**), the residue-level graph is processed by DeepGCN to learn local structure-aware residue representations. In parallel, ProtT5-derived residue embeddings are projected into the same latent space through a linear transformation, providing complementary global evolutionary information. The local graph-derived structural features and the ProtT5-derived evolutionary features are then concatenated and integrated through a linear fusion layer to generate the final residue-level enzyme representation. This fusion strategy allows SCOPE to combine conserved sequence information with local active-site geometry, both of which are critical for identifying catalytic residues in enzymes with limited homology to annotated proteins.

To complement structure-aware graph learning, SCOPE incorporates Gaussian smoothing as an auxiliary training objective. This operation aggregates graph-derived residue representations within a local sequence window, assigning progressively smaller weights to positions farther from the central residue. An auxiliary prediction loss provides additional supervision alongside the main prediction objective, encouraging the learned representations to support catalytic-residue recognition under this local aggregation. Three-dimensional relationships are therefore captured through the graph topology, whereas Gaussian smoothing operates along sequence order. Final catalytic-residue probabilities are produced by the main prediction branch, without inference-time smoothing or filtering.

### SCOPE outperforms state-of-the-art approaches through spatial catalytic-context modelling

To assess whether spatial catalytic-context modelling improves catalytic residue prediction, we first compared SCOPE with a sequence-only one-dimensional convolutional neural network (CNN) baseline (**Fig. 2a**). Model performance was evaluated using standard classification metrics, including precision, recall, AUPRC, and MCC. Across different independent test datasets, graph-based modelling consistently outperformed the sequence-only baseline in most metrics. These results support incorporating three-dimensional residue neighbourhoods into catalytic-residue prediction.

**Fig. 2.**
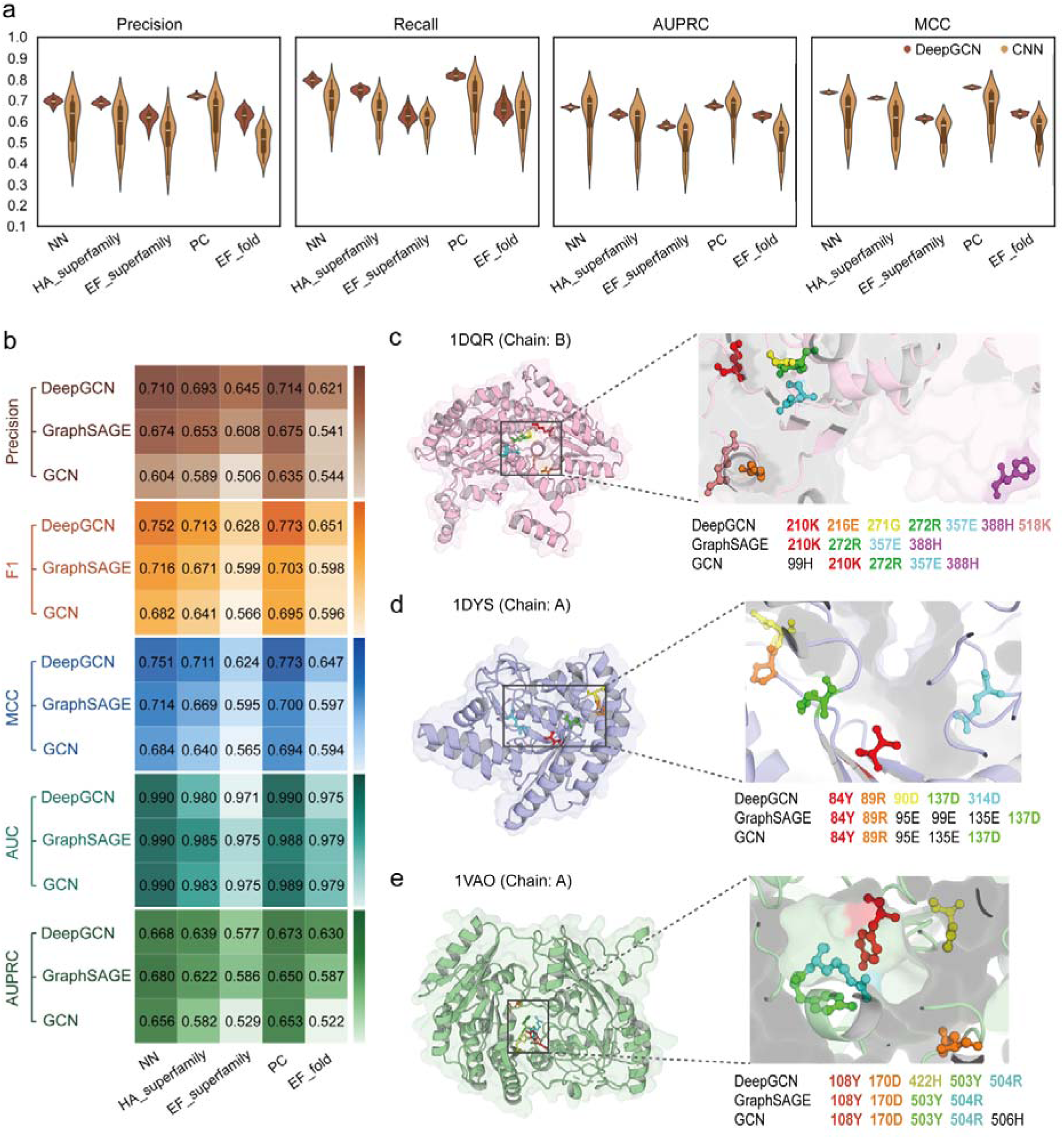
Performance benchmarking and structural comparison. **a,** Performance comparison between the sequence-only CNN baseline and SCOPE on five test sets: NN, HA_superfamily, EF_superfamily, PC and EF_fold. Models were evaluated using Precision, Recall, AUPRC and MCC. **b,** Comparison of graph neural network backbones, including GCN, GraphSAGE and DeepGCN, across the same five test sets. **c-e,** Structural visualisation of predicted catalytic residues in three representative enzymes: 1DQR (chain B), 1DYS (chain A) and 1VAO (chain A). Predicted catalytic residues from SCOPE, DeepGCN, GraphSAGE and GCN are highlighted on the same structures for comparison.

We next examined the contribution of graph neural network architecture by comparing representative graph backbones, including GCN ^16^, GraphSAGE ^17^, and DeepGCN ^18^. Catalytic residues often form spatially coordinated groups ^19^, motivating graph-based aggregation of structural context. Residual connections and identity-preserving mechanisms can mitigate the over-smoothing associated with repeated graph propagation ^20^. Consistent with this design rationale, DeepGCN showed the most favourable overall performance across the five benchmark datasets (shown in **Fig. 2b**, including NN, HA_superfamily, EF_superfamily, PC, and EF_fold). For example, on the PC dataset, DeepGCN achieved an *F*_1_score of 0.773, outperforming GCN (0.695) and GraphSAGE (0.703)

To further examine whether these performance gains reflected biologically meaningful active-site recognition, we visualised predictions for three representative enzymes with well-characterised catalytic mechanisms and distinct structural contexts. As shown in **Fig. 2c**, for phosphoglucose isomerase (PGI, PDB: 1DQR), a key glycolytic enzyme ^21^, catalysis depends on a group of conserved residues positioned within the active pocket that cooperatively mediate proton transfer and stabilise substrate conformations. In this process, Lys residues act as general bases, whereas His and Glu residues contribute to substrate positioning and stabilisation of reaction intermediates. In **Fig. 2d**, for endoglucanase Cel6B (PDB: 1DYS), a typical cellulose-degrading hydrolase whose catalytic centre is dominated by acidic residues distributed along the substrate-binding groove ^22^. In **Fig. 2e**, for vanillyl-alcohol oxidase (PDB: 1VAO), an FAD-dependent oxidoreductase whose catalytic activity relies on key residues surrounding the cofactor-binding site, among which His residues interact covalently or non-covalently with FAD to stabilise the cofactor conformation ^23^. Structural visualisation shows that the DeepGCN-based model more accurately recovered annotated catalytic residues within the active-site region, whereas GCN and GraphSAGE either missed true catalytic residues or assigned catalytic predictions to residues displaced from the active centre. We also examined the influence of the protein language model used to generate residue embeddings. SCOPE variants using frozen ProtT5, ESM-2, or ProtBERT-BFD embeddings were independently retrained under the same downstream training and evaluation protocol. ProtT5 achieved the highest AUPRC across all five test sets under the evaluated extraction settings, supporting its use in the present implementation (**Table S14**).

In addition, we systematically compared SCOPE with two recently proposed graph-based catalytic-site predictors, SCREEN ^24^ and AEGAN ^25^ (**Table S17**). SCREEN was trained on the same training and validation protein split as SCOPE, and AEGAN was retrained on this split for the revised comparison. Details of baseline training and matched-residue evaluation are provided in Supplemental Information, “Matched baseline comparison and methodological increments” (**Table S17**). We also evaluated published Squidly and EasIFA models on matched subsets of the five original benchmarks; these comparison values were calculated from predictions generated in this study rather than taken from the original publications (**Tables S30-S31**). We further evaluated SCOPE on CataloDB against published Squidly ^26^and EasIFA ^27^models and a training-restricted BLAST baseline. On the common cohort of 121 records representing 120 unique sequences, SCOPE achieved sequence-group macro-AUPRC/AP of 0.6881/0.7322, compared with 0.6080/0.6410 for Squidly, 0.5775/0.6507 for EasIFA and 0.5909/0.2250 for BLAST (**Table S32**). Under CATH homologous-superfamily-disjoint retraining with three fixed seeds, SCOPE exceeded SCREEN in per-protein AUPRC by 0.0467 (95% CI, 0.0285–0.0654; Holm-adjusted P = 2.94 × 10^-6^), whereas its difference from the no-Gaussian variant was not significant (**Table S25**). We additionally evaluated an EnzyMM structural motif baseline using templates constructed from the training partition, with comparisons performed on matched evaluable residues (**Table S29**). Across different test datasets, SCOPE achieved overall competitive or superior performance in terms of Precision,*F*_1_, AUC, and MCC. Ablation results assessing the contribution of the Gaussian auxiliary objective are provided in **Table S2**.

### SCOPE complete-recovery enzymes show compact catalytic spatial organisation and differences in functional composition

To investigate the biological contexts in which SCOPE provides additional predictive value, we first systematically identified enzymes for which SCOPE correctly recovered all annotated catalytic residues without false-positive predictions, whereas at least one of the competing methods, SCREEN or retrained AEGAN, made a classification error on the matched evaluation positions. This screening yielded 119 dataset-level SCOPE complete-recovery records, corresponding to 95 distinct protein-chain records after deduplication across test sets (**Table S3**). Compared with the remaining test enzymes, SCOPE complete-recovery enzymes displayed shorter local distances between annotated catalytic residues. The median of the per-record mean nearest-neighbour distances between catalytic residues was shorter in SCOPE complete-recovery enzymes than in other test enzymes (6.61Å versus 7.30Å; n = 77 versus n = 204 records with evaluable distance measurements; two-sided Mann-Whitney U test, P = 0.0241). The median of the per-record mean pairwise distances among catalytic residues was also lower in SCOPE complete-recovery enzymes than in other test enzymes, although this difference was not statistically significant (10.12 Å versus 10.62 Å; P = 0.161; **Fig.3a**). Consistent with this observation, annotated catalytic residues showed pronounced spatial clustering across the full benchmark set (**Fig. S1a**), supporting the biological rationale for modelling catalytic residues as spatially coordinated active-site ensembles.

We next evaluated whether residues predicted by different methods within the SCOPE complete-recovery enzymes recapitulated the biophysical properties of annotated catalytic residues. SCOPE-predicted residues coincided with annotated catalytic residues by the cohort-selection criterion and therefore had identical distributions of per-record average hydrophobicity, charged-residue fraction, B-factor z-score and distance to the pocket-centre. SCREEN predictions were also broadly similar to annotated catalytic residues, although with modest deviations. By contrast, retrained AEGAN-predicted residues showed more pronounced differences, including higher hydrophobicity, a lower fraction of charged residues, higher B-factor z-scores and greater distances from the pocket centre (**Fig. 3b**). These comparisons describe the spatial and physicochemical properties of predicted residues within the selected cohort rather than providing an independent assessment of comparative accuracy. Consistently, global physicochemical analyses showed that SCOPE-predicted residues recapitulated the amino acid composition, charge distribution, hydrophilicity, structural rigidity and pocket localisation characteristic of annotated catalytic residues (**Fig. S2**).

**Fig. 3.**
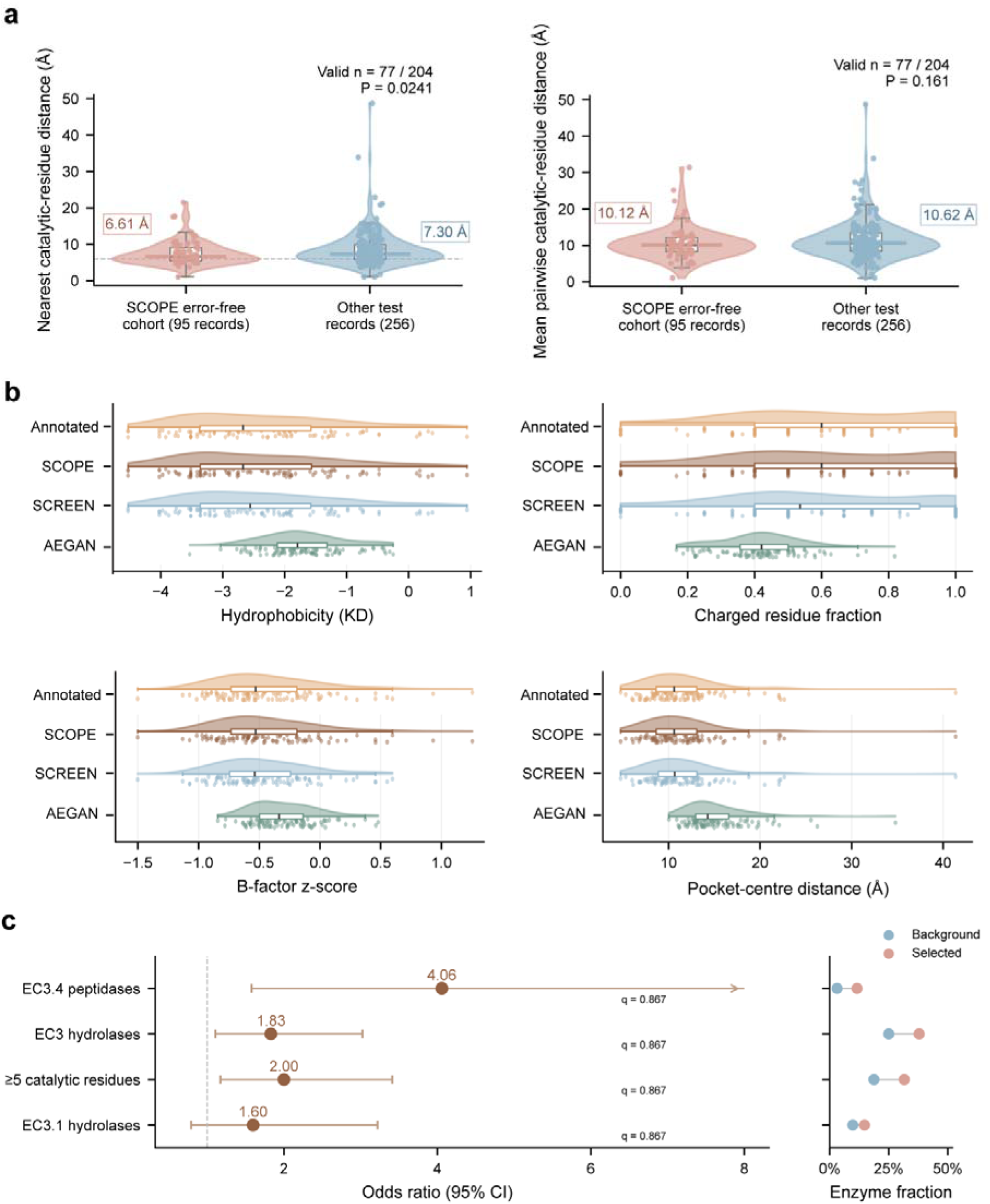
Spatial, biophysical and functional characteristics of SCOPE complete-recovery enzymes. **a**, Spatial organisation of annotated catalytic residues in SCOPE complete-recovery enzymes and other test enzymes. Violin plots show the record-level distributions of mean nearest-neighbour catalytic-residue distance (left) and mean pairwise catalytic-residue distance (right). Dots indicate individual records; boxes show medians and interquartile ranges, with 1.5×IQR whiskers. Distances were evaluated for 77 complete-recovery and 204 background records. P values were calculated using two-sided Mann-Whitney U tests. **b**, Biophysical properties of annotated catalytic residues and residues predicted by SCOPE, SCREEN and retrained AEGAN in SCOPE complete-recovery enzymes, including Kyte-Doolittle hydrophobicity, charged residue fraction, B-factor z-score and distance to the pocket centre. Half-violins and dots show per-record distributions and values; box plots are defined in panel a. **c**, Functional category comparisons of SCOPE complete-recovery enzymes. Left, odds ratios with 95% confidence intervals for selected functional categories. The arrow indicates a confidence interval extending beyond the displayed axis range. Right, fractions of records assigned to each category in the background set and SCOPE complete-recovery set. BH correction covered 878 annotation terms; all displayed q values were ≥0.05.

To determine whether SCOPE complete-recovery enzymes were associated with specific functional categories, we performed enrichment analysis using the 95 SCOPE complete-recovery records as the foreground set and the 256 remaining test records as the background. EC3.4 peptidases were more frequent in the SCOPE complete-recovery cohort (11/95 versus 8/256; odds ratio = 4.06), but this association did not reach statistical significance after multiple-comparison correction (two-sided Fisher’s exact P = 0.00560; Benjamini-Hochberg-adjusted P = 0.867 across 878 tested annotation terms). The corresponding category frequencies and effect estimates are shown in **Fig. 3c** (**Table S23**).

A broader geometric classification of catalytic-residue clusters is provided in **Fig. S3**. Illustrative comparisons of predictor outputs for two enzymes with known catalytic architectures are provided in **Fig. S6**.

### Gaussian smoothing stabilizes catalytic-residue recognition within local active-site contexts

The preceding analyses described compact catalytic-site organisation within the SCOPE complete-recovery cohort. Building on this observation, we next asked whether these SCOPE complete-recovery features were associated with the Gaussian smoothing component of the model. Among the 95 SCOPE complete-recovery records, full SCOPE achieved error-free classification on the matched evaluation positions in all records, whereas the model trained without the Gaussian auxiliary objective retained error-free classification for only 82 records. The remaining 13 records were therefore classified as Gaussian smoothing (GS)-dependent records(**Fig. 4a**). Of these, five contained false-negative catalytic-residue predictions in the model without the Gaussian auxiliary objective, whereas the remaining eight contained false-positive predictions only.

**Fig. 4.**
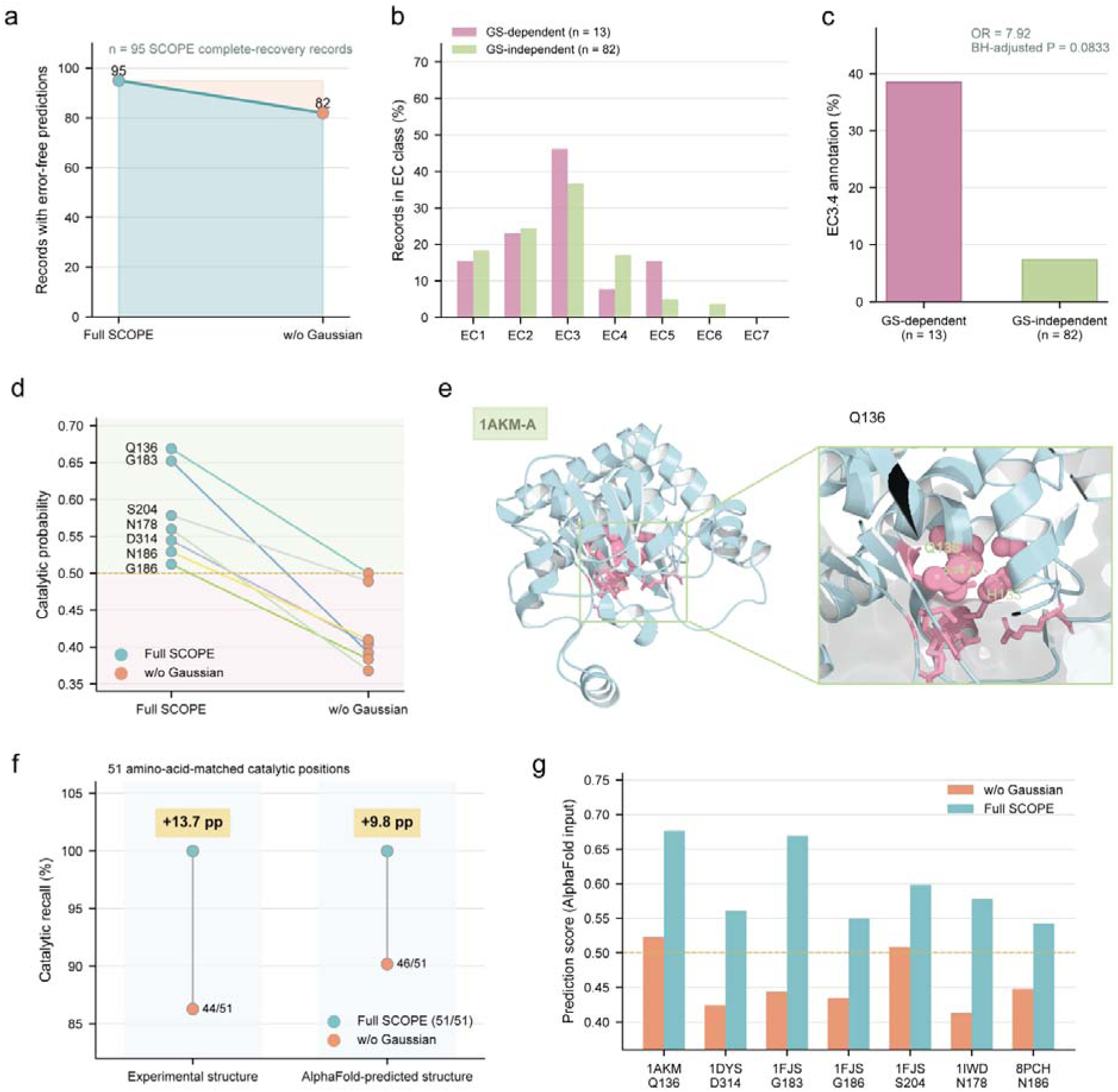
Gaussian smoothing stabilises catalytic-residue recognition within local active-site contexts. **a**, Record-level ablation of the Gaussian auxiliary objective in 95 SCOPE complete-recovery records. Complete recognition denotes error-free classification on the matched evaluation positions, using a prediction threshold of 0.5. **b**, Distribution of GS-dependent and GS-independent records across first-level EC classes. GS-dependent records (n = 13) lost complete recognition in the model trained without the Gaussian auxiliary objective; GS-independent records (n = 82) retained complete recognition without this objective. **c**, Frequency of EC3.4 peptidase/protease annotations in GS-dependent records compared with GS-independent records. The association was not significant after BH correction across 13 EC comparisons (adjusted P = 0.08334). **d**, Paired prediction scores of seven Gaussian-dependent catalytic residues. Each line represents an annotated catalytic residue that was correctly identified by full SCOPE but fell below the 0.5 threshold in the model trained without the Gaussian auxiliary objective. The w/o Gaussian score for Q136 was 0.499943. **e**, Structural example of a Gaussian-dependent residue in 1AKM chain The residue Q136 is located 5.94Å from the annotated catalytic residue H133, placing it within the 6Å residue-neighbourhood threshold used for graph construction in SCOPE. **f**, Catalytic-residue recall using experimental and AlphaFold-predicted structures as input, evaluated on 51 paired catalytic positions with matching amino acid identities. Yellow labels indicate the recall difference between full SCOPE and w/o Gaussian. **g**, Prediction scores of the seven Gaussian-dependent residues selected using experimental structures, using AlphaFold-predicted structures as input. The dashed line indicates the prediction threshold of 0.5. Q136 and S204 exceeded the threshold in both models.

To determine whether this dependence reflected a broad enzyme-class effect, we compared the distributions of GS-dependent enzymes and GS-independent enzymes across enzyme EC1-EC7 classes. Although EC3 hydrolases were modestly more frequent among GS-dependent enzymes (**Fig. 4b**), EC3.4 peptidases/proteases also showed a higher observed frequency in this group. Specifically, 38.5% of GS-dependent records carried EC3.4 annotations, compared with 7.3% of GS-independent records (5/13 versus 6/82; **Fig. 4c**). This association did not reach statistical significance after correction across the 13 EC comparisons (odds ratio = 7.92; two-sided Fisher’s exact P = 0.00641; BH-adjusted P = 0.08334; **Table S24**). The higher frequency of peptidases and proteases among GS-dependent enzymes suggests a possible role for the Gaussian auxiliary objective in recognising residues within locally organised active-site environments.

At the residue level, the 13 GS-dependent enzymes contained 52 annotated catalytic residues, of which seven were missed by the model trained without the Gaussian auxiliary objective. These residues scored above the 0.5 decision threshold in full SCOPE but fell below this threshold in the model without the auxiliary objective. The Gaussian-dependent residues included 1AKM chain A Q136, 1DYS chain A D314, 1FJS chain A G183, G186, and S204, 1IWD chain A N178, and 8PCH chain A N186 (**Fig. 4d**). Thus, the Gaussian auxiliary objective affects both the recovery of annotated catalytic residues and the avoidance of false-positive predictions in the evaluated models.

A representative structural case further illustrates the local context of a residue whose classification differed between the models. In 1AKM chain A, the Gaussian-rescued residue Q136 was located 5.94Å from the annotated catalytic residue H133, placing it within the 6Å residue-neighbourhood threshold used for graph construction in SCOPE (**Fig. 4e**). In the model trained without the Gaussian auxiliary objective, the prediction score of Q136 was just below the decision threshold and the residue was classified as a false negative, whereas full SCOPE classified it as a true positive. This spatial relationship describes the structural context of Q136. Consistent with the incorporation of structural context by the graph network, GNNExplainer ^28^ analysis showed that SCOPE focuses on locally connected structural regions around active sites (**Fig. S4**).

We finally tested whether the contribution of Gaussian smoothing was retained when predicted structures were used as input. We replaced the experimental structures of the 13 GS-dependent enzymes with AlphaFold-predicted structures ^29^. The paired analysis retained 51 annotated catalytic positions with matching amino acid identities and available coordinates in both structural inputs; one position was excluded because of an amino acid mismatch. Full SCOPE achieved complete catalytic-residue recall with both experimental and AlphaFold-predicted structures (51/51 in each setting), whereas w/o Gaussian reached 86.3% (44/51) and 90.2% (46/51) recall, respectively. Thus, full SCOPE had higher recall by 13.7 and 9.8 percentage points under the two structural-input settings (**Fig. 4f**). Residue-level score analysis further showed that, among the seven Gaussian-rescued residues identified using experimental structures, all sites had higher full SCOPE scores than w/o Gaussian under AlphaFold input. Five of seven residues retained the strict Gaussian-dependent pattern, with full SCOPE scores ≥0.5 and w/o Gaussian scores <0.5; Q136 and S204 exceeded the threshold in both models and therefore no longer represented strict rescued residues, but still received higher scores from full SCOPE (**Fig. 4g**). Together, these results indicate that the higher catalytic-residue recall of full SCOPE within this selected cohort was retained with AlphaFold-predicted structural inputs. A separate supplementary analysis found comparable SCOPE performance using experimental and AlphaFold structures, with no clear association between catalytic-site pLDDT and prediction performance within the evaluated confidence range (**Tables S15–S16**).

### Experimental validation confirms SCOPE-predicted functional sites across diverse enzymes

To experimentally assess the functional relevance of SCOPE-predicted positions, we selected four industrially relevant enzymes from our in-house enzyme library: tyrosine ammonia lyase (Fc TAL2)^30^, laccase (CotA) ^31^, diacetylchitobiose deacetylase (TGDac) and alkaline phosphatase (rsAP) ^32^. These enzymes span three major catalytic classes widely used in industrial biotechnology, including oxidoreductases, hydrolases and lyases. Importantly, all four enzymes showed low sequence identity to proteins in the training set, including Fc TAL2 (35.03%), CotA (20.00%), TGDac (23.78%) and rsAP (30.99%). Their distinct catalytic mechanisms, cofactor dependencies and active-site architectures allow us to assess the functional relevance of SCOPE-selected positions across different enzyme systems. Training-template correspondences are provided in **Table S22**. In addition, all four enzymes have established expression, purification and activity-assay protocols, and their substrates are commercially available, enabling systematic experimental validation.

SCOPE identified reported catalytic residues and proposed additional candidate positions for functional testing in the four enzymes (**Table S4**). The CotA positions H107 and H155 have established copper-coordinating roles and are not presented as newly discovered catalytic sites. In Fc-TAL2, SCOPE identified the known catalytic residues Y50 and S140 ^30^ and predicted R291, F337, N420 and D489, which were located within the substrate-binding pocket or at subunit-interface-associated regions (**Fig. 5a**). In TGDac, the predicted residues D38, D107, H144 and D146 were positioned near the active pocket (**Fig. 5b**). In CotA, H107A and H155A substitutions eliminated ABTS oxidation activity, consistent with their roles as copper-coordinating histidines required for multicopper oxidase function (**Fig. 5c**). In rsAP, the SCOPE-predicted residue H484 was located near the centre of the active pocket, suggesting a potential role in substrate binding or catalysis (**Fig. 5d**).

**Fig. 5.**
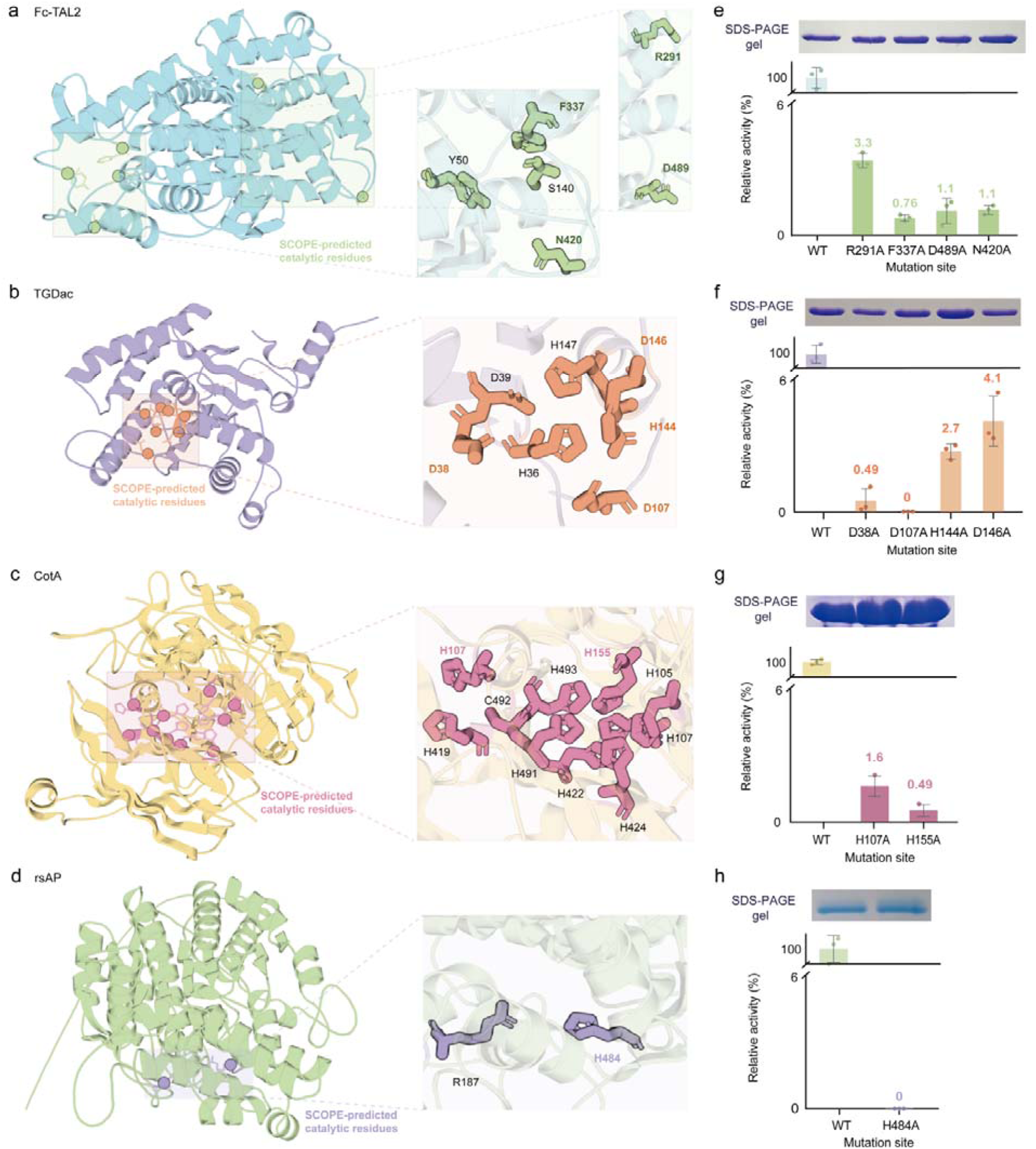
Experimental validation of SCOPE-predicted functional residues across diverse enzymes. **a-d**, Structural visualisation of SCOPE-predicted residues in four low-homology enzymes used for prospective validation: Fc-TAL2 (a), TGDac (b), CotA (c), and rsAP (d), respectively. Black residue labels indicate SCOPE-predicted residues corresponding to previously annotated catalytic residues. **e-h**, Experimental validation of predicted functional residues through site-directed mutagenesis and enzymatic activity assays. Protein expression levels were verified by SDS-PAGE to exclude expression bias. Bars show mean ± SD from three independent experiments (n = 3), with relative activities normalised to the mean activity of the corresponding wild-type enzyme. Broken y-axes are used to visualise both wild-type activity and low residual mutant activities.

We next tested the functional contribution of these predicted sites using site-directed mutagenesis followed by enzymatic activity assays (“Plasmid construction and mutagenesis” and “Protein expression, purification and relative activity assay”). In Fc-TAL2, R291A, F337A, N420A and D489A substitutions abolished or markedly reduced enzymatic activity, supporting the functional importance of these positions (**Fig. 5e**). In TGDac, D38A, D107A, H144A and D146A substitutions strongly impaired catalytic activity, supporting their involvement in active-pocket function (**Fig. 5f**). In CotA, H107A and H155A substitutions eliminated ABTS oxidation activity ^31^, consistent with their roles as copper-coordinating histidines required for multicopper oxidase function (**Fig. 5g**). In rsAP, H484A substantially reduced phosphoester hydrolysis activity ^32^, supporting the functional importance of H484 (**Fig. 5h**). Protein expression and purification of the mutants were verified by SDS-PAGE, and protein concentrations were normalised before activity measurements to minimise expression-level differences as the primary cause of activity loss. Relative activities shown in Fig. 5e-h were derived from three independent experiments and normalised to the mean activity of the corresponding wild-type enzyme. Across the experimentally tested SCOPE-predicted residues, alanine substitution consistently reduced residual activity to 0-4.1% of wild-type activity, providing systematic experimental support for the functional relevance of these predicted sites (**Table S5**). These prospective experiments support the functional importance of SCOPE-predicted residues across four enzymes with distinct catalytic mechanisms.

### Local structural contexts of experimentally validated functional sites

After confirming the functional importance of SCOPE-predicted sites, we next asked whether these experimentally validated sites reflected the local catalytic-context features identified in the benchmark analyses. Across the 11 mutagenesis-validated functional sites from the four enzymes, full SCOPE identified all sites, whereas w/o Gaussian and SCREEN missed different subsets of validated residues (**Fig. 6a**). At the aggregate level, full SCOPE detected all 11 validated sites, whereas w/o Gaussian and SCREEN detected seven (63.6%) and six (54.5%) sites, respectively (**Fig. 6b**). Predicted residue lists for the four enzymes are provided in **Table S6**. Because these sites were selected for experimental testing based on SCOPE predictions, this comparison is descriptive rather than an unbiased benchmark of relative model performance.

**Fig. 6.**
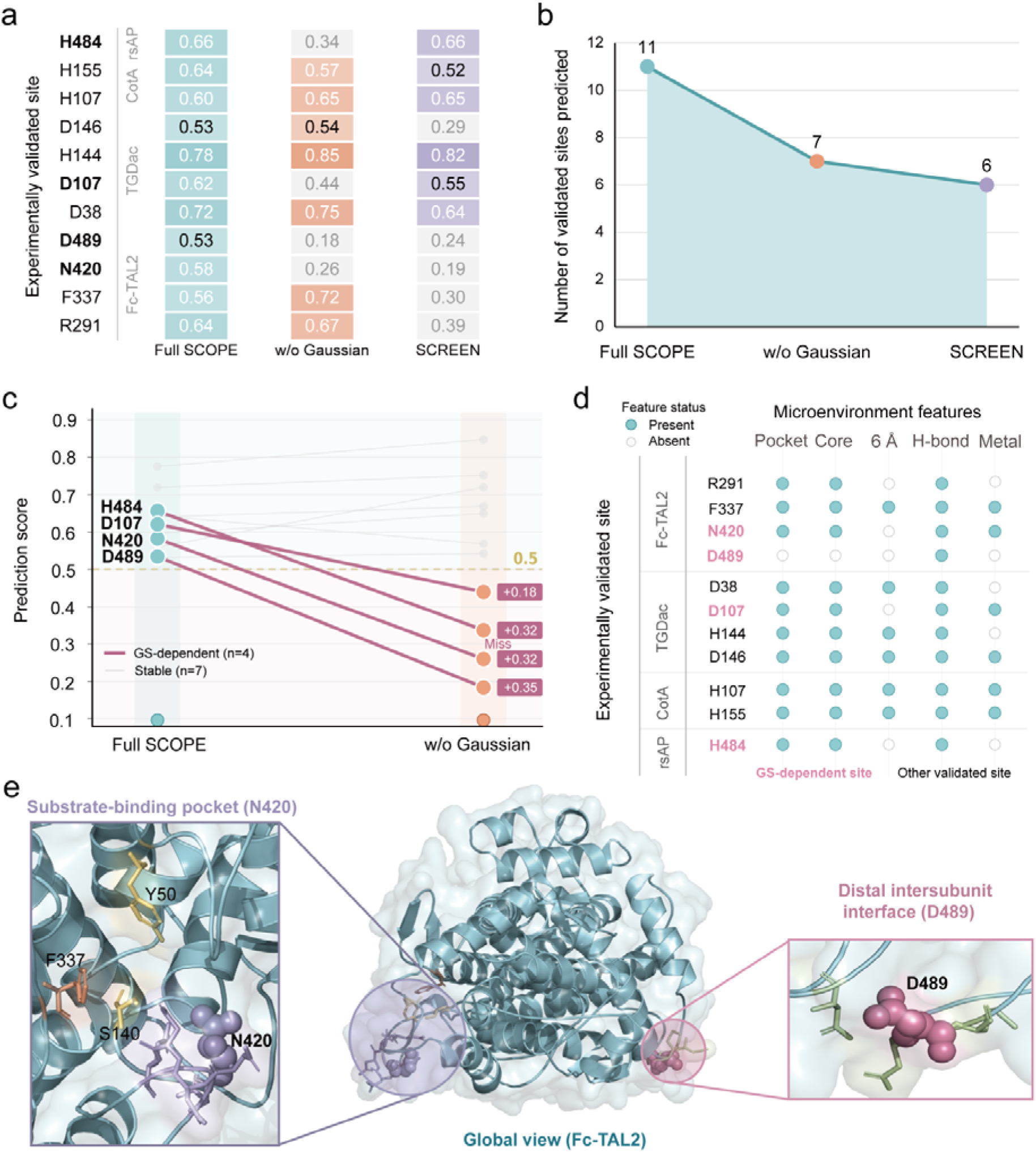
Model predictions and local structural contexts of experimentally validated functional sites. **(a)** Prediction scores of 11 experimentally validated functional sites from four enzymes, Fc-TAL2, TGDac, CotA and rsAP, across Full SCOPE, w/o Gaussian and SCREEN. **(b)** Number of experimentally validated sites identified by each model at a prediction threshold of 0.5. Full SCOPE identified all 11 sites, whereas w/o Gaussian and SCREEN identified seven and six sites, respectively. **(c)** Paired prediction-score comparison between Full SCOPE and w/o Gaussian for the 11 validated sites. Highlighted lines denote four Gaussian-dependent sites, N420, D489, D107 and H484, whose scores fell below the 0.5 threshold in the model trained without the Gaussian auxiliary objective. Grey lines denote sites that remained positive in both settings. **(d)** Local microenvironment features of the 11 experimentally validated sites. Filled teal circles indicate the presence of each feature, open circles indicate absence, and rose labels mark Gaussian-dependent sites. Features include active-pocket localisation, proximity to the catalytic core, presence of a 6 Å spatial neighbour, hydrogen-bond or salt-bridge involvement, and metal-associated environment. **(e)** Structural example of Fc-TAL2. The global structure highlights two experimentally validated sites with distinct local contexts: N420 lies near the substrate-binding pocket together with Y50, S140 and F337, whereas D489 is located at a distal intersubunit-interface-associated region.

Paired score analysis between full SCOPE and w/o Gaussian further identified four Gaussian-dependent validated sites: Fc-TAL2 N420, Fc-TAL2 D489, TGDac D107 and rsAP H484. These residues scored above the 0.5 decision threshold in full SCOPE but fell below the threshold in the separately trained model without the Gaussian auxiliary objective, resulting in missed recognition by the w/o Gaussian model (**Fig. 6c**). Among these sites, Fc-TAL2 D489 showed the largest score decrease relative to full SCOPE, whereas N420, D107 and H484 also crossed below the decision threshold. In contrast, the other seven validated sites remained above threshold in both settings. These observations indicate that the Gaussian auxiliary objective contributed to the identification of four experimentally validated functional sites in the evaluated models.

We then examined the structural context of these 11 validated sites. These 11 sites were broadly associated with active-pocket localisation, catalytic-core proximity, 6Å spatial neighbourhoods, hydrogen-bond or salt-bridge networks, or metal-associated environments (**Fig. 6d**). Notably, the four GS-dependent validated residues, N420, D489, D107 and H484, also exhibited local catalytic-context features, suggesting that Gaussian smoothing may help SCOPE recover functionally relevant residues that are not direct 6 Å neighbours of annotated catalytic residues but are still embedded in supportive structural environments. D489 showed a distinct structural context, with hydrogen-bond network involvement in **Fig. 6d** and localisation at a distal intersubunit interface in **Fig. 6e**, suggesting that its functional effect may arise through structural coupling rather than direct participation in the compact catalytic core. These results indicate that the experimentally tested SCOPE predictions include positions in both active-site-associated and distal structural environments.

The Fc-TAL2 structure provided a representative example of two validated residues with distinct local context. N420 was positioned near the substrate-binding pocket together with the known catalytic residues Y50 and S140, consistent with a role in the local active-site environment. By contrast, D489 was located at a distal intersubunit-interface-associated region, suggesting a possible contribution to catalytic conformation or substrate-channel stability (**Fig. 6e**). For positions outside the active-site core, the observed activity loss may reflect effects on subunit organization, stability or structural communication. These remain possible explanations rather than experimentally established mechanisms, and the tested positions are therefore described as functionally important without assigning a direct catalytic role. Both residues were experimentally validated as functionally critical and showed the experimentally tested SCOPE predictions include positions in both active-site-associated and distal structural environments.

To extend the assessment beyond the selected experimental positions, we compared predictions from the full and no-Gaussian models across a broader enzyme cohort and analysed public PafA mutagenesis data. These analyses characterised changes in catalytic-residue predictions and assessed the coverage of mutation-sensitive positions, with alternative models and sequence conservation included as comparators in the PafA analysis (**Tables S20-S21**).

## Discussion

Accurate *de novo* prediction of catalytic residues remains challenging for enzymes with limited sequence or structural similarity to annotated proteins, because catalytic function is rarely encoded by isolated sequence positions alone. Instead, enzyme catalytic often emerges from local constellations of residues that are organised within three-dimensional active-site environments. Our results support this view: catalytic residues show pronounced spatial clustering, and enzymes preferentially resolved by SCOPE were characterised by compact catalytic-residue arrangements, physicochemical consistency with annotated catalytic sites and descriptive differences in functional-category frequencies. These findings indicate that catalytic-site prediction is better framed as recognition of local structural context than as independent residue classification. These findings support incorporating local structural context into catalytic-residue prediction.

SCOPE was designed around this principle by combining residue-level structural graphs with evolutionary and physicochemical information ^33^, together with Gaussian smoothing constraint that reinforces locally coherent signals among neighbouring residues. Ablation analyses showed that Gaussian smoothing was required for complete recognition of a subset of SCOPE complete-recovery enzymes and stabilised several annotated catalytic residues that otherwise fell below the prediction threshold. These Gaussian-dependent residues were frequently embedded within local catalytic neighbourhoods, suggesting that the Gaussian auxiliary objective can support the recognition of structurally relevant catalytic residues. Importantly, this effect remained detectable when AlphaFold-predicted structures were used as input, indicating that the contribution of the Gaussian auxiliary objective is not restricted to experimentally resolved structures. These results support its use as an auxiliary training constraint for catalytic-residue prediction from protein structural graphs.

The prospective mutagenesis experiments further support the functional relevance of SCOPE-predicted sites in experimentally tractable enzyme systems. Across four enzymes with distinct catalytic mechanisms, substitutions at the SCOPE-selected positions substantially reduced enzyme activity, supporting the functional importance of these sites. These effects may arise through changes in folding, stability, substrate recognition, cofactor coordination or subunit organization, and therefore do not by themselves establish a direct catalytic role for each tested residue. These findings support the utility of SCOPE for prioritising candidate residues for enzyme functional annotation and rational biocatalyst engineering. To further assess robustness to conformational variability, motivated by recent ensemble-based evaluations in protein-nucleic acid binding site prediction ^34^, we evaluated SCOPE using BioEmu-generated ensembles ^35^. Across 20 nonredundant benchmark enzymes, SCOPE achieved similar macro-AUPRC with experimental structures and ensemble-averaged predictions (0.7109 and 0.7118, respectively), with limited variation in residue probabilities across conformations (**Tables S9-S11**). These results support robustness to sampled backbone and contact-graph variability under fixed residue features. Nevertheless, SCOPE currently relies on static structural representations, and this evaluation does not fully address conformation-dependent atomic environments, ligand-induced rearrangements, transient catalytic states or reaction-specific chemical information. Future extensions that integrate these aspects may further improve the discovery of catalytic and functionally critical residues across diverse enzyme families.

## Methods

### Datasets

We utilised the enzyme catalytic residue dataset compiled from SCREEN, which contains 1,055 enzymes with experimentally determined catalytic residues. Catalytic-residue annotations and corresponding reaction-mechanism information were first collected from the M-CSA database, which provides experimentally validated catalytic sites and their positions in protein sequences. Subsequently, we further integrated the EF family dataset, which contains catalytic-site annotations for enzymes from diverse SCOP families, to increase dataset diversity. To reduce sequence redundancy and minimise overfitting, we clustered the combined dataset using CD-HIT at a 40% sequence-identity threshold and randomly retained one representative protein from each cluster ^36^. For each enzyme sequence, we retrieved the corresponding three-dimensional structure from the PDB database. Missing residues in processed structures were handled by global sequence alignment and gap filling to ensure consistency between sequence and structural data. To ensure reliable model evaluation, the dataset was split into training and validation sets at a 9:1 ratio. The 40% sequence-identity threshold was used to reduce sequence redundancy and was not interpreted as ensuring independence of homologous families or catalytic-site architectures. We therefore performed an additional evaluation using a CATH homologous-superfamily-disjoint partition of the development cohort, with models retrained on the resulting training set and selected using the corresponding validation set (**Table S25)**.

To comprehensively evaluate model performance, we further collected five widely used independent test datasets, comprising a total of 3,142 catalytic residues (**Table S7**). These datasets include the EF-fold, EF-superfamily, HA-superfamily, and PC and NN datasets, which cover enzymes across different SCOP folds, superfamilies, and enzyme classes ^37, 38, 39^. To assess residual train-test homology, we additionally evaluated the frozen SCOPE model on nested test subsets with maximum BLASTP sequence identity <40%, <30%, or <25% to all training and validation proteins, considering alignments covering at least 70% of both sequences. Identical sequences across test sets were counted once in the primary combined analysis (**Tables S12-S13**).We additionally evaluated frozen SCOPE and SCREEN predictions on a structurally restricted test subset **(Table S36)**.

For prospective experimental validation, four additional enzymes, including tyrosine ammonia-lyase (Fc-TAL2) ^40^, laccase (CotA) ^31^, diacetylchitobiose deacetylase (TGDac), and alkaline phosphatase (rsAP) ^32^, were used as an independent experimental validation set. To remain consistent with the 40% sequence-identity threshold used for CD-HIT deduplication of the training corpus, each experimental enzyme was screened against all retained training and validation sequences using BLASTP ^41^. Proteins were considered homologous only when an alignment showed ≥40% sequence identity with both query and subject alignment coverage ≥70%. Under this criterion, none of the four experimental enzymes shared a homologous sequence with any retained training or validation protein, supporting their use as an independent prospective validation set.

For external evaluation, we retained the official CataloDB test partition and constructed a fixed validation partition from official training sequence groups. SCOPE and SCREEN were retrained on the resulting split, with checkpoints selected using validation data only. Published Squidly and EasIFA weights were evaluated on matched available inputs, and a BLAST label-transfer baseline used only the corresponding SCOPE training records as references (**Supplemental Information, “CataloDB evaluation and a training restricted BLAST baseline”**).

### Residue-level Graph Representation of Enzymes

To model the three-dimensional organisation of enzyme structures, each enzyme was represented as a residue-level graph,, where denotes the set of residue nodes and denotes the set of edges connecting residue pairs. Each residue was represented by a 64-dimensional feature vector comprising PSSM profiles (20 dimensions), HMM profiles (30 dimensions), atomic structural descriptors (7 dimensions), and amino acid lookup features (7 dimensions). The graph and aligned residue features were used as inputs to the DeepGCN module. Graph construction and normalisation are described in Supplemental Information, “Graph Construction”, and feature extraction and alignment are described in “Multimodal residue feature construction”. Contact-threshold sensitivity results are provided in **Table S26**.

### ProtT5-based sequence representation

To capture contextual and evolutionary information from enzyme sequences, we used the pre-trained protein language model ProtT5 to extract residue-level sequence embeddings ^13^. ProtT5 is a Transformer-based protein language model from the ProtTrans framework that was trained in a self-supervised manner on large-scale protein sequence databases. In this study, the encoder component of ProtT5-XL-UniRef50 was used as the feature extractor, implemented using T5EncoderModel and T5Tokenizer from the Hugging Face framework. Enzyme sequences were pre-processed, tokenised using the ProtT5 tokenizer and passed through the encoder to obtain residue-level embedding features. Batching, padding removal and embedding storage are described in Supplemental Information, “ProtT5 embedding extraction and batching”.

### DeepGCN for enzyme structure representation

In graph convolutional networks, increasing network depth can cause residue representations to become progressively similar after repeated neighbourhood aggregation, leading to over-smoothing and reduced discriminative capacity. To mitigate this issue, the GCNII model introduces an initial residual connection and depth-dependent identity mapping, allowing deeper graph propagation while preserving informative node-level representations ^42^. Based on this principle, we constructed a DeepGCN module to learn residue-level structural representations from enzyme contact graphs for catalytic-residue prediction.

In SCOPE, the DeepGCN comprises three stacked graph convolutional layers. The residue input dimension is 64, and both the hidden and output dimensions were set to 512. Dropout was applied between layers to mitigate overfitting. This architecture produces residue-level representations that encode local structural neighbourhood information. In parallel, ProtT5-derived residue embeddings were projected through a linear layer followed by nonlinear activation to generate global sequence-derived representations. The local structural representations and global evolutionary representations were concatenated and integrated through an additional linear transformation and nonlinear activation. This resulting integrated feature jointly captures the three-dimensional spatial topology of enzyme residues and their evolutionary semantic context. The layer-wise update equations and adjacency normalisation are provided in Supplemental Information, “DeepGCN implementation and update equations”.

### Gaussian Smoothing

To incorporate the spatial clustering properties of catalytic residues into model training, SCOPE applies a Gaussian smoothing operation to residue-level representations generated by the DeepGCN module ^43^. This operation acts as an auxiliary regularisation constraint that encourages neighbouring residues to maintain locally coherent catalytic signals, thereby improving the stability of catalytic-residue recognition within active-site microenvironments.

The Gaussian smoothing operation was implemented as a fixed one-dimensional convolution along the residue dimension. The Gaussian kernel length was set to 9, and the standard deviation was denoted as *σ* = 2. For a kernel position (*i*), the corresponding Gaussian weight was calculated as:

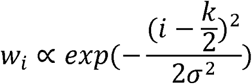

The kernel weights were so that the smoothing operation was equivalent to a weighted average over local residue neighbourhoods. During convolution, the kernel was applied to the fused residue features with a stride of 1 and padding of 4, ensuring that the output sequence length matched the input sequence length. Each residue representation was therefore updated by integrating information from itself and up to four neighbouring residues on either side, with the central residue receiving the highest weight and more distant residues receiving progressively lower weights. This design is consistent with the observation that catalytic residues often occur within locally organised structural environments. By encouraging local consistency in residue-level representations, Gaussian smoothing helps stabilise weak but spatially supported catalytic signals and reduces the likelihood that residues embedded within catalytic neighbourhoods are missed during prediction.

### Model Training and Evaluation

Model parameters were optimised using the Adam optimiser, which adaptively estimates the firstand second-order moments of the gradients to support stable and efficient training ^44^. We set ***β*_1_ = 0.9** and ***β*_2_=0.999**, and a learning rate of **5×10**^-5^ Residue-level catalytic site prediction was formulated as a binary classification problem, with cross-entropy loss used as the primary optimisation objective:

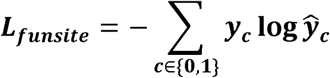

Here, ***y_c_*** denotes the true label, while ***ŷ_c_*** represents the model’s predicted probability for the category *c*. Concurrently, an auxiliary prediction based on Gaussian smoothing representation is introduced, with its corresponding loss defined as:

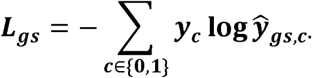

The composite loss function is expressed as:

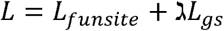

Here, λ=10^-4^ denotes the auxiliary task weight, introducing a smoothing constraint to promote stable representation learning. During each training iteration, backpropagation was performed on the composite loss, and model parameters were updated by the optimiser. The training loss value is recorded as the average of the primary task loss *L_funsite_*. To mitigate overfitting, dropout regularisation is applied with a dropout rate of 0.1.

The trained model was evaluated sequentially on five independent test datasets: NN, HA-superfamily, EF-superfamily, PC and EF-fold. Binary predictions were obtained using a fixed decision threshold of 0.5. Performance was assessed using accuracy, precision, recall, F1, AUROC, AUPRC and MCC. Metric definitions and equations are provided in Supplemental Information, “Evaluation metrics”.

We measured end-to-end runtime and peak CPU and GPU memory usage for three proteins (245-723 residues), starting from sequences and supplied experimental PDB structures. All input features were regenerated before prediction using frozen SCOPE weights. Measurements included feature generation, model initialization, inference and output writing, but excluded structure acquisition or prediction and one-time database and model installation. Hardware specifications, measurement procedures and per-protein results are provided in **Table S18**.

## Resource Availability

### Lead Contact

Further information and requests for resources should be directed to and will be fulfilled by the lead contact, Fuyi Li.

### Materials Availability

Plasmids and recombinant proteins generated in this study are available from the lead contact upon reasonable request and may require a completed materials transfer agreement.

### Data and Code Availability

The SCOPE source code, released model checkpoint and usage documentation are publicly available at https://github.com/ABILiLab/SCOPE. The reproducibility directory provides a Python 3.10 CPU environment specification, configurable feature-generation and prediction scripts, example inputs, and a Google Colab notebook for inference from prepared features.The original benchmark datasets and precomputed features are available at https://huggingface.co/datasets/Biocollab/SCREEN. Sources of additional public datasets and the evaluation protocols are described in the Methods and supplemental information.

## Acknowledgments

This work was supported by the National Key Research and Development Program of China [2022YFF1000100]; the National Natural Science Foundation of China [62202388]; and the Qin Chuangyuan Innovation and Entrepreneurship Talent Project [QCYRCXM-2022-230]. F.L. was supported by the Australian National Health and Medical Research Council (NHMRC) Investigator Fellowship [2041439]. The South Australian immunoGENomics Cancer Institute (SAiGENCI) received grant funding from the Australian Government.

## Author Contributions

F.L. conceptualised and supervised the project. Z.R. developed the model. T.P. processed the data. X.G. developed the graphical user interface. Z.R., T.P., Y.Z., Y.H., and H.S. contributed to writing the manuscript. J.W., X.S., Z.L., X.C., S.Z., and Z.W. performed the biological experiments. J.S., G.Z., and F.L. designed the experiments and revised the manuscript. All authors reviewed and approved the final manuscript.

## Declaration of generative AI and AI-assisted technologies in the manuscript preparation process

No generative AI or AI-assisted technologies were used in the preparation of this manuscript.

## References

1. Holliday, G. L., Fischer, J. D., Mitchell, J. B. O. & Thornton, J. M. Characterizing the complexity of enzymes on the basis of their mechanisms and structures with a bio-computational analysis. The FEBS Journal 278, 3835–3845 10.1111/j.1742-4658.2011.08190.x (2011).

2. Bornscheuer, U. T., et al. Engineering the third wave of biocatalysis. Nature 485, 185–194 10.1038/nature11117 (2012).

3. Ribeiro, A. J. M., et al. Mechanism and Catalytic Site Atlas (M-CSA): a database of enzyme reaction mechanisms and active sites. Nucleic Acids Research 46, D618–D623 10.1093/nar/gkx1012 (2018).

4. Markin, C. J., et al. Revealing enzyme functional architecture via high-throughput microfluidic enzyme kinetics. Science 373, 10.1126/science.abf8761 (2021).

5. Schnoes, A. M., Brown, S. D., Dodevski, I. & Babbitt, P. C. Annotation Error in Public Databases: Misannotation of Molecular Function in Enzyme Superfamilies. PLOS Computational Biology 5, e1000605 10.1371/journal.pcbi.1000605 (2009).

6. Hackett, R. E., et al. Investigating enzyme function by geometric matching of catalytic motifs. Protein Science 35, e70744 10.1002/pro.70744 (2026).

7. Sun, J., Wang, J., Xiong, D., Hu, J. & Liu, R. CRHunter: integrating multifaceted information to predict catalytic residues in enzymes. Scientific Reports 6, 34044 10.1038/srep34044 (2016).

8. Zhang, T., et al. Accurate sequence-based prediction of catalytic residues. Bioinformatics 24, 2329–2338 10.1093/bioinformatics/btn433 (2008).

9. Song, J., et al. PREvaIL, an integrative approach for inferring catalytic residues using sequence, structural, and network features in a machine-learning framework. Journal of Theoretical Biology 443, 125–137 10.1016/j.jtbi.2018.01.023 (2018).

10. Guo, X., et al. Language Model-Enhanced Geometric Learning for Metalloprotein Pathogenic Mutations. IEEE J Biomed Health Inform Pp, 10.1109/jbhi.2026.3713319 (2026).

11. Song, Y., et al. Accurately predicting enzyme functions through geometric graph learning on ESMFold-predicted structures. Nat Commun 15, 8180 10.1038/s41467-024-52533-w (2024).

12. Ran, Z., et al. A scalable equivariant graph network framework for precise protein function prediction. Genome Biol 26, 407 10.1186/s13059-025-03886-y (2025).

13. Elnaggar, A., et al. ProtTrans: Toward Understanding the Language of Life Through Self-Supervised Learning. IEEE Trans Pattern Anal Mach Intell 44, 7112–7127 10.1109/tpami.2021.3095381 (2022).

14. Remmert, M., Biegert, A., Hauser, A. & Söding, J. HHblits: lightning-fast iterative protein sequence searching by HMM-HMM alignment. Nature Methods 9, 173–175 10.1038/nmeth.1818 (2012).

15. Berman, H. M. The Protein Data Bank. Nucleic Acids Research 28, 235–242 (2000).

16. Kipf, T. N. & Welling, M.Semi-Supervised Classification with Graph Convolutional Networks.In:International Conference on Learning Representations(2017).

17. Hamilton, W. L., Ying, R. & Leskovec, J. Inductive Representation Learning on Large Graphs. (2017).

18. Li, G., Müller, M., Thabet, A. & Ghanem, B. DeepGCNs: Can GCNs Go As Deep As CNNs? IEEE, (2020).

19. Bartlett, G. J., Porter, C. T., Borkakoti, N. & Thornton, J. M. Analysis of catalytic residues in enzyme active sites. Journal of molecular biology 324(1), 105–121 (2002).

20. Li, Q., Han, Z. & Wu, X. M. Deeper Insights into Graph Convolutional Networks for Semi-Supervised Learning. (2018).

21. Jeffery, C. J., Bahnson, B. J., Chien, W., Ringe, D. & Petsko, G. A. Crystal structure of rabbit phosphoglucose isomerase, a glycolytic enzyme that moonlights as neuroleukin, autocrine motility factor, and differentiation mediator. Biochemistry 39, 955–964 10.1021/bi991604m (2000).

22. Davies, G. J., Brzozowski, A. M., Dauter, M., Varrot, A. & Schülein, M. Structure and function of Humicola insolens family 6 cellulases: structure of the endoglucanase, Cel6B, at 1.6 A resolution. Biochem J 348 Pt 1, 201–207 (2000).

23. Mattevi, A., et al. Crystal structures and inhibitor binding in the octameric flavoenzyme vanillyl-alcohol oxidase: the shape of the active-site cavity controls substrate specificity. Structure 5, 907–920 10.1016/s0969-2126(97)00245-1 (1997).

24. Pan, T., et al. SCREEN: A Graph-based Contrastive Learning Tool to Infer Catalytic Residues and Assess Enzyme Mutations. Genomics, Proteomics & Bioinformatics 22, 10.1093/gpbjnl/qzae094 (2024).

25. Shen, X., et al. A Highly Sensitive Model Based on Graph Neural Networks for Enzyme Key Catalytic Residue Prediction. J Chem Inf Model 63, 4277–4290 10.1021/acs.jcim.3c00273 (2023).

26. Rieger, W. J. F., Bodén, M., Arnold, F. & Mora, A. Squidly harnesses enzyme functional hierarchy and contrastive learning to efficiently predict catalytic residues from sequence. Elife 14, 10.7554/eLife.108186 (2026).

27. Wang, X., et al. Multi-modal deep learning enables efficient and accurate annotation of enzymatic active sites. Nature Communications 15, 7348 10.1038/s41467-024-51511-6 (2024).

28. Ying, R., Bourgeois, D., You, J., Zitnik, M. & Leskovec, J. GNNExplainer: Generating Explanations for Graph Neural Networks. Advances in neural information processing systems 32, 9240–9251 (2019).

29. Jumper, J., et al. Highly accurate protein structure prediction with AlphaFold. Nature 596, 583–589 10.1038/s41586-021-03819-2 (2021).

30. Huang, Y., Jiang, X., Chen, W., Zhang, G. & Wang, Q. Characterization of highly active tyrosine ammonia lyase and its application in biosynthesis of p-coumaric acid. Chinese Journal of Biotechnology 38, 4553–4566 10.13345/j.cjb.220182 (2022).

31. Sun, X., et al. Engineering Bacterial Laccase with Improved Catalytic Activity and Thermostability by Rational Design. Applied Biochemistry and Biotechnology 197, 4685–4701 10.1007/s12010-025-05240-1 (2025).

32. Xie, Y., et al. A highly efficient alkaline phosphatase for dephosphorylation of macromolecular substrates. Int J Biol Macromol 352, 151134 10.1016/j.ijbiomac.2026.151134 (2026).

33. Guo, X., Ran, Z. & Li, F. Kinase–inhibitor binding affinity prediction with pretrained graph encoder and language model. Briefings in Bioinformatics 26, 10.1093/bib/bbaf338 (2025).

34. Li, P., Liu, Y., Liang, L. & Liu, R. DyProL: Dynamic Ensemble Representation Learning for Protein-Nucleic Acid Binding Site Prediction. Adv Sci (Weinh), e77501 10.1002/advs.77501 (2026).

35. Lewis, S., et al. Scalable emulation of protein equilibrium ensembles with generative deep learning. Science 389, eadv9817 10.1126/science.adv9817 (2025).

36. Li, W. & Godzik, A. Cd-hit: a fast program for clustering and comparing large sets of protein or nucleotide sequences. Bioinformatics 22, 1658–1659 10.1093/bioinformatics/btl158 (2006).

37. Gutteridge, A., Bartlett, G. J. & Thornton, J. M. Using a neural network and spatial clustering to predict the location of active sites in enzymes. J Mol Biol 330, 719–734 10.1016/s0022-2836(03)00515-1 (2003).

38. Petrova, N. V. & Wu, C. H. Prediction of catalytic residues using Support Vector Machine with selected protein sequence and structural properties. BMC Bioinformatics 7, 312 10.1186/1471-2105-7-312 (2006).

39. Chea, E. & Livesay, D. R. How accurate and statistically robust are catalytic site predictions based on closeness centrality? BMC Bioinformatics 8, 153 10.1186/1471-2105-8-153 (2007).

40. Wang, J., Zhang, N., Huang, Y., Li, S. & Zhang, G. Simple and efficient enzymatic procedure for p-coumaric acid synthesis: Complete bioconversion and biocatalyst recycling under alkaline condition. Biochemical Engineering Journal 188, 108693 10.1016/j.bej.2022.108693 (2022).

41. Camacho, C., et al. BLAST+: architecture and applications. BMC Bioinformatics 10, 421 10.1186/1471-2105-10-421 (2009).

42. Chen, M., Wei, Z., Huang, Z., Ding, B. & Li, Y. Simple and Deep Graph Convolutional Networks. In: 37th International Conference on Machine Learning: ICML 2020, Online, 13-18 July 2020, Part 3 of 15) (2021).

43. Yamashita, Y. & Wakahara, T. Stabilized Calculation of Gaussian Smoothing and Its Differentials Using Attenuated Sliding Fourier Transform. In: 2020 25th International Conference on Pattern Recognition (ICPR)) (2021).

44. Kingma, D. & Ba, J. Adam: A Method for Stochastic Optimization. Computer Science, (2014).

