## Supplementary Information for "SCOPE: Gaussian-regularized graph learning for enzyme catalytic residue prediction"

**Supplemental Tables**

**Table S1.** Summary of the reviewed predictors for enzyme catalytic site prediction.

| **Tool** | **Year** | **Feature encoding scheme** | **Algorithm** | **Evaluation metrics** |
| --- | --- | --- | --- | --- |
| CRpred | 2008 | PSI-BLAST, AveCH, PSSM | SVM | Precision, TP |
| CRHunter | 2016 | PSSM, ${MEscore}_{DT}$ | SVM | Precision, $F_{1}$, Recall, AUC, ACC, MCC |
| PREvaIL | 2018 | DSSP, PSSM, HSE | RF | TP, PRE |
| AEGAN | 2023 | AASM, PSSM, | GAT | Precision, $F_{1}$, Recall, AUC, AUPR |
| SCREEN | 2024 | HMM, PSSM, ProtT5 | Contrastive learning, GCN | AUPR, Precision, $F_{1}$, Recall, AUC, ACC, MCC |

1. Feature encoding scheme: HMM—hidden markov model; PSSM—positional specific scoring matrix; HSE—half sphere exposure; AveCH—average cumulative hydrophobicity; AASM—amino acid spatial model.

2. Algorithm: GCN—graph convolutional network; SVM—support vector machine; RF—random forest; GAT—graph attention network.

3. Evaluation metrics: AUC—Area under the curve; AUPR—area under the precision-recall curve; MCC—Matthews correlation coefficient; TP—true positive.

**Table S2.** Ablation study of the Gaussian smoothing module in SCOPE.

| **Model Variant** | **Test Dataset** | **Precision** | **F1** | **AUC** | **AUPRC** | **MCC** |
| --- | --- | --- | --- | --- | --- | --- |
| SCOPE | NN | 0.7096 | 0.7523 | 0.9902 | 0.6683 | 0.7510 |
|  | HA_superfamily | 0.6934 | 0.7133 | 0.9804 | 0.6388 | 0.7110 |
|  | EF_superfamily | 0.6452 | 0.6283 | 0.9710 | 0.5767 | 0.6244 |
|  | PC | 0.7143 | 0.7731 | 0.9903 | 0.6730 | 0.7728 |
|  | EF_fold | 0.6207 | 0.6506 | 0.9754 | 0.6300 | 0.6472 |
| SCOPE w/o Gaussian smoothing | NN | 0.6787 | 0.7370 | 0.9900 | 0.6633 | 0.7368 |
|  | HA_superfamily | 0.7044 | 0.7105 | 0.9803 | 0.6367 | 0.7080 |
|  | EF_superfamily | 0.5625 | 0.6193 | 0.9686 | 0.5724 | 0.6177 |
|  | PC | 0.7075 | 0.7576 | 0.9901 | 0.6664 | 0.7564 |
|  | EF_fold | 0.5978 | 0.6350 | 0.9697 | 0.6097 | 0.6318 |

**Table S3.** Complete-recovery records associated with SCOPE, SCREEN and retrained AEGAN across benchmark test sets.

| **Test set** | **SCOPE** | **SCREEN** | **Retrained AEGAN** |
| --- | --- | --- | --- |
| NN | 28 | 28 | 0 |
| HA-superfamily | 62 | 66 | 0 |
| EF-superfamily | 10 | 9 | 0 |
| PC | 14 | 13 | 0 |
| EF-fold | 5 | 5 | 0 |
| Total dataset-level records | 119 | 121 | 0 |

**Table S4.** Predicted catalytic residues by SCOPE

| **Enzyme name** | **Experimentally validated catalytic residues** | **SCOPE-predicted functionally important residues** | **SCOPE prediction score** |
| --- | --- | --- | --- |
| Fc-TAL2 | Y50 | Y50 | 0.6341 |
|  | A139 | S140 | 0.5417 |
|  | S140 | R291 | 0.6404 |
|  | G141 | F337 | 0.5599 |
|  | Y287 | N420 | 0.5824 |
|  |  | D489 | 0.5344 |
| CotA | H105 | H105 | 0.5117 |
|  | H153 | H107 | 0.5956 |
|  | H419 | H155 | 0.6442 |
|  | H422 | H419 | 0.6642 |
|  | H424 | H422 | 0.5892 |
|  | H491 | H424 | 0.7038 |
|  | C492 | H491 | 0.6575 |
|  | H493 | C492 | 0.6862 |
|  | H497 | H493 | 0.6392 |
|  | M502 | H497 | 0.5358 |
| TGDac | H36 | H36 | 0.6224 |
|  | D39 | D38 | 0.7195 |
|  | H147 | D39 | 0.7668 |
|  |  | D107 | 0.6209 |
|  |  | H144 | 0.7758 |
|  |  | D146 | 0.5316 |
|  |  | H147 | 0.7714 |
| rsAP | S123 | R187 | 0.6570 |
|  | R187 | H484 | 0.6565 |
|  | H374 |  |  |

**Table S5.** Experimental enzyme activity measurements for wild-type enzymes and alanine-substitution mutants.

| **Enzyme** | **Mutant** | **Activity R1** | **Activity R2** | **Activity R3** | **Activity mean** | **Activity SD** |
| --- | --- | --- | --- | --- | --- | --- |
| Fc-TAL2 | WT | 2.49 | 2.19 | 2.57 | 2.41 | 0.20 |
|  | R291A | 0.08 | 0.09 | 0.08 | 0.08 | 0.01 |
|  | F337A | 0.02 | 0.01 | 0.02 | 0.02 | 0.00 |
|  | D489A | 0.01 | 0.03 | 0.04 | 0.03 | 0.01 |
|  | N420A | 0.02 | 0.03 | 0.03 | 0.03 | 0.00 |
| CotA | WT | 1.75 | 1.77 | 1.78 | 1.77 | 0.01 |
|  | H107A | 0.03 | 0.04 | 0.02 | 0.03 | 0.01 |
|  | H155A | 0.01 | 0.01 | 0.01 | 0.01 | 0.00 |
| TGDac | WT | 96.97 | 99.72 | 110.97 | 102.55 | 7.42 |
|  | D38A | 0.11 | 0.22 | 1.16 | 0.50 | 0.57 |
|  | D107A | 0.00 | 0.00 | 0.00 | 0.00 | 0.00 |
|  | H144A | 2.91 | 2.40 | 3.07 | 2.79 | 0.35 |
|  | D146A | 3.64 | 3.40 | 5.51 | 4.18 | 1.15 |
| rsAP | WT | 1.57 | 2.49 | 2.76 | 2.27 | 0.62 |
|  | H484A | 0.00 | 0.00 | 0.00 | 0.00 | 0.00 |

**Table S6.** Reference-annotated catalytic residues and predictions from SCOPE, its no-Gaussian variant and SCREEN in four prospective enzymes

| **Enzyme name** | **Reference-annotated catalytic residues** | **SCOPE predictions** | **w/o Gaussian predictions** | **SCREEN predictions** |
| --- | --- | --- | --- | --- |
| Fc-TAL2 | Y50 | Y50 | Y50 | Y50 |
|  | A139 | S140 | R291 | N192 |
|  | S140 | R291 | F337 |  |
|  | G141 | F337 | H338 |  |
|  | Y287 | N420 |  |  |
|  |  | D489 |  |  |
| CotA | H105 | H105 | H105 | H105 |
|  | H153 | H107 | H107 | H107 |
|  | H419 | H155 | H153 | H153 |
|  | H422 | H419 | H155 | H155 |
|  | H424 | H422 | H419 | M157 |
|  | H491 | H424 | H422 | T160 |
|  | C492 | H491 | H424 | H419 |
|  | H493 | C492 | H491 | H424 |
|  | H497 | H493 | C492 | H491 |
|  | M502 | H497 | H493 | C492 |
|  |  |  |  | H493 |
|  |  |  |  | H497 |
| TGDac | H36 | H36 | H36 | H36 |
|  | D39 | D38 | D38 | D38 |
|  | H147 | D39 | D39 | D39 |
|  |  | D107 | Y141 | D107 |
|  |  | H144 | H144 | H144 |
|  |  | D146 | D146 | H147 |
|  |  | H147 | H147 |  |
| rsAP | S123 | R187 | S123 | S123 |
|  | R187 | H484 | R187 | D330 |
|  | H374 |  | H374 | H374 |
|  |  |  |  | H484 |

**Table S7.** Training, validation and test datasets

| **Type** | **Dataset name** | **Enzymes** | **Catalytic residues** | **Non-catalytic residues** |
| --- | --- | --- | --- | --- |
| Training and validation | Curated M-CSA | 762 | 3344 | 272287 |
|  | EF family | 293 | 1018 | 86013 |
| Test | EF fold | 189 | 605 | 48223 |
|  | EF superfamily | 236 | 716 | 62677 |
|  | HA superfamily | 282 | 933 | 97976 |
|  | NN | 163 | 599 | 56309 |
|  | PC | 81 | 289 | 24493 |

**Table S8.** The primers for mutants

| **Enzyme name** | **The primers name** | **Sequence (5’-3’)** |
| --- | --- | --- |
| Fc-TAL2 | R291A-F | TTACAGCCTGGCTT**GCG**TTCCGCAGATCC |
|  | R291A-R | GCGGAA**CGC**AAGCCAGGCTGTAATATTCCTG |
|  | F337A-F | CGGTGGTAAC**GCC**CACGGCGATTACATC |
|  | F337A-R | AATCGCCGTG**GGC**GTTACCACCGTGGTAAAC |
|  | N420A-F | CTATCCCGAAC**GCC**AACGACAACCAGGAC |
|  | N420A-R | GGTTGTCGTTGGCGTT**CGG**GATAGAATGAAC |
|  | D489A-F | GTTCAAAGAT**GCT**ATGATCATGTATCCGTAC |
|  | D489A-R | ACATGATCAT**AGC**ATCTTTGAACGGCGGAAC |
| CotA | H107A-F | TTCATTTAGCCGGCG**GCG**TCACGCCAGATG |
|  | H107A-R | GTGACGC**CGC**CGGCTAAATGAACAACAGTCTTTA |
|  | H155A-F | ATCACGATGCCGCCATG**GCG**CTCACCAGGC |
|  | H155A-R | AG**CGC**CATGGCGGCATCGTGATACCACAATATAG |
| TGDac | D38A-F | CATCCG**GCG**GATTGCGTTATTGGC |
|  | D38A-R | CAATC**CGC**CGGATGCGGTTC |
|  | D107A-F | ATCGT**GCG**ACCGAACTGCCG |
|  | D107A-R | CGGT**CGC**ACGATAATCCAGCC |
|  | H144A-F | GAAGCA**GCG**CCGGATCATGTGACC |
|  | H144A-R | CCGG**CGC**TGCTTCATAAGGCAGCC |
|  | D146A-F | CATCCG**GCG**CATGTGACCGCCGG |
|  | D146A-R | CACATG**CGC**CGGATGTGCTTCATAAGGC |
| rsAP | H484A-F | GTGCCTCT**GCA**GTGGATGTGAATATTTATGCA |
|  | H484A-R | ATCCAC**TGC**AGAGGCACCGTGGGTGGTCCAACC |

The mutation site is bolded and underlined in the primer sequence.

**Supplemental Methods and Results**

**Catalytic residues exhibit spatial clustering in enzyme structures**

To obtain a structural biology-informed understanding of the spatial organisation of catalytic residues, we further analysed the spatial distribution of experimentally annotated catalytic residues in three-dimensional enzyme structures. We calculated the Euclidean distances between catalytic residues and compared them with the corresponding distance distributions for non-catalytic residues ^[1, 2]^ (**Fig. S1a**). The median distance from a true catalytic residue to its nearest catalytic neighbour is approximately 6.55$Å$, whereas the median distance from a non-catalytic residue to its nearest catalytic residue is approximately 17.37$Å$, indicating that true catalytic residues are significantly clustered in spatial regions. This spatial clustering provides a structural rationale for representing local catalytic neighbourhoods in the residue graph, but does not establish an optimal distance threshold. The sensitivity of predictive performance to residue-contact thresholds of 4, 6 and 8 Å is evaluated separately in **Table S26**. Based on this observation, we further compared the consistency of spatial distribution between predicted and experimentally annotated catalytic residues. Residues predicted as catalytic showed distance distributions to the nearest true catalytic residues that closely resembled those of annotated catalytic residues (**Fig. S1b-c**). These results indicate that the SCOPE not only distinguishes catalytic from non-catalytic residues but also positions the predicted residues reasonably within structurally plausible regions surrounding true active sites.

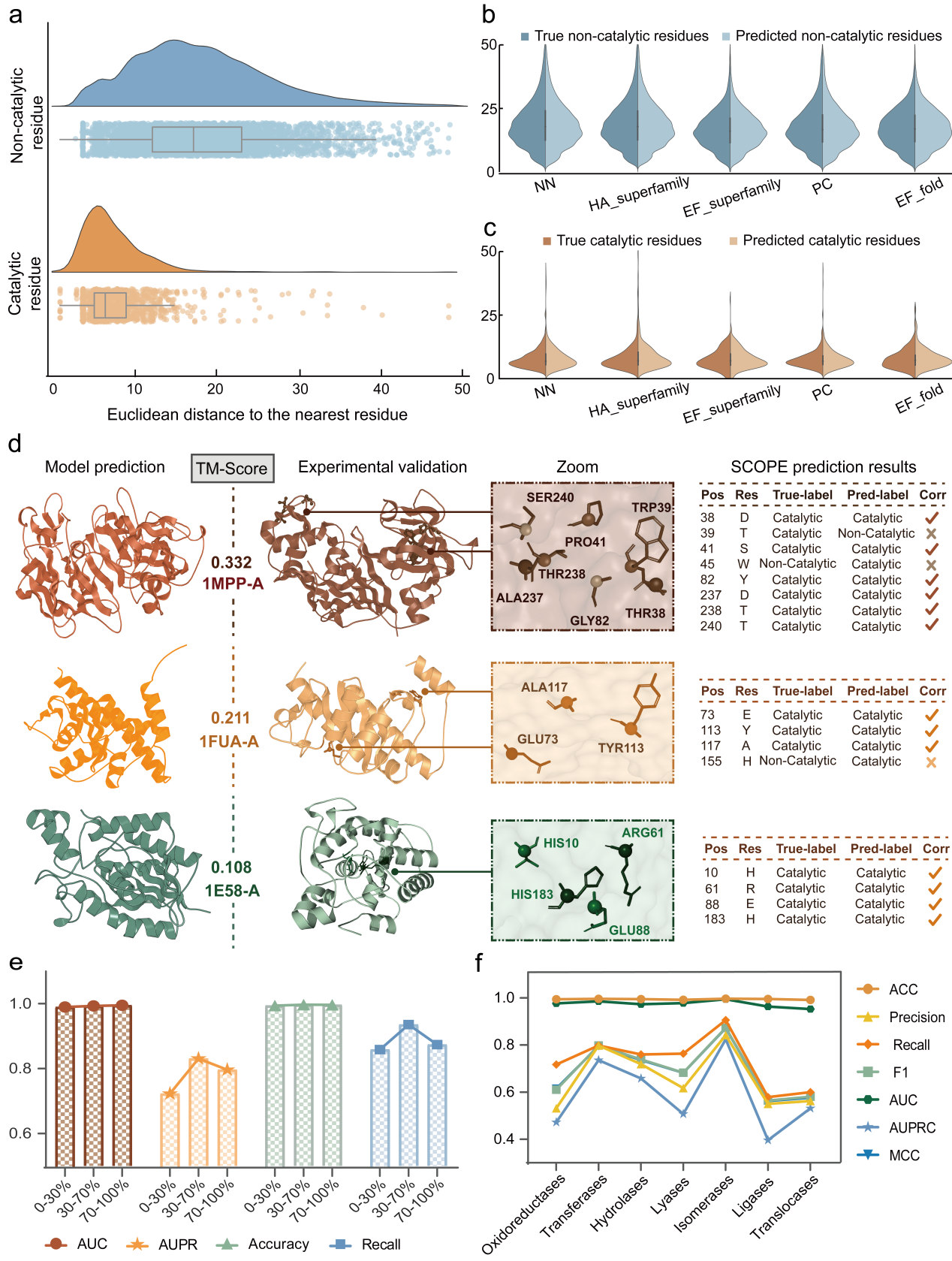

**Fig. S1 Spatial clustering of catalytic residues and structural localisation of SCOPE predictions. (a)** Raincloud plots showing the distributions of Euclidean distances from residues to their nearest catalytic residue. The upper panel corresponds to true non-catalytic residues, and the lower panel to true catalytic residues. **(b-c)** Violin plots showing the Euclidean distances ($Å$) to the nearest true catalytic residue across five test datasets. (b) True and predicted non-catalytic residues; (c) true and predicted catalytic residues.

**Structural and biochemical characterisation of predicted catalytic residues**

We then examined whether the residues predicted by SCOPE exhibit physicochemical properties consistent with experimentally annotated catalytic residues. We first analysed amino acid composition. As shown in **Fig. S2a**, the distribution of SCOPE-predicted catalytic residues across the 20 amino acids closely matched that of actual catalytic residues. For example, Asp (D) accounts for 17.3% of predicted catalytic residues and 16.1% of true catalytic residues; His (H) accounts for 20.0% and 17.2%, respectively; and Glu (E) accounts for 10.3% and 9.9%. Other residues frequently involved in catalysis, including Cys, Ser and Tyr, showed similarly consistent distributions. In contrast, predicted non-catalytic residues almost completely overlapped with true non-catalytic residues in their amino acid composition (**Fig. S2b**), indicating SCOPE robustly distinguishes catalytic from non-catalytic residue classes. We next quantified amino acid enrichment using the log-odds ratio (LOR) ^[3]^. As shown in **Fig. S2c**, several typical catalytic residues were significantly enriched in the prediction results, including His (H, +3.02), Cys (C, +2.57), Asp (D, +1.58), Arg (R, +1.08), Glu (E, +0.67), Tyr (Y, +0.64), and Lys (K, +0.51). These residues typically contain polar or charged side chains and frequently participate in proton transfer, nucleophilic attack or the stabilisation of transition states during catalytic reactions. In contrast, hydrophobic residues such as Val (V), Leu (L) and Ile (I) exhibit markedly negative LOR values, consistent with their tendency to reside in the hydrophobic protein core rather than directly participate in catalysis.

Analysis of residue charge (**Fig. S2d**) further reveals that approximately two-thirds of the catalytic residues are charged, whereas only about one-quarter of the non-catalytic residues carry charges. Specifically, among predicted catalytic residues, 38.6% are positively charged, 27.6% are negatively charged, and 33.8% are neutral, closely matching the corresponding proportions of 38.4%, 26.0%, and 35.7% observed in the true catalytic residues. We further analysed the local physicochemical environment of the predicted catalytic residues across multiple scales. At the single-residue level, Kyte-Doolittle hydrophobicity analysis (**Fig. S2e**) shows that catalytic residues are significantly more hydrophilic than non-catalytic residues (median KD$\text{≈}\text{-}$3.5 versus $\text{-}$0.7) ^[4]^. At the local sequence-neighbourhood scale, window-based analysis centred on catalytic residues (**Fig. S2f**) demonstrates that they were more enriched around catalytic residues than around non-catalytic residues, with the largest differences observed for window lengths of 3-7. Consistently, solvent accessibility-weighted hydrophobicity analysis (**Fig. S2g**) further indicates that catalytic regions are more hydrophilic, whereas non-catalytic residues tend to occupy more hydrophobic environments.

We then investigated the structural environments of predicted catalytic residues, including their backbone flexibility, secondary structure, and solvent accessibility. **Figs. S2h-i** show that predicted catalytic residues preferentially reside in regions with lower B-factors, indicating relatively rigid and stable structural environments, consistent with the general observation that active sites are often embedded in conformationally stable regions. DSSP-based secondary structure analysis (**Fig. S2j**) indicates that catalytic residues are predominantly located in loop regions. As shown in **Figs. S2k-l**, solvent accessibility and pocket analyses further reveal that catalytic sites are enriched in charged residues and form high-charge-density pockets. These structural features closely mirrored those of experimentally annotated catalytic residues, indicating that SCOPE not only localises catalytic sites accurately but also captures their native structural context.

To visualise these biochemical and structural features, we selected a representative transferase, 1B02 (Chain A), for which all catalytic residues were correctly predicted ^[5]^. As shown in **Fig. S2m**, the predicted catalytic residues formed a clear spatial cluster rather than a random distribution and were primarily located within a high-charge-density pocket composed of positively charged residues (Lys, Arg, His) and negatively charged residues (Asp, Glu). At the secondary-structure level, these catalytic residues are primarily located in loop regions and are supported by surrounding α-helices and β-sheets, forming a stable catalytic pocket architecture. At the local interaction level, catalytic residues and charged residues within their 4$Å$ neighbourhood constitute potential hydrogen-bond and salt-bridge interaction networks. Overall, these analyses indicate that SCOPE predicts catalytic residues with biochemical and structural properties that closely resemble those of native catalytic sites.

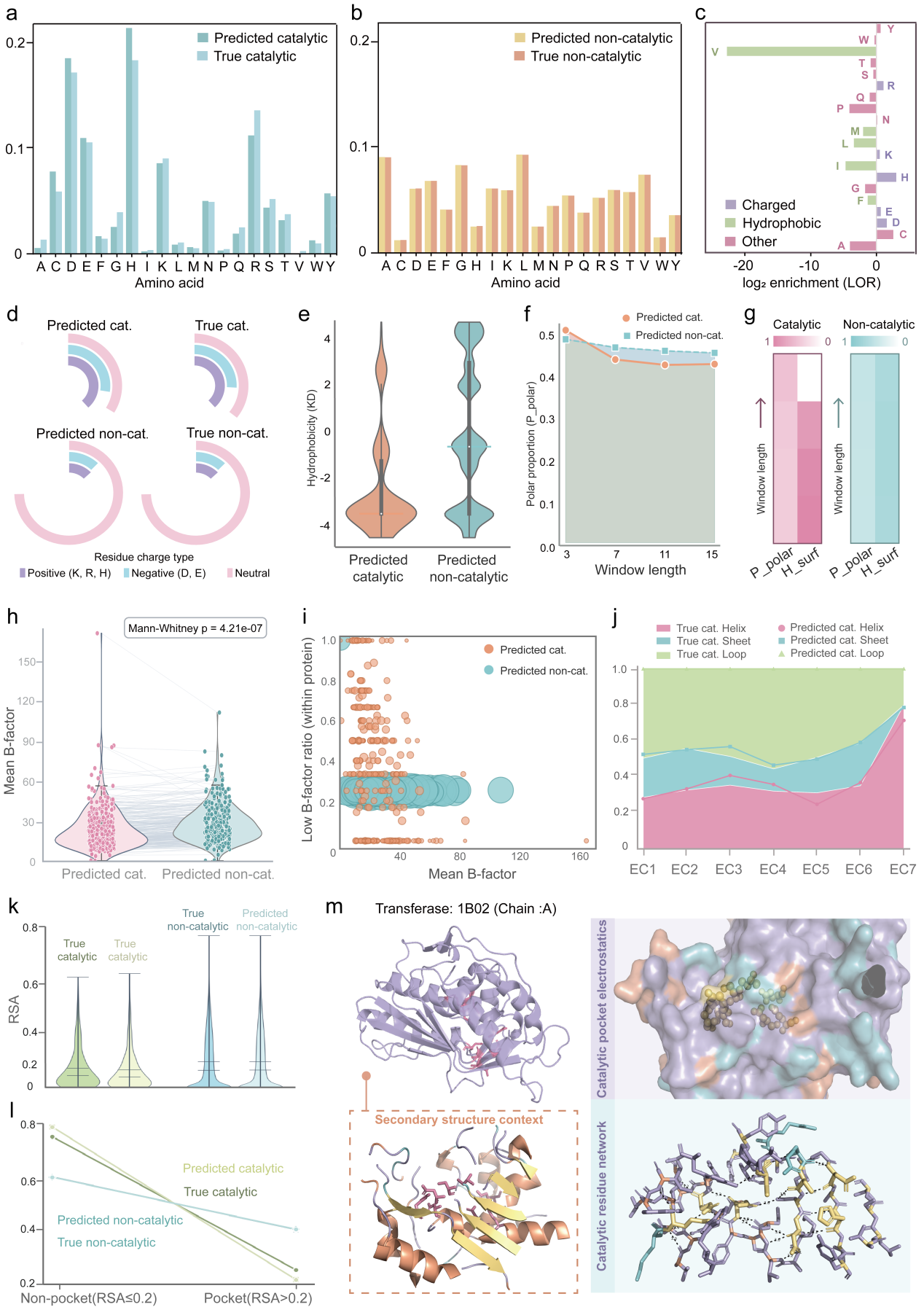

**Fig. S2 Compositional and structural characteristics** **of predicted catalytic residues. (a)** Amino acid distributions of predicted catalytic residues and experimentally annotated catalytic residues across the 20 standard amino acids. **(b)** Amino acid distributions of predicted non-catalytic residues and experimentally annotated non-catalytic residues. **(c)** Log-odds ratio (LOR) showing amino acid enrichment in predicted catalytic residues relative to true non-catalytic residues. Positive LOR values indicate enrichment, and negative values indicate depletion. Purple denotes amino acids enriched in predicted catalytic residues; green denotes amino acids depleted in predicted catalytic residues. **(d)** Distribution of residue charge classes. Purple indicates positively charged residues, green indicates negatively charged residues, and pink indicates neutral residues. **(e)** Violin plot of Kyte–Doolittle hydrophobicity indices for predicted catalytic and predicted non-catalytic residues. **(f)** Proportion of polar residues within window lengths of 3, 7, 11, and 15 centred on surface-exposed loop/coil residues (RSA > 0.2). **(g)** Heatmaps of the polar proportion and surface-weighted hydrophobicity under different window lengths. **(h)** Comparison of average B-factors between predicted catalytic and predicted non-catalytic residues. **(i)** Density distribution of B-factors. Circle size represents the number of residues within each B-factor interval. **(j)** Secondary structure distribution of predicted catalytic and true catalytic residues across seven enzyme classes. **(k)** Comparison of relative solvent accessibility (RSA). **(l)** Pocket analysis of predicted catalytic residues. **(m)** Structural visualisation of a transferase. Catalytic residues are shown as pink sticks. The upper-right panel shows the electrostatic environment of the catalytic pocket, with positively charged residues in blue, negatively charged residues in orange and catalytic residues highlighted as yellow spheres. The lower-left panel shows the secondary-structure context, with α-helices in orange, β-sheets in yellow, and loops in green. The lower-right panel shows the local interaction network, with dashed lines indicating potential hydrogen bonds or salt bridges.

**Diverse geometric organisation of catalytic residue clusters across structurally diverse enzymes**

To characterise the macroscopic geometry of catalytic clusters, we performed three-dimensional spatial analysis and geometric classification of predicted catalytic residue assemblies across diverse enzyme classes. Based on the three-dimensional distribution of residues, we classified catalytic residue clusters into four representative geometric types: Compact, Planar, Encircling, and Extended. These geometric categories were defined by the spatial distribution of residues within each cluster, considering both cluster size and the maximum pairwise residue distance in three-dimensional space. As shown in **Fig. S3a**, cluster radius varied systematically with geometric type: Compact clusters were generally small and tightly packed, whereas Extended clusters occupied a broader spatial range. Consistently, different geometric types differ significantly in both maximum cluster distance and cluster size (**Fig. S3b**), reflecting distinct modes of three-dimensional residue organisation. Enrichment analysis across EC classes further revealed functional preferences in cluster geometry (**Fig. S3c**). For example, Encircling clusters are broadly present across multiple EC classes, whereas Compact and Extended clusters tend to be associated with specific enzyme categories.

To illustrate how the same geometric type can arise in functionally distinct enzymes, we selected several examples of Encircling clusters for visualisation and functional analysis (**Fig. S3d**). In Lactate dehydrogenase A chain (PDB: 1DII, Chain A), an oxidoreductase that catalyses the reversible conversion of lactate and pyruvate in glycolysis, the predicted cluster overlaps with key residues including 473Y and 474R ^[6]^. In Hormone-sensitive lipase fragment (PDB: 1EI5, Chain A), a hydrolase involved in lipid metabolism, the predicted cluster captures multiple residues of the canonical Ser-His-Asp catalytic triad and adopts an encircling spatial arrangement that supports the binding of long-chain lipid substrates ^[7]^. In Fumarate hydratase class II (PDB: 1B93, Chain B), a lyase in the tricarboxylic acid cycle, the predicted cluster covers most core catalytic residues and forms a broad spatial arrangement that accommodates the catalytic conversion ^[8]^. This indicates that clusters with similar geometric shapes can support distinct substrates and reaction chemistries.

To evaluate SCOPE’s ability to identify catalytic clusters across divergent structural contexts, we analysed enzyme families that share catalytic functions but exhibit substantial fold variability. In the EC 3.5.1 amidohydrolase family (**Fig. S3e**), SCOPE captured distinct geometric phenotypes, ranging from encircling clusters in urease (PDB: 1KRA) ^[9]^ to Extended configurations in structurally divergent members. Similarly, in the EC 2.1.1 methyltransferase family, catalytic clusters ranged from encircling to Planar arrangements, with thymidylate synthase A (PDB: 1B02) ^[10]^ adopting an approximately coplanar geometry (**Fig. S3f**). These observations show that even for enzymes with the same catalytic function, differences in three-dimensional structures can lead to variations in the spatial geometry of catalytic residue clusters. These observations illustrate the diversity of predicted catalytic-cluster geometries across the examined enzymes.

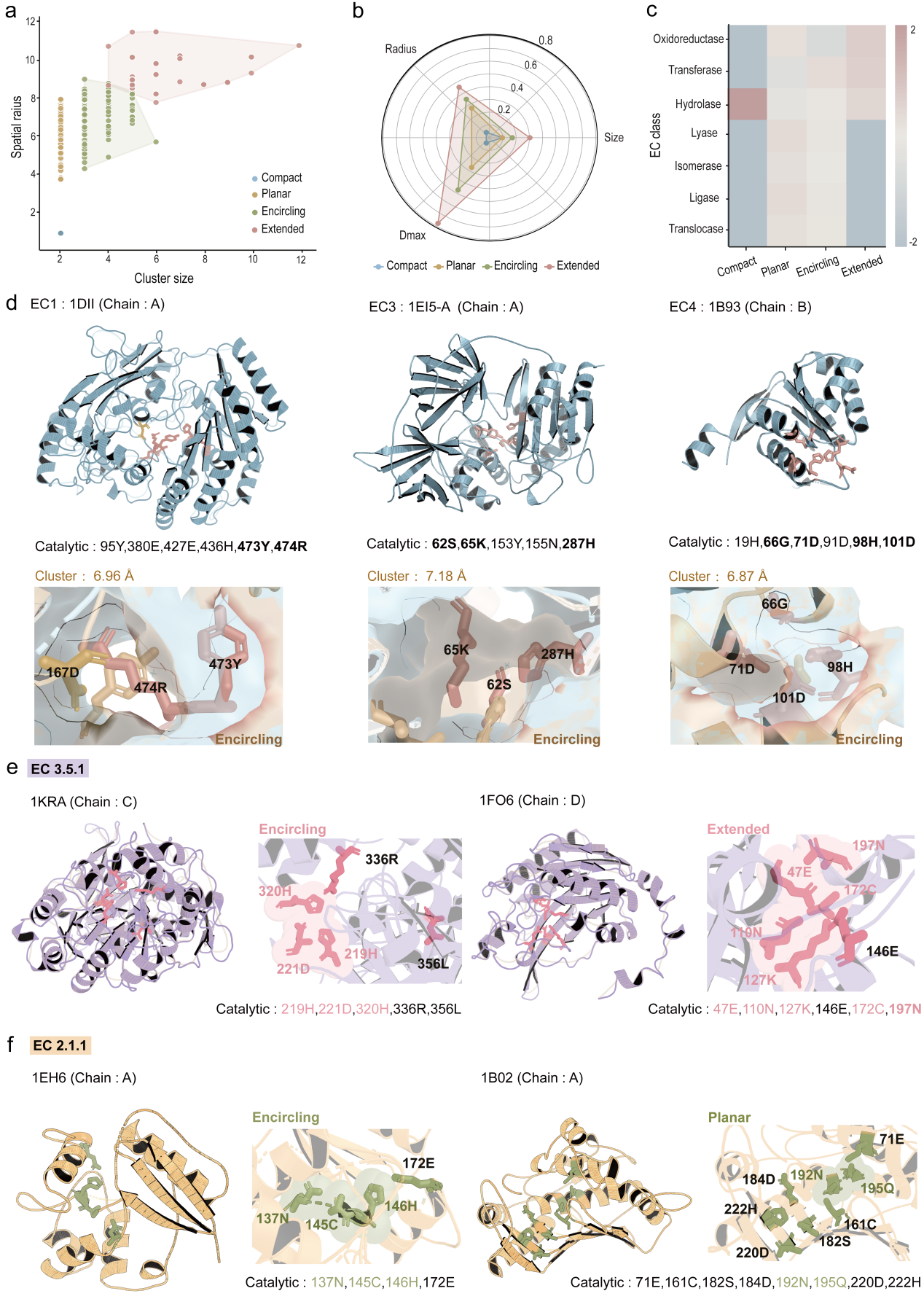

**Fig. S3 Geometric organization of catalytic residue clusters across enzymes. (a)** Scatter plot of cluster size versus spatial radius for all catalytic residue clusters. Each point represents a predicted cluster, colored by geometric type: Compact (blue), Planar (yellow), Encircling (green), Extended (pink). **(b)** Radar plot comparing four cluster geometries. Features include cluster Radius, Size, and Maximum inter-residue distance. **(c)** Heatmap showing enrichment of cluster types across EC classes. **(d)** Representative examples of Encircling clusters from different EC classes: EC1 (PDB: 1DII, Chain A), EC3 (1EI5, Chain A), EC4 (1B93, Chain B). Top, cartoon representation of the protein structure with catalytic residues shown as sticks. Bottom, close-up views of predicted cluster residues (pink) overlaid with nearby true catalytic residues (yellow), with cluster radius ($Å$) indicated. **(e)** Examples of Encircling (left) and Extended (right) clusters in the EC3.5.1 family. Close-up views show the spatial overlap between predicted cluster residues (pink) and true catalytic residues (sticks). **(f)** Examples of Encircling and Planar clusters in the EC2.1.1 family. Predicted clusters are shown together with true catalytic residues (sticks).

**Interpretability analysis reveals the Structural and Biophysical basis of SCOPE predictions**

We performed interpretability analyses to identify the sequence and structural signals driving SCOPE predictions and to assess whether the model captures the spatial organisation patterns of catalytic residues. We first applied GNNExplainer ^[11]^, a post hoc interpretability method for graph neural networks that quantifies the importance of input features by constructing a compact explanatory subgraph, to assess the importance of four categories of node features: PSSM, HMM, atom-level, and Sequence-level. As shown in **Fig. S4a,** PSSM features contributed most strongly, followed by sequence features and atom-level features, whereas HMM features showed the lowest importance. These results indicate that SCOPE primarily relies on evolutionary conservation and primary sequence information, with additional contributions from structural and physicochemical descriptors. This is consistent with previous evidence that catalytic residues are often highly conserved across homologous enzymes ^[12]^.

To further interpret atom-level features, we examined the contributions of their seven constituent dimensions (**Fig. S4b**). The importance scores varied substantially across atomic attributes, with the aromatic ring indicator (ring) and the B-factor contributing most prominently. This suggests that SCOPE exploits atomic-level properties associated with molecular interaction potentials and local structural stability, thereby providing biophysical interpretability.

We next examined the edge-level explanations to determine whether the residue-residue interactions prioritised by SCOPE are spatially associated with catalytic sites. As shown in **Fig. S4c**, predicted catalytic residues are highly enriched in regions connected by high-importance edges (left), whereas their proportion is markedly reduced in regions associated with low-importance edges. This indicates that the connectivity patterns identified as important by SCOPE are concentrated around catalytic regions. To construct residue-level spatial graphs, a distance threshold of 8$Å$ was used to define edges between residues. This threshold was chosen to slightly extend beyond the median nearest-neighbour distance of catalytic residues (6 $\text{Å}$), thereby capturing not only direct catalytic contacts but also the surrounding local structural neighbourhood involved in active-site organisation. To further quantify this spatial, we constructed residue-level spatial graphs using an 8$Å$ distance threshold and defined spatial clusters as connected components. As shown in **Fig. S4d**, both experimentally annotated and SCOPE-predicted catalytic residues formed substantially larger clusters than random or non-catalytic residues, as measured by average and maximum cluster size. This demonstrates that catalytic residues form compact structural units in three-dimensional space and that SCOPE preserves this organisation in its predictions.

To validate the structural plausibility of these explanations, we selected three representative proteins for which all catalytic residues were correctly predicted and visualised their three-dimensional structures. In glycerol-3-phosphate acyltransferase 1K30, an enzyme involved in lipid metabolism ^[13]^. The high-importance substructure formed a continuous local region that tightly enclosed the catalytic residues H139 and D144 and extended into the surrounding structural environment (**Fig. S4e**). The key structural regions emphasised by SCOPE exhibit a consistent spatial localisation with the true catalytic sites in three-dimensional space. A similar pattern was observed for caseinolytic protease P, ClpP (1TYF), which catalyses peptide bond hydrolysis ^[14]^. As shown in **Fig. S4f**, the experimentally annotated catalytic residues G68, S97, M98, H122 and D171 overlapped with the high-importance substructure identified by SCOPE, suggesting that the model captures continuous active-site regions rather than isolated residue-level signals. In N-carbamoylsarcosine amidohydrolase (1NBA) ^[15]^, the distance-based catalytic residue cluster showed strong concordance with the important substructure and was surrounded by nearby non-catalytic residues, together forming a complete local reaction environment (**Fig. S4g**). Together, these interpretability analyses indicate that SCOPE focuses on spatially coherent multi-residue structural units rather than isolated residues.

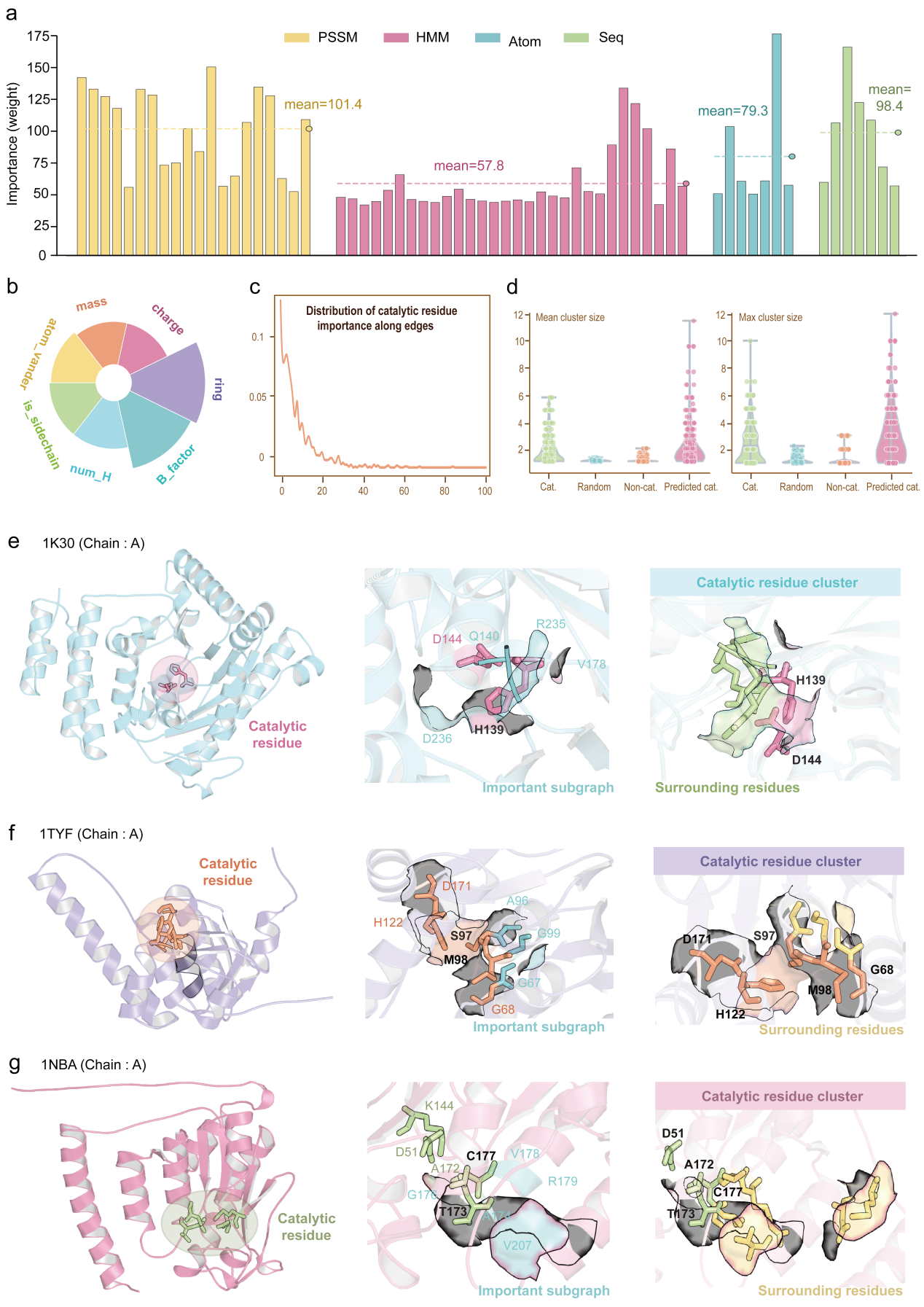

**Fig. S4 Interpretability analysis of SCOPE using GNNExplainer. (a)** Bar plots showing the importance of the four node-feature categories identified by GNNExplainer. The dashed horizontal line in each block denotes the mean importance of each feature group, with the value annotated. **(b)** Pie chart illustrating the contributions of the seven atom-level features, including atomic mass, B-factor, side-chain indicator, atomic charge, hydrogen-bonding capacity, aromatic ring membership and van der Waals radius. **(c)** Distribution of predicted catalytic residues along ranked edge importance. **(d)** Spatial clustering statistics of residues based on a distance threshold of 8 $Å$. **(e-g)** Structural visualization of representative proteins: 1K30 (chain A), 1TYF (chain A) and 1NBA (chain A). Left: full protein structure with true catalytic residues highlighted. Middle: important subgraph extracted from high-importance edges identified by GNNExplainer. Right: spatial catalytic residue cluster and surrounding residues.

**SCOPE generalises to low-homology and structurally diverse enzymes**

Given the pronounced spatial clustering of catalytic residues in enzyme structures and the fact that such spatial organisation is typically independent of overall sequence similarity or fold type, we next evaluated whether SCOPE could identify catalytic residues in enzymes with substantial sequence and structural divergence.

To assess the structural generalisation of SCOPE, we focused on test enzymes with low structural similarity to the training set, defined by TM-score < 0.4. As shown in **Fig. S5a** for Mucorpepsin (PDB ID = 1MPP, Chain A) ^[16]^, L-Fuculose-1-phosphate aldolase (PDB: 1FUA, Chain A) ^[17]^, and the cofactor-dependent phosphoglycerate mutase (PDB: 1E58, Chain A) ^[18]^, SCOPE correctly identified key catalytic residues despite marked differences in overall protein fold.

We further stratified the test enzymes into three sequence-identity ranges (0-30%, 30-70%, and 70-100%) relative to the training set. SCOPE maintains high AUC and AUPRC values even in the low-homology group (0-30%) (**Fig. S5b**). These results suggest that SCOPE retains predictive ability in the lower-identity group; a complementary analysis using stricter train-test identity thresholds is reported below. We also assessed model performance across seven major enzyme functional classes (**Fig. S5c**). To evaluate functional generalizability, we also assessed SCOPE across the seven first-level Enzyme Commission (EC) classes: oxidoreductases, transferases, hydrolases, lyases, isomerases, ligases, and translocases ^[19]^. SCOPE achieved stable, comparable performance across all enzyme classes, with consistently high AUC values. This suggests that the model captures structural features associated with enzymatic catalysis.

Taken together, these results indicate that SCOPE accurately identifies catalytic residues in enzymes with both low sequence identity and low structural similarity to the training set. These analyses describe SCOPE predictions across the examined sequence-similarity groups and structural examples. More stringent sequence-identity sensitivity analyses and CATH homologous-superfamily-disjoint retraining are reported separately.

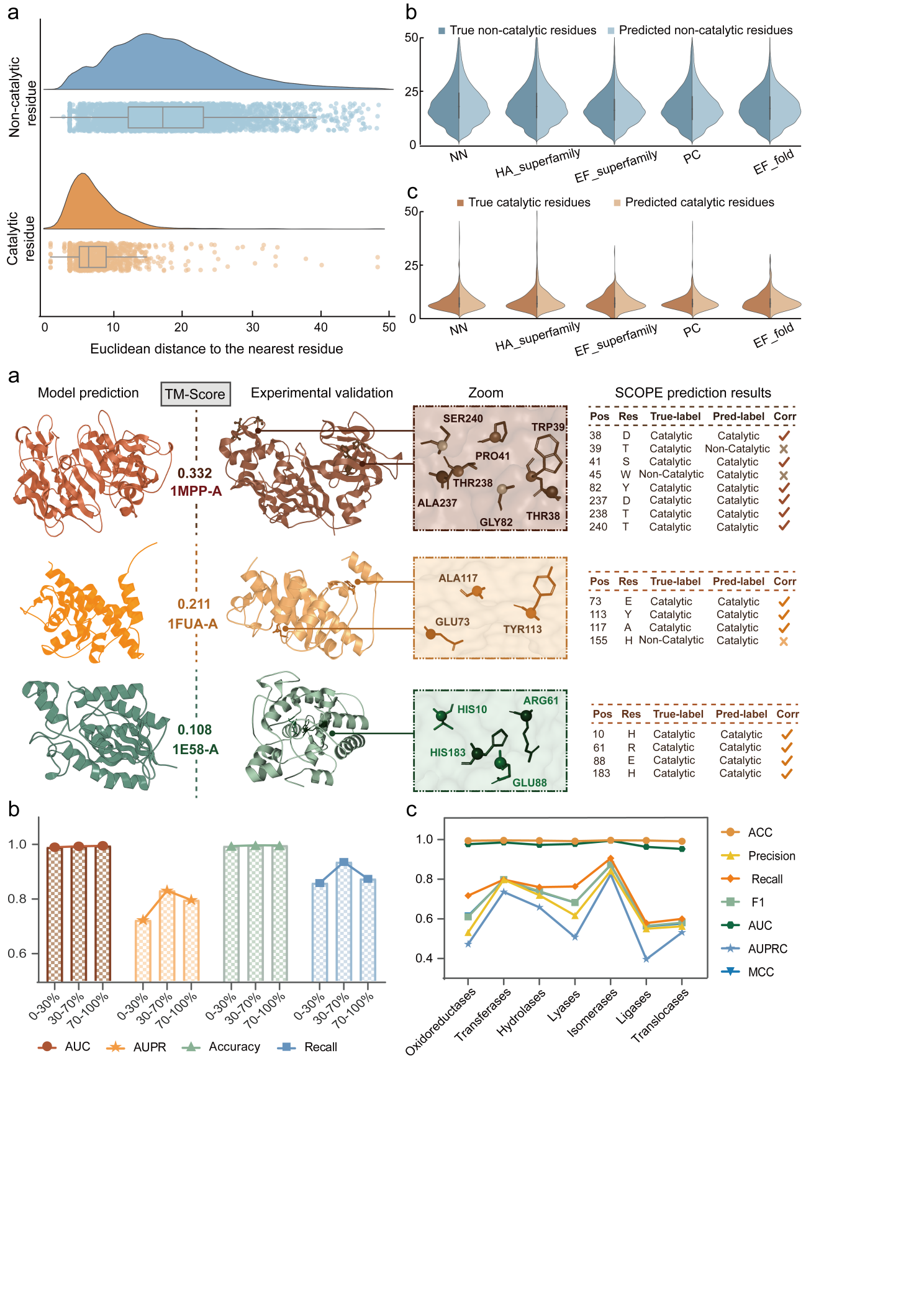

**Fig. S5 (a)** Representative cases illustrating SCOPE predictions under low structural similarity conditions, as measured by TM-score. **(b)** Model performance across diverse sequence identity ranges (0-30%, 30-70%, 70-100%). **(c)** Prediction performance across seven enzyme classes. Different curves denote different performance metrics, including ACC, Precision, Recall, $F_{1}$, AUC, AUPRC, and MCC.

**Robustness to BioEmu-sampled conformational variability**

We evaluated 20 nonredundant enzymes, with four selected from each of the NN, HA-superfamily, EF-superfamily, PC, and EF-fold test sets. Eligible enzymes were 100-350 residues long and had complete catalytic-residue annotations and SCOPE input features. Selection was stratified by sequence length using the fixed random seed, without reference to model predictions. Repeated protein identifiers and identical sequences across test sets were excluded. The selected cohort and candidate audit are provided in **Table S9**.

**Table S9.** Ensemble statistics for the catalytic-residue benchmark cohort (N=20).

| **Dataset** | **Protein_id** | **Sequence_**  **length** | **Valid_**  **conformations** | **mean_pairwise_**  **CA_RMSD_A** | **mean_CA_**  **RMSF_A** | **mean_contact_turnover** |
| --- | --- | --- | --- | --- | --- | --- |
| NN | 1dco-A | 104 | 19 | 1.9772 | 1.0062 | 0.0071 |
|  | 2bmi-A | 232 | 20 | 2.2716 | 1.1578 | 0.0044 |
|  | 1diz-A | 282 | 20 | 0.9847 | 0.5919 | 0.0017 |
|  | 1d8h-A | 311 | 16 | 9.5898 | 4.6167 | 0.0034 |
| HA_  superfamily | 1e2a-C | 105 | 20 | 3.2174 | 3.0302 | 0.0095 |
|  | 1wnw-C | 215 | 16 | 2.9235 | 1.4533 | 0.0051 |
|  | 1dve-A | 267 | 17 | 14.7637 | 8.5268 | 0.0019 |
|  | 1qmh-B | 347 | 17 | 2.5685 | 1.1607 | 0.0026 |
| EF_  superfamily | 1kmv-A | 186 | 18 | 1.4966 | 0.9349 | 0.0049 |
|  | 1jje-A | 220 | 20 | 1.7205 | 0.9014 | 0.0043 |
|  | 1fsg-A | 233 | 14 | 4.3868 | 2.2993 | 0.0059 |
|  | 1j33-A | 333 | 20 | 1.3066 | 0.7526 | 0.0024 |
| PC | 1bs4-A | 168 | 19 | 3.5098 | 1.6757 | 0.0061 |
|  | 1grc-A | 212 | 20 | 1.9075 | 1.1236 | 0.0044 |
|  | 1wgi-A | 286 | 20 | 1.6936 | 0.9356 | 0.0031 |
|  | 1mla-A | 309 | 20 | 1.3463 | 0.8548 | 0.0031 |
| EF_fold | 1hnj-A | 174 | 18 | 2.595 | 1.5407 | 0.0068 |
|  | 1hsk-A | 194 | 20 | 2.0354 | 1.2483 | 0.0061 |
|  | 1jd0-A | 260 | 19 | 1.8256 | 1.0651 | 0.0039 |
|  | 1f4l-A | 350 | 17 | 3.3201 | 1.7734 | 0.0027 |

For each enzyme, 20 conformations were generated using BioEmu version 1.1.0 with the bioemu-v1.1 checkpoint, and all conformations passing BioEmu's default structural filters were retained. Following Cα superposition, the conformation with the smallest mean pairwise Cα RMSD was selected as the label-independent medoid. Frozen full SCOPE, SCOPE without Gaussian smoothing, and SCREEN were evaluated using the experimental PDB structure, the BioEmu medoid, and the complete BioEmu ensemble. For each generated conformation, only the 6Å Cα contact graph was recalculated; all sequence and residue features were held constant. Ensemble predictions were obtained by equally averaging residue probabilities across all valid conformations. The softmax temperature and decision threshold were fixed at 8 and 0.5, respectively.

Per-enzyme AUPRC was the primary endpoint, while AUROC, precision, recall, F1, and MCC were secondary metrics. Robustness was summarized from catalytic-residue detection consistency, conformer-wise probability variation, predicted-set Jaccard similarity, and variation in single-conformation AUPRC. The enzyme was treated as the independent statistical unit. Paired conditions were compared using two-sided Wilcoxon signed-rank tests with Holm correction, and 95% confidence intervals were estimated from 10,000 enzyme-level bootstrap resamples.

All 20 enzymes completed BioEmu generation and model evaluation without failure. A total of 370 valid conformations were retained, corresponding to a mean of 18.5 conformations per enzyme and a range of 14-20. The mean per-enzyme pairwise Cα RMSD was 3.272Å, and the mean per-enzyme Cα RMSF was 1.832Å, confirming that the generated ensembles contained measurable backbone variation (**Table S9**).

SCOPE achieved macro-AUPRC values of 0.7109 using the experimental PDB structures, 0.7121 using the BioEmu medoids, and 0.7118 using ensemble-averaged predictions **Table S10**). The paired mean AUPRC difference between the BioEmu ensemble and the experimental PDB input was +0.00091 (95% enzyme-level bootstrap CI, $\text{-}$0.00013 to +0.00280; Holm-adjusted P = 1.000). Thus, replacing the experimental structures with representative generated conformations or ensemble-averaged predictions did not reduce SCOPE performance in this subset. Under the ensemble condition, the macro-AUPRC values of SCOPE without Gaussian smoothing and SCREEN were 0.7202 and 0.6774, respectively. Paired comparisons between full SCOPE and either model were not statistically significant after Holm correction (both adjusted P = 0.8655).

**Table S10.** Model performance across structure inputs.

| **Model** | **Condition** | **AUPRC_**  **mean** | **AUPRC_**  **median** | **Macro_AUPRC_**  **bootstrap_CI_low** | **Macro_AUPRC_bootstrap_CI_high** |
| --- | --- | --- | --- | --- | --- |
| SCOPE | BioEmu_ensemble_mean | 0.7118 | 0.7903 | 0.5701 | 0.8386 |
|  | BioEmu_medoid | 0.7121 | 0.7903 | 0.5675 | 0.8396 |
|  | experimental_PDB | 0.7109 | 0.7903 | 0.572 | 0.8379 |
| wo_Gaussian | BioEmu_ensemble_mean | 0.7202 | 0.8208 | 0.5709 | 0.8509 |
|  | BioEmu_medoid | 0.7194 | 0.8208 | 0.574 | 0.8498 |
|  | experimental_PDB | 0.7202 | 0.8208 | 0.5757 | 0.8524 |
| SCREEN | BioEmu_ensemble_mean | 0.6774 | 0.7375 | 0.5209 | 0.8231 |
|  | BioEmu_medoid | 0.6185 | 0.5742 | 0.4611 | 0.7718 |
|  | experimental_PDB | 0.616 | 0.5636 | 0.4601 | 0.7717 |

Full SCOPE showed limited conformer-to-conformer prediction variation. The mean fraction of annotated catalytic residues detected in at least 80% of valid conformations was 0.688333, and the mean pairwise Jaccard similarity of predicted-positive residue sets was 1.000. The mean probability standard deviations for catalytic and noncatalytic residues were 0.000405 and 0.000239, respectively, while the mean single-conformation AUPRC IQR was 0.000365 (**Table S11**). The corresponding probability standard deviations were 0.001042 and 0.000882 for SCOPE without Gaussian smoothing and 0.018134 and 0.003637 for SCREEN. Their mean AUPRC IQR values were 0.003160 and 0.038755, respectively. Although the no-Gaussian model had a slightly higher stable catalytic-residue fraction than full SCOPE, full SCOPE showed smaller probability and AUPRC fluctuations across conformations. These results indicate that SCOPE is robust to the BioEmu-sampled conformational and contact-graph perturbations.

**Table S11.** Conformer-wise prediction stability for SCOPE, wo Gaussian, and SCREEN.

| **model** | **SCOPE** | **Wo Gaussian** | **SCREEN** | **SCOPE** |
| --- | --- | --- | --- | --- |
| valid_conformations | 18.5 | 18.5 | 18.5 | 18.5 |
| fraction_catalytic_detection_frequency_ge_0.8 | 0.688333 | 0.713333 | 0.550833 | 0.688333 |
| prediction_SD_catalytic_mean | 0.000405 | 0.001042 | 0.018134 | 0.000405 |
| prediction_SD_noncatalytic_mean | 0.000239 | 0.000882 | 0.003637 | 0.000239 |
| pairwise_Jaccard_mean | 1 | 0.99614 | 0.970319 | 1 |
| conformer_AUPRC_IQR | 0.000365 | 0.00316 | 0.038755 | 0.000365 |
| conformer_AUPRC_range | 0.001125 | 0.010311 | 0.153501 | 0.001125 |
| Spearman_RMSF_prediction_SD | 0.284024 | 0.176242 | 0.112437 | 0.284024 |
| Spearman_contact_turnover_prediction_SD | 0.340221 | 0.365581 | 0.365001 | 0.340221 |

**Sensitivity analysis under stricter train-test sequence-identity thresholds**

To assess the sensitivity of SCOPE performance to residual train-test homology, each sequence from the five benchmark datasets was searched against all 1,054 proteins used for model development, comprising 948 training and 106 validation proteins, using BLASTP v2.17.0+. Maximum sequence identity was determined from alignments covering at least 70% of both the query and subject sequences. Nested test subsets were defined using maximum identity thresholds of <40%, <30%, and <25%. Identical sequences occurring across benchmark datasets were counted once in the primary combined analysis, while dataset-specific results were retained for descriptive comparison.

Performance was summarized using pooled residue-level AUPRC, AUROC, F1 and MCC, together with the mean of per-enzyme AUPRC values. Confidence intervals for enzyme-macro AUPRC were estimated using 2,000 enzyme-level bootstrap resamples with the fixed random seed.

The five benchmark datasets contained 439 evaluable dataset records corresponding to 324 unique enzyme sequences. The <40%, <30%, and <25% subsets retained 114, 87, and 75 unique enzymes, respectively (**Table S12**). Pooled AUPRC decreased from 0.637 in the original combined set to 0.579, 0.527, and 0.515 under these progressively stricter thresholds. At <25% identity, enzyme-macro AUPRC was 0.551 (95% confidence interval, 0.466-0.632), with AUROC, F1 and MCC values of 0.950, 0.521 and 0.517, respectively. This subset contained 22,771 residues, including 220 catalytic residues, corresponding to a catalytic-residue prevalence of 0.97%.

**Table S12.** Combined sequence-deduplicated sensitivity analysis.

| **Metric** | **Original** | **<40%** | **<30%** | **<25%** |
| --- | --- | --- | --- | --- |
| Enzymes (n) | 324 | 114 | 87 | 75 |
| Residues (n) | 112,557 | 34,747 | 26,519 | 22,771 |
| Catalytic residues (n) | 1,106 | 355 | 262 | 220 |
| Catalytic prevalence | 0.98% | 1.02% | 0.99% | 0.97% |
| Pooled AUPRC | 0.637 | 0.579 | 0.527 | 0.515 |
| Macro-AUPRC | 0.686 | 0.611 | 0.569 | 0.551 |
| 95% CI lower | 0.649 | 0.546 | 0.495 | 0.466 |
| 95% CI upper | 0.723 | 0.680 | 0.647 | 0.632 |
| Pooled AUROC | 0.979 | 0.964 | 0.956 | 0.950 |
| Precision | 0.606 | 0.544 | 0.505 | 0.492 |
| Recall | 0.765 | 0.623 | 0.565 | 0.555 |
| F1 | 0.676 | 0.581 | 0.533 | 0.521 |
| MCC | 0.677 | 0.578 | 0.529 | 0.517 |

At the <25% threshold, NN, HA-superfamily, EF-superfamily, PC and EF-fold retained 7, 28, 28, 3 and 22 enzymes, respectively (**Table S13**). Because several dataset-specific subsets were small, the combined sequence-deduplicated analysis was treated as the primary sensitivity result. These findings support predictive ability on remote-homology enzymes, while not excluding shared folds, functional similarity, or evolutionary information incorporated through PSSM and HMM features.

**Table S13.** Dataset-specific sequence-identity sensitivity analysis.

| **Dataset** | **Maximum identity** | **Enzymes (n)** | **Pooled AUPRC** | **Macro-AUPRC (95% CI)** | **AUROC** | **F1** | **MCC** |
| --- | --- | --- | --- | --- | --- | --- | --- |
| NN | Original | 94 | 0.668 | 0.736 (0.674–0.798) | 0.990 | 0.726 | 0.730 |
|  | <40% | 13 | 0.783 | 0.759 (0.577–0.924) | 0.986 | 0.737 | 0.733 |
|  | <30% | 9 | 0.768 | 0.702 (0.448–0.928) | 0.986 | 0.687 | 0.683 |
|  | <25% | 7 | 0.805 | 0.746 (0.491–0.974) | 0.984 | 0.711 | 0.708 |
| HA-  superfamily | Original | 188 | 0.639 | 0.675 (0.624–0.725) | 0.980 | 0.689 | 0.691 |
|  | <40% | 35 | 0.533 | 0.475 (0.342–0.598) | 0.957 | 0.488 | 0.487 |
|  | <30% | 29 | 0.431 | 0.437 (0.299–0.584) | 0.948 | 0.398 | 0.397 |
|  | <25% | 28 | 0.445 | 0.445 (0.302–0.581) | 0.946 | 0.405 | 0.405 |
| EF-  superfamily | Original | 60 | 0.577 | 0.660 (0.576–0.740) | 0.971 | 0.610 | 0.608 |
|  | <40% | 48 | 0.568 | 0.637 (0.540–0.729) | 0.966 | 0.593 | 0.589 |
|  | <30% | 34 | 0.504 | 0.595 (0.480–0.700) | 0.957 | 0.536 | 0.530 |
|  | <25% | 28 | 0.508 | 0.591 (0.475–0.704) | 0.954 | 0.546 | 0.541 |
| PC | Original | 48 | 0.673 | 0.752 (0.654–0.837) | 0.990 | 0.751 | 0.753 |
|  | <40% | 6 | 0.782 | 0.680 (0.401–0.952) | 0.986 | 0.700 | 0.696 |
|  | <30% | 4 | 0.579 | 0.520 (0.185–0.856) | 0.975 | 0.500 | 0.498 |
|  | <25% | 3 | 0.658 | 0.624 (0.161–1.000) | 0.975 | 0.583 | 0.587 |
| EF-fold | Original | 49 | 0.630 | 0.703 (0.614–0.789) | 0.975 | 0.638 | 0.637 |
|  | <40% | 37 | 0.637 | 0.715 (0.617–0.808) | 0.970 | 0.643 | 0.640 |
|  | <30% | 27 | 0.618 | 0.693 (0.561–0.807) | 0.965 | 0.643 | 0.640 |
|  | <25% | 22 | 0.671 | 0.682 (0.535–0.816) | 0.959 | 0.657 | 0.653 |

**Comparison of protein language model embeddings**

We compared frozen residue embeddings from ProtT5-XL-UniRef50, ESM-2 (esm2_t33_650M_UR50D), and ProtBERT-BFD while retaining the same structural graphs, non-PLM features, and downstream SCOPE architecture. ProtT5 and ProtBERT-BFD embeddings had 1,024 dimensions, whereas ESM-2 embeddings had 1,280 dimensions; each was projected to 512 dimensions using a trainable linear layer.

The three downstream models were trained separately using the same 948 training and 106 validation proteins, random seed, Adam learning rate (5×${10}^{-5}$), and 500-epoch budget. Checkpoints were selected solely by validation pooled AUPRC. The graph cutoff, maximum input length, Gaussian auxiliary-loss weight, softmax temperature, and decision threshold were fixed at 6 Å, 1,000 residues, ${10}^{-4}$, 8, and 0.5, respectively. Evaluation used the same 94, 188, 60, 48, and 49 protein records in NN, HA-superfamily, EF-superfamily, PC, and EF-fold. AUPRC was calculated by trapezoidal integration of the precision-recall curve; macro-AUPRC was the unweighted mean of per-protein AUPRC values. AUROC, precision, recall, F1, and MCC were also reported.

ProtT5 achieved pooled AUPRC values of 0.6631, 0.6124, 0.5525, 0.6510, and 0.5606 across the five test sets, compared with 0.5974, 0.5211, 0.4968, 0.6501, and 0.4815 for ESM-2 and 0.5877, 0.5330, 0.4128, 0.6072, and 0.3904 for ProtBERT-BFD, respectively. ProtT5 also had the highest macro-AUPRC, F1, and MCC on each test set, although precision was not consistently highest (**Table S14**).

**Table S14.** Performance of SCOPE with alternative protein language model embeddings.

| **Dataset** | **PLM** | **Pooled AUPRC** | **Macro-**  **AUPRC** | **AUROC** | **F1** | **MCC** |
| --- | --- | --- | --- | --- | --- | --- |
| NN | ProtT5 | 0.6631 | 0.6895 | 0.9906 | 0.6771 | 0.6819 |
|  | ESM-2 | 0.5974 | 0.6422 | 0.9868 | 0.5496 | 0.5513 |
|  | ProtBERT-BFD | 0.5877 | 0.6396 | 0.9845 | 0.5897 | 0.5855 |
| HA-superfamily | ProtT5 | 0.6124 | 0.6587 | 0.9840 | 0.6339 | 0.6373 |
|  | ESM-2 | 0.5211 | 0.6039 | 0.9801 | 0.4924 | 0.4973 |
|  | ProtBERT-BFD | 0.5330 | 0.5951 | 0.9804 | 0.5296 | 0.5254 |
| EF-superfamily | ProtT5 | 0.5525 | 0.5947 | 0.9777 | 0.5685 | 0.5743 |
|  | ESM-2 | 0.4968 | 0.5523 | 0.9725 | 0.5141 | 0.5122 |
|  | ProtBERT-BFD | 0.4128 | 0.5306 | 0.9708 | 0.4672 | 0.4615 |
| PC | ProtT5 | 0.6510 | 0.7137 | 0.9896 | 0.7059 | 0.7085 |
|  | ESM-2 | 0.6501 | 0.6760 | 0.9847 | 0.5974 | 0.6054 |
|  | ProtBERT-BFD | 0.6072 | 0.6665 | 0.9866 | 0.6044 | 0.5998 |
| EF-fold | ProtT5 | 0.5606 | 0.6094 | 0.9811 | 0.5714 | 0.5737 |
|  | ESM-2 | 0.4815 | 0.5861 | 0.9747 | 0.4689 | 0.4697 |
|  | ProtBERT-BFD | 0.3904 | 0.5399 | 0.9714 | 0.4575 | 0.4517 |

**Association between AlphaFold confidence and catalytic-residue prediction**

We reused the prediction-independent cohort of 20 nonredundant enzymes selected for the conformational robustness analysis. PDB chains were mapped to UniProt using PDBe/SIFTS, and structures were downloaded from the official AlphaFold Protein Structure Database. Inclusion required at least 95% exact sequence coverage and exact mapping of all catalytic residues. Nineteen enzymes were evaluable, comprising 59 catalytic residues and 4,559 evaluated residues; 1wnw-C was excluded because its catalytic positions did not all match the database sequence. Cα pLDDT was extracted from the PDB B-factor field. Contact graphs and atom features were recalculated while sequence features, model weights, the softmax temperature of 8 and the decision threshold of 0.5 were retained (**Table S15**).

**Table S15.** SCOPE performance using AlphaFold structures.

| **PDB chain** | **Mean catalytic pLDDT** | **Catalytic recall** | **AUPRC** | **Mean catalytic probability** |
| --- | --- | --- | --- | --- |
| 1dco-A | 97.6700 | 1.0000 | 1.0000 | 0.8166 |
| 2bmi-A | 97.6233 | 0.6667 | 0.7639 | 0.6620 |
| 1diz-A | 94.5600 | 1.0000 | 1.0000 | 0.7773 |
| 1d8h-A | 96.2800 | 1.0000 | 1.0000 | 0.6811 |
| 1e2a-C | 98.3425 | 0.7500 | 0.8839 | 0.6041 |
| 1dve-A | 95.4225 | 0.2500 | 0.5101 | 0.2965 |
| 1qmh-B | 92.8800 | 1.0000 | 1.0000 | 0.7105 |
| 1kmv-A | 96.9350 | 1.0000 | 1.0000 | 0.8339 |
| 1jje-A | 96.8600 | 0.5000 | 0.5661 | 0.4478 |
| 1fsg-A | 94.6000 | 0.6000 | 0.6220 | 0.5380 |
| 1j33-A | 98.6200 | 1.0000 | 1.0000 | 0.7028 |
| 1bs4-A | 98.4225 | 0.2500 | 0.1620 | 0.3027 |
| 1grc-A | 97.1550 | 0.7500 | 0.8661 | 0.6592 |
| 1wgi-A | 98.5000 | 0.0000 | 0.0333 | 0.2783 |
| 1mla-A | 98.4400 | 1.0000 | 0.5139 | 0.8109 |
| 1hnj-A | 98.5300 | 0.5000 | 0.6625 | 0.5259 |
| 1hsk-A | 95.3800 | 1.0000 | 1.0000 | 0.6799 |
| 1jd0-A | 98.4800 | 1.0000 | 0.8167 | 0.7274 |
| 1f4l-A | 83.4350 | 0.5000 | 0.7083 | 0.6066 |

Spearman correlations were calculated between catalytic-site pLDDT and residue prediction probability or detection. Confidence intervals were estimated using 10,000 whole-enzyme bootstrap resamples. Enzyme-level analyses related mean catalytic-site pLDDT to catalytic recall, mean catalytic prediction probability, AUPRC and the AUPRC change from experimental to AlphaFold structures. Two-sided enzyme-level P values were Holm-corrected across eight comparisons within each model, covering native and neutralized B-factor inputs. Because the atom features include a normalized B-factor channel, we additionally fixed this channel at 0.5 to examine its direct contribution while retaining the other inputs and frozen weights.

SCOPE achieved macro-AUPRC values of 0.7423 and 0.7426 using experimental and AlphaFold structures, respectively. Catalytic-site pLDDT showed weak associations with prediction probability (Spearman ρ = 0.086; 95% CI, $\text{-}$0.206 to 0.314) and detection (ρ $\text{=}$ 0.036; 95% CI, $\text{-}$0.252 to 0.246). Enzyme-level associations with catalytic recall and AUPRC were nonsignificant after Holm correction. Neutralizing the B-factor channel yielded a macro-AUPRC of 0.7424 and a similar probability association (ρ = 0.086; 95% CI, $\text{-}$0.204 to 0.311). SCOPE detected 1 of 3 catalytic sites with 70 ≤ pLDDT < 90 and 39 of 56 sites with pLDDT ≥90; the former group contained only two enzymes and was treated descriptively (**Table S16**).

**Table S16.** SCOPE performance by structural input.

| **Structural input** | **Macro-AUPRC** | **Macro-AUROC** | **Macro-recall** | **Structural input** |
| --- | --- | --- | --- | --- |
| Experimental PDB | 0.7423 | 0.9844 | 0.7246 | Experimental PDB |
| AlphaFold | 0.7426 | 0.9848 | 0.7246 | AlphaFold |
| AlphaFold; B-factor input = 0.5 | 0.7424 | 0.9848 | 0.7246 | AlphaFold; B-factor input = 0.5 |

**Matched baseline comparison and methodological increments**

SCREEN was evaluated using weights retrained on the same 948 training and 106 validation proteins as SCOPE. For the revised comparison, AEGAN was retrained using the same protein split, with checkpoint selection based on validation AUPRC. Each method retained its own input representation and training configuration. Evaluation was restricted to shared, structurally resolved residue positions within SCOPE’s original 1,000-residue input limit. NN, HA-superfamily, EF-superfamily, PC and EF-fold included 94, 188, 60, 48 and 49 protein records, respectively.

Relative to SCREEN, SCOPE retains the 6 Å Cα contact graph, 64-dimensional local features, ProtT5 embeddings and development corpus. The principal changes are a DeepGCN backbone with residual connections and initial-representation injection, and a Gaussian auxiliary classification objective, replacing SCREEN’s shallower GCN and staged classification/EC-triplet training. Gaussian smoothing operates along residue order on fused graph-derived representations.

Full SCOPE achieved higher pooled and per-protein macro-AUPRC than SCREEN and retrained AEGAN on all five test sets (**Table S17**).

**Table S17.** Performance under matched training splits and residue evaluation positions.

| **Dataset** | **Model** | **Pooled AUPRC** | **Macro-**  **AUPRC** | **AUROC** | **Precision** | **Recall** | **F1** | **MCC** |
| --- | --- | --- | --- | --- | --- | --- | --- | --- |
| NN | SCOPE | 0.6700 | 0.7362 | 0.9900 | 0.6385 | 0.8453 | 0.7275 | 0.7315 |
|  | SCOPE w/o Gaussian | 0.6401 | 0.7359 | 0.9902 | 0.6370 | 0.8395 | 0.7244 | 0.7281 |
|  | SCREEN | 0.6562 | 0.7252 | 0.9908 | 0.6403 | 0.8109 | 0.7155 | 0.7173 |
|  | AEGAN | 0.1962 | 0.3100 | 0.9421 | 0.0562 | 0.8997 | 0.1058 | 0.2016 |
| HA-  superfamily | SCOPE | 0.6442 | 0.6800 | 0.9802 | 0.6179 | 0.7853 | 0.6916 | 0.6934 |
|  | SCOPE w/o Gaussian | 0.6435 | 0.6917 | 0.9811 | 0.6330 | 0.7740 | 0.6965 | 0.6969 |
|  | SCREEN | 0.6278 | 0.6735 | 0.9832 | 0.6447 | 0.7532 | 0.6948 | 0.6938 |
|  | AEGAN | 0.1700 | 0.3019 | 0.9309 | 0.0498 | 0.8686 | 0.0943 | 0.1859 |
| EF-  superfamily | SCOPE | 0.5804 | 0.6617 | 0.9710 | 0.5570 | 0.6769 | 0.6111 | 0.6092 |
|  | SCOPE w/o Gaussian | 0.5929 | 0.6992 | 0.9694 | 0.6179 | 0.6718 | 0.6437 | 0.6401 |
|  | SCREEN | 0.5731 | 0.6369 | 0.9730 | 0.5943 | 0.6462 | 0.6192 | 0.6152 |
|  | AEGAN | 0.1668 | 0.2696 | 0.9211 | 0.0566 | 0.8564 | 0.1061 | 0.1946 |
| PC | SCOPE | 0.6730 | 0.7524 | 0.9902 | 0.6653 | 0.8579 | 0.7494 | 0.7521 |
|  | SCOPE w/o Gaussian | 0.6235 | 0.7217 | 0.9894 | 0.6553 | 0.8415 | 0.7368 | 0.7391 |
|  | SCREEN | 0.6675 | 0.7414 | 0.9905 | 0.6565 | 0.8251 | 0.7312 | 0.7324 |
|  | AEGAN | 0.2289 | 0.3199 | 0.9494 | 0.0650 | 0.9180 | 0.1214 | 0.2197 |
| EF-fold | SCOPE | 0.6326 | 0.7048 | 0.9756 | 0.5864 | 0.7089 | 0.6418 | 0.6402 |
|  | SCOPE w/o Gaussian | 0.6486 | 0.6915 | 0.9699 | 0.6322 | 0.6962 | 0.6627 | 0.6593 |
|  | SCREEN | 0.5956 | 0.6570 | 0.9774 | 0.6131 | 0.6519 | 0.6319 | 0.6278 |
|  | AEGAN | 0.1823 | 0.3123 | 0.9274 | 0.0625 | 0.8291 | 0.1162 | 0.2030 |

**Runtime and memory requirements**

We benchmarked three test proteins, 1smn-A (245 residues), 1nba-A (264 residues), and 1d8c-A (723 residues), starting from their sequences and supplied experimental PDB structures. Each protein was processed once in a fresh process using frozen SCOPE weights. PSI-BLAST and HHblits profiles, ProtT5 embeddings, atom and physicochemical features, and contact graphs were regenerated without reusing previously computed features. Total elapsed time included initialization, feature generation, prediction and output writing. The existing DSSP preprocessing step was also included, although its outputs were not used by SCOPE.

Runs were performed serially on an Intel Xeon Platinum 8369B server with approximately 125 GiB RAM and one NVIDIA GeForce RTX 3090 Ti GPU. PSI-BLAST 2.17.0+ searched UniRef90 using three iterations and 40 threads, while HHblits 3.3.0 searched Uniclust30 (2018_08) using six threads. Peak CPU memory was measured as the maximum summed resident set size of the process tree, and GPU memory was monitored using NVML, both sampled every 50 ms. Structure acquisition or prediction and one-time database/model installation were excluded. Operating-system caches were not controlled, and measurements were obtained on a shared server.

Total elapsed times were 1250.1, 462.1, and 591.7 s for 1smn-A, 1nba-A and 1d8c-A, respectively. Peak CPU memory ranged from 68.73 to 68.99 GiB, and peak GPU memory was approximately 5.69 GiB. PSI-BLAST and HHblits together accounted for 93.8-97.1% of total runtime. These measurements describe the computational cost under the specified conditions when a structure is already available (**Table S18**).

**Table S18.** Breakdown of runtime by processing stage for three benchmark proteins.

| **Stage** | **1smn-A (s)** | **1nba-A (s)** | **1d8c-A (s)** |
| --- | --- | --- | --- |
| input_preparation | 0.0003 | 0.0004 | 0.0662 |
| PSI_BLAST | 1099.7333 | 309.9686 | 495.3731 |
| HHblits | 114.0898 | 123.6417 | 70.9879 |
| profile_conversion | 0.0120 | 0.0144 | 0.0231 |
| structure_and_physicochemical_generation | 2.2120 | 2.0180 | 7.8987 |
| DSSP_GUI_overhead | 0.0351 | 0.0405 | 0.4015 |
| ProtT5_model_initialization | 20.7160 | 14.4238 | 10.2241 |
| ProtT5_embedding_and_write | 3.9160 | 0.7695 | 1.1605 |
| SCOPE_model_initialization | 0.2530 | 0.0476 | 0.0656 |
| feature_read_and_graph_normalization | 0.0362 | 0.0082 | 0.0389 |
| SCOPE_prediction_with_transfer | 0.0970 | 0.0076 | 0.0051 |
| prediction_write | 0.0147 | 0.0014 | 0.0016 |

**Seven-dimensional amino acid encoding**

SCOPE uses a fixed lookup table to assign seven numerical features to each of the 20 standard amino acids. The table was inherited from the SCREEN implementation. Comparison with the archived public SCREEN lookup confirmed agreement of all 140 numerical entries. The complete 20×7 mapping is provided at full numerical precision in seven_descriptor_lookup.json in the SCOPE GitHub repository referenced in the Code availability statement. The channel order and stored values in this file define the encoding used by the trained model.

The channel definitions and numerical correspondences are summarized in **Table S19**. Numerical correspondence to a molecular descriptor does not establish the original descriptor-generation procedure.

In the inherited table, alanine has the same stored values as cysteine in channels 4–6. These entries were retained because changing them would alter the input representation used by the existing model. Their presence in the upstream table establishes their provenance but does not establish why they were originally assigned.

The descriptor reconstruction supports the numerical correspondences in **Table S19**; it does not recover the original generation script or resolve the historical interpretation of every channel. The complete numerical lookup in the SCOPE GitHub repository is therefore the authoritative specification for reproduction. Users should retrieve the stored values from seven_descriptor_lookup.json rather than regenerate the features from molecular descriptors or from the category descriptions in **Table S19**.

For each residue, the seven lookup values were retrieved by amino acid identity and aligned with the sequence position. They were concatenated with the 20-dimensional PSSM profile, 30-dimensional HMM profile, and seven-dimensional atomic structural features to form the 64-dimensional node input to the graph network.

**Table S19.** Definitions and numerical correspondences of the seven amino acid encoding channels.

| **Channel** | **Definition or correspondence** | **Encoding details** |
| --- | --- | --- |
| 1 | Isoelectric-point-related encoding | Matches scaled conventional pI values: (pI $\text{-}$ 2.77)/8.38. |
| 2 | Four-level amino acid grouping | AFILMPVW: 0; CGNQSTY: 1/3; DE: 2/3; HKR: 1. The original biochemical interpretation could not be verified. |
| 3 | Acidic/basic residue grouping | DE: 0.5; HKR: 1; other amino acids: 0. These are category codes, not signed physical charges. |
| 4 | Hydrogen-bond acceptor count | Matches min–max-scaled RDKit acceptor counts for 19 amino acids; alanine is an inherited exception. |
| 5 | Hydrogen-bond donor count | Matches min–max-scaled RDKit donor counts for 19 amino acids; alanine is an inherited exception. |
| 6 | MolLogP | Matches min–max-scaled RDKit MolLogP for 19 amino acids; alanine is an inherited exception. |
| 7 | Topological polar surface area | Matches min–max-scaled RDKit topological polar surface area for all 20 amino acids. |

**Public PafA mutagenesis analysis**

PafA mutagenesis measurements were mapped to the native protein sequence, excluding the fusion tag and linker. The primary endpoint was wild-type-normalized catalytic efficiency for cMUP. For the position-level analysis, the lowest available relative activity across the evaluated glycine and valine substitutions was used. Severe activity loss was defined as relative activity ≤0.1. Censored measurements were assigned to a binary category only when the reported bound permitted an unambiguous classification; continuous correlation analyses used uncensored measurements.

SCOPE, SCOPE without the Gaussian auxiliary objective, SCREEN and a PSSM-derived conservation score were evaluated on matched positions. Experimental measurements were not used to tune model weights or select the prediction threshold. Additional analyses considered the glycine and valine libraries separately, alternative activity cutoffs, structurally resolved positions and the available active-fraction measurements. For structure-based inputs, the modified threonine in the 5TJ3 scaffold was represented by its parent-residue atom subset without adding or moving coordinates.

Among 525 evaluable positions, 91 met the severe-loss criterion. At a probability threshold of 0.5, all six SCOPE-positive positions met this criterion, yielding a precision of 1.000 and a recall of 0.066 for this functional endpoint. SCOPE achieved an average precision of 0.342 and an AUROC of 0.609, compared with 0.299 and 0.574 for the no-Gaussian model and 0.281 and 0.574 for SCREEN. The PSSM-derived conservation score achieved higher ranking performance, with an average precision of 0.455 and an AUROC of 0.793 (**Table S20**).

**Table S20.** Prediction of severe mutational activity loss in PafA.

| **Predictor** | **Average precision** | **AUROC** | **Spearman correlation with activity loss** | **Predicted-positive positions** | **Severe-loss positions identified** | **Precision** | **Recall** |
| --- | --- | --- | --- | --- | --- | --- | --- |
| SCOPE | 0.342 | 0.609 | 0.038 | 6 | 6 | 1.000 | 0.066 |
| SCOPE w/o Gaussian | 0.299 | 0.574 | $\text{-}$0.043 | 8 | 6 | 0.750 | 0.066 |
| SCREEN | 0.281 | 0.574 | $\text{-}$0.028 | 6 | 5 | 0.833 | 0.055 |
| PSSM-derived conservation | 0.455 | 0.793 | 0.468 | — | — | — | — |

**Cohort-wide analysis of prediction transitions**

We compared full SCOPE with the separately trained model without the Gaussian auxiliary objective across 323 unique enzyme sequences after excluding exact sequence matches to the development corpus. Predictions were compared on a shared residue mask at a probability threshold of 0.5, without selecting proteins according to model performance.

Relative to the no-Gaussian model, full SCOPE recovered 37 additional reference-positive residues, whereas 24 reference-positive residues changed to negative predictions. Among reference-negative residues, 83 changed from positive to negative and 137 changed from negative to positive (**Table S21**). Thus, the differences involved both addition and removal of positive predictions, rather than exclusively filtering reference-negative predictions.

**Table S21.** Prediction transitions from the no-Gaussian model to full SCOPE across 323 unique enzyme sequences.

| **Reference annotation** | **No-Gaussian prediction → Full SCOPE prediction** | **Number of residues** |
| --- | --- | --- |
| Positive | Negative → positive | 37 |
| Positive | Positive → negative | 24 |
| Negative | Positive → negative | 83 |
| Negative | Negative → positive | 137 |

**Training-template context of the experimentally tested enzymes**

We examined sequence correspondences between the four experimental enzymes and proteins in the training set. For each query with a significant hit, the template with the highest BLAST bit score was selected using an E-value cutoff of 0.001. Query activity measurements and query catalytic labels were not used for template selection. Sequence identity was calculated over the local alignment; query and template coverage were recorded separately. Experimental positions were mapped through the alignment to determine whether the corresponding template positions carried catalytic annotations.

Fc-TAL2 aligned to 2RJR-A with 35.03% local identity and 99.60% query coverage. Its four tested positions mapped to template residues with the same amino acid identities, but these template positions were not annotated as catalytic in the training labels. CotA aligned to 2QT6-A with 20.00% local identity and 49.51% query coverage. CotA H107 and H155 mapped to annotated template positions H66 and H111, respectively. rsAP aligned to 1EW2-A with 30.99% local identity and 57.07% query coverage; H484 was outside the selected alignment. TGDac had no significant training-set hit under the specified search cutoff (**Table S22**).

A separate search against the published structural motif library recovered an M-CSA 390-associated match from template 1GYC-A that included CotA H107 and H155. This search describes known structural context; it is not an independent accuracy benchmark because the published library was not restricted to the training set.

Lack of an alignment, lack of a transferred annotation, or absence of a reported motif match does not establish mechanistic novelty or evolutionary independence. These analyses distinguish functional validation of selected positions from evidence of generalization to previously unseen catalytic architectures.

**Table S22.** Training-template correspondences for the four experimental enzymes.

| **Enzyme** | **Training template** | **Local identity (%)** | **Query coverage (%)** | **Template coverage (%)** | **Correspondence of tested positions** |
| --- | --- | --- | --- | --- | --- |
| Fc-TAL2 | 2RJR-A | 35.03 | 99.60 | 96.38 | R291, F337, N420 and D489 align to R297, F342, N424 and D493; these template positions lack catalytic annotations. |
| TGDac | No significant hit | — | — | — | No correspondence established under the specified cutoff. |
| CotA | 2QT6-A | 20.00 | 49.51 | 44.58 | H107 and H155 align to annotated catalytic positions H66 and H111. |
| rsAP | 1EW2-A | 30.99 | 57.07 | 70.98 | H484 lies outside the selected local alignment. |

Templates were selected by the highest training-set BLAST bit score among hits meeting the E-value cutoff of 0.001. Absence of a catalytic annotation in the evaluated template labels does not imply absence of biological function. Residue correspondences use the sequence numbering recorded in the alignment audit.

**Statistical testing and multiple-comparison correction**

Annotation-enrichment analyses were performed using two-sided Fisher’s exact tests. The updated cohorts comprised 95 SCOPE complete-recovery records versus 256 background records for Fig. 3c, and 13 GS-dependent versus 82 GS-independent records for Fig. 4c. Complete recovery required no false-negative or false-positive predictions on the matched evaluation positions. GS-dependent records were those that no longer met this criterion when the Gaussian auxiliary objective was omitted.

For Fig. 3c, Benjamini-Hochberg (BH) correction was applied across all 878 annotation terms tested within the cohort analysis, without filtering terms according to their observed P values. For Fig. 4c, the 13 EC-class and subclass comparisons formed a separate correction family. Raw and adjusted P values are reported separately. Statistical significance was assessed using BH-adjusted P < 0.05. Odds ratios describe the direction and magnitude of associations and were interpreted alongside the corresponding adjusted P values.

In the updated SCOPE complete-recovery cohort, EC3.4 annotations occurred in 11/95 foreground records and 8/256 background records, corresponding to an odds ratio of 4.0595. The two-sided Fisher’s exact P value was 0.00560, and the BH-adjusted P value across all 878 tested terms was 0.86728. The EC3, EC3.1 and ≥5-catalytic-residue comparisons also did not reach statistical significance after correction (**Table S23**).

In the updated GS-dependent and GS-independent groups, EC3.4 annotations occurred in 5/13 and 6/82 records, respectively. The odds ratio was 7.9167, with a two-sided P value of 0.00641 and a BH-adjusted P value of 0.08334 across the 13 EC comparisons (**Table S24**). This association did not reach the adjusted significance threshold. Of the 13 GS-dependent records, five contained false-negative catalytic-residue predictions in the model without the Gaussian auxiliary objective, whereas the remaining eight contained false-positive predictions only.

**Table S23.** Annotation-enrichment statistics for the updated cohort in Figure 3c.

| **Annotation** | **SCOPE complete-recovery records** | **Background records** | **Odds ratio** | **Raw P value** | **BH-adjusted P value** |
| --- | --- | --- | --- | --- | --- |
| EC3.4 peptidases | 11/95 | 8/256 | 4.0595 | 0.00560 | 0.86728 |
| EC3 hydrolases | 36/95 | 64/256 | 1.8305 | 0.02330 | 0.86728 |
| ≥5 catalytic residues | 30/95 | 48/256 | 2.0000 | 0.01374 | 0.86728 |
| EC3.1 subclass | 14/95 | 25/256 | 1.5970 | 0.18677 | 0.86728 |

**Table S24.** EC-class and subclass comparisons for the updated cohorts in Figure 4c.

| **Annotation** | **GS-dependent records** | **GS-independent records** | **Odds ratio** | **Raw P value** | **BH-adjusted P value** |
| --- | --- | --- | --- | --- | --- |
| EC1 oxidoreductases | 2/13 | 15/82 | 0.8121 | 1.00000 | 1.00000 |
| EC2 transferases | 3/13 | 20/82 | 0.9300 | 1.00000 | 1.00000 |
| EC3 hydrolases | 6/13 | 30/82 | 1.4857 | 0.54835 | 1.00000 |
| EC4 lyases | 1/13 | 14/82 | 0.4048 | 0.68475 | 1.00000 |
| EC5 isomerases | 2/13 | 4/82 | 3.5455 | 0.18897 | 1.00000 |
| EC6 ligases | 0/13 | 3/82 | 0.0000 | 1.00000 | 1.00000 |
| EC7 translocases | 0/13 | 0/82 | Not estimable | 1.00000 | 1.00000 |
| EC3.1 subclass | 1/13 | 13/82 | 0.4423 | 0.68384 | 1.00000 |
| EC3.2 subclass | 1/13 | 5/82 | 1.2833 | 1.00000 | 1.00000 |
| EC3.4 peptidases | 5/13 | 6/82 | 7.9167 | 0.00641 | 0.08334 |
| EC3.5 subclass | 0/13 | 6/82 | 0.0000 | 0.59187 | 1.00000 |
| EC3.6 subclass | 1/13 | 2/82 | 3.3333 | 0.36018 | 1.00000 |
| EC3.8 subclass | 0/13 | 2/82 | 0.0000 | 1.00000 | 1.00000 |

**Evaluation using a CATH homologous-superfamily-disjoint partition**

We constructed an additional partition of the original development cohort using available CATH homologous-superfamily annotations. Proteins sharing any annotated CATH homologous superfamily, or an identical full-length sequence, were linked and assigned to the same connected component. All annotated domains on each protein chain were considered, rather than only the domain containing catalytic residues. Consequently, proteins connected through multidomain relationships were retained within the same partition.

Components were assigned to training, validation and test sets using a fixed, size-balanced procedure targeting a 70:15:15 allocation, without using model performance. The resulting sets contained 734, 158 and 157 protein records, respectively. Five records lacking the required annotations were excluded from this additional analysis: 1JQN-A, 1APY-A, 5VSM-A, 3A15-A and 1MHL-A. The grouping produced 248 connected components overall; the held-out test set contained 76 components.

AUPRC and average precision were calculated separately for each test protein. For paired comparisons, each protein’s metric was first averaged across the three training seeds. Confidence intervals were obtained by resampling entire connected family components, preserving the dependence among proteins within each component. Paired Wilcoxon tests used component-level mean differences. Holm correction was applied across four comparisons: two metrics and two comparators. Repeated predictions for the same protein across seeds were not treated as independent test samples. The confidence intervals are conditional on the three evaluated seeds and do not represent all sources of training variability.

SCOPE showed a positive AUPRC difference relative to SCREEN for each of the three evaluated seeds. The mean paired difference was 0.04669, with a family-component bootstrap 95% confidence interval of 0.02849-0.06536 and a Holm-adjusted P value of 2.94×10⁻⁶. The corresponding average-precision difference was 0.04425, with a 95% confidence interval of 0.02898-0.05972 (**Table S25**).

Differences between SCOPE and the model without the Gaussian auxiliary objective were smaller. The mean AUPRC difference was 0.00215 and the average-precision difference was 0.00121; both confidence intervals included zero, and neither comparison was significant after correction.

**Table S25.** Paired performance differences under the CATH homologous-superfamily-disjoint evaluation.

| **Comparison** | **Metric** | **Mean paired difference** | **Family-component bootstrap 95% CI** | **Holm-adjusted P value** |
| --- | --- | --- | --- | --- |
| SCOPE- SCREEN | Per-protein AUPRC | 0.04669 | 0.02849 to 0.06536 | 2.94 × 10⁻⁶ |
| SCOPE- SCREEN | Per-protein average precision | 0.04425 | 0.02898 to 0.05972 | 1.75 × 10⁻⁶ |
| SCOPE-no-Gaussian model | Per-protein AUPRC | 0.00215 | $\text{-}$0.00207 to 0.00715 | 1.000 |
| SCOPE-no-Gaussian model | Per-protein average precision | 0.00121 | $\text{-}$0.00169 to 0.00459 | 1.000 |

**Sensitivity to the residue-contact distance threshold**

We evaluated residue-contact thresholds of 4, 6 and 8 Å to assess the sensitivity of SCOPE to graph connectivity. Residue pairs were connected according to their Cα distances, while the remaining model configuration was held constant. Models used the same training and validation split, input features, initialization seed, optimizer and training budget of 500 epochs, with a learning rate of 5 × 10⁻⁵. The Gaussian auxiliary objective was retained with a weight of 10⁻⁴.

The 6 Å model achieved higher pooled AUPRC than the 4 Å model on NN and PC, whereas 4 Å performed better on HA-superfamily and EF-superfamily; EF-fold results were nearly identical. For macro-AUPRC, 4 Å performed better on four datasets and 6 Å on PC. The 8 Å model performed worse than 6 Å on both measures across all five datasets. These results indicate dataset- and metric-dependent sensitivity rather than a universally optimal cutoff. The original 6 Å setting was retained for the main model, with all evaluated configurations reported in **Table S26**.

**Table S26.** Sensitivity of SCOPE to the residue-contact distance threshold.

| **Dataset** | **No. of proteins** | **Pooled AUPRC 4 Å** | **Pooled AUPRC 6 Å** | **Pooled AUPRC 8 Å** | **Macro-AUPRC 4 Å** | **Macro-AUPRC 6 Å** | **Macro-AUPRC 8 Å** |
| --- | --- | --- | --- | --- | --- | --- | --- |
| NN | 93 | 0.6527 | 0.6642 | 0.6316 | 0.7152 | 0.6941 | 0.6738 |
| HA-  superfamily | 187 | 0.6197 | 0.6170 | 0.5758 | 0.6803 | 0.6646 | 0.6237 |
| EF-  superfamily | 60 | 0.5723 | 0.5560 | 0.5281 | 0.6257 | 0.5995 | 0.5940 |
| PC | 48 | 0.6317 | 0.6498 | 0.6460 | 0.7114 | 0.7141 | 0.6778 |
| EF-fold | 49 | 0.5681 | 0.5681 | 0.5237 | 0.6187 | 0.6115 | 0.6064 |

**Comparison of auxiliary smoothing kernels**

Gaussian weighting provides a symmetric, smoothly decaying weighting profile that emphasizes the central position and nearby sequence positions. This is a design rationale for local aggregation, not a guarantee of superior predictive performance. We therefore compared the Gaussian kernel with uniform and triangular weighting. These kernels operate along sequence order on graph-derived representations; three-dimensional contact information is encoded by the graph network rather than by the smoothing distance itself.

Gaussian weighting yielded higher pooled and macro-AUPRC than triangular weighting on all five datasets. Comparisons with uniform weighting were mixed: Gaussian yielded higher pooled AUPRC on PC and EF-fold, whereas uniform weighting yielded higher values on NN, HA-superfamily and EF-superfamily. For macro-AUPRC, Gaussian was higher on HA-superfamily, while uniform weighting was higher on the other four datasets (**Tables S27-S28**).

**Table S27. Pooled residue-level AUPRC with alternative auxiliary smoothing kernels at a fixed 6 Å graph cutoff.**

| **Dataset** | **No. of proteins** | **Gaussian** | **Uniform** | **Triangular** |
| --- | --- | --- | --- | --- |
| NN | 93 | 0.6642 | 0.6657 | 0.6307 |
| HA-superfamily | 187 | 0.6170 | 0.6172 | 0.5772 |
| EF-superfamily | 60 | 0.5560 | 0.5618 | 0.5243 |
| PC | 48 | 0.6498 | 0.6449 | 0.6487 |
| EF-fold | 49 | 0.5681 | 0.5662 | 0.5224 |

**Table S28. Macro-AUPRC with alternative auxiliary smoothing kernels at a fixed 6 Å graph cutoff.**

| **Dataset** | **No. of proteins** | **Gaussian** | **Uniform** | **Triangular** |
| --- | --- | --- | --- | --- |
| NN | 93 | 0.6941 | 0.6959 | 0.6732 |
| HA-superfamily | 187 | 0.6646 | 0.6573 | 0.6195 |
| EF-superfamily | 60 | 0.5995 | 0.6121 | 0.5861 |
| PC | 48 | 0.7141 | 0.7233 | 0.6792 |
| EF-fold | 49 | 0.6115 | 0.6185 | 0.6069 |

**Training-template-restricted structural motif comparison**

We constructed a structural motif library exclusively from the training records and catalytic annotations of the CATH-grouped partition. One template containing the full annotated catalytic residue set was constructed per eligible training chain. Sequence-to-structure mapping retained residue pairs shared by all optimal global alignments; insertion-coded catalytic residues were excluded. Templates were successfully constructed for 703 of 734 training records. Construction failed for 28 records, and three records were excluded under the mapping criteria. These records and their exclusion reasons were retained in the audit. Test-set catalytic annotations were not used for template construction or selection.

Matching used EnzyMM 0.3.2 with frozen published defaults, including its match-quality filter. Accepted matches were converted to binary residue predictions by taking the union of matched positions. All 157 held-out proteins were retained, including proteins with no accepted matches. Comparisons used the same 48,960 exactly mapped, resolved residue positions within the first 1,000 reference sequence positions.

Average precision (AP), rather than trapezoidal AUPRC, was used for the ranking comparison. Pooled AP was calculated over the combined evaluated residues, and macro-AP was the unweighted mean of per-protein AP values. EnzyMM supplied binary match indicators, whereas the neural models supplied continuous scores; AP therefore summarizes different score resolutions. Neural-model metrics in the table are arithmetic means across the three seeds, not metrics calculated after averaging residue probabilities.

The restricted-template EnzyMM analysis identified eight positive residue positions, of which four carried catalytic reference annotations, giving a pooled precision of 0.5000 and recall of 0.0061. Pooled AP was 0.0164 and macro-AP was 0.0211. The neural-model results on the same evaluated residue positions are reported in **Table S29**.

**Table S29.** Structural motif and neural-model predictions on matched residues in the CATH-held-out set.

| **Method** | **Pooled AP** | **Macro-AP** | **Pooled precision** | **Pooled recall** |
| --- | --- | --- | --- | --- |
| SCOPE | 0.3965 | 0.5508 | 0.5140 | 0.3237 |
| SCOPE without Gaussian auxiliary objective | 0.3963 | 0.5496 | 0.5144 | 0.3247 |
| SCREEN | 0.3738 | 0.5068 | 0.5075 | 0.2830 |
| EnzyMM, training-template-restricted | 0.0164 | 0.0211 | 0.5000 | 0.0061 |

**Illustrative prediction comparisons within known catalytic architectures**

We present 1FJS and 1EI5 as examples of how the evaluated predictors recover reference-annotated catalytic residues within known active-site architectures. These examples illustrate differences among predictor outputs and are not used to establish previously unknown catalytic mechanisms, superiority over homology- or motif-based identification, or independent generalization. Overall predictive performance is assessed through the quantitative benchmark analyses.

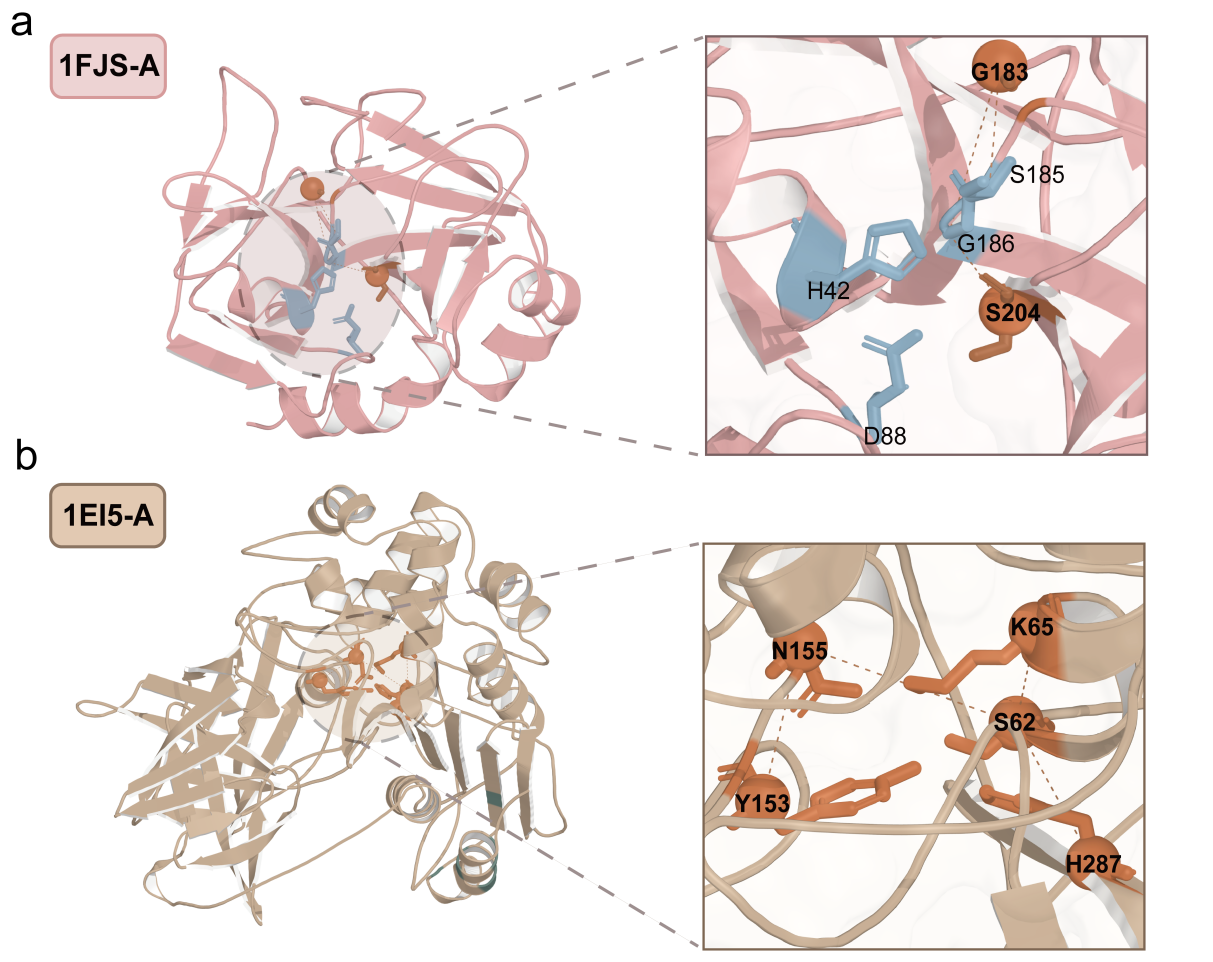

**Supplementary Fig. S6. Illustrative prediction comparisons for enzymes with known catalytic architectures. a,** Predictions for 1FJS chain A. Blue residues indicate reference-annotated catalytic residues identified by both SCOPE and SCREEN; orange residues indicate annotated positions identified by SCOPE but missed by SCREEN. **b,** Predictions for 1EI5 chain A. Orange residues indicate reference-annotated catalytic positions identified by SCOPE within the active-site pocket.

Protein backbones are shown as cartoons and highlighted residues as sticks. Enlarged views show the positions of the annotated residues within the known active-site architectures. These selected examples provide a descriptive comparison of predictor outputs rather than an estimate of comparative performance across enzymes.

**Published weight comparisons on the original benchmarks**

We evaluated SCOPE and published Squidly and EasIFA weights on the original NN, HA-superfamily, EF-superfamily, PC and EF-fold benchmark datasets. Values were calculated from predictions generated in this study rather than copied from the original tool publications. No model weights were retrained for these published-weight comparisons. Each comparison used the available paired records and matched residue evaluation for the two methods. Macro-AUPRC denotes the unweighted mean of per-record trapezoidal precision–recall area, not average precision.

The two comparator analyses had different coverage and were not treated as a three-method comparison on a common cohort. Exact-sequence checks against available EasIFA pretraining and finetuning development tables identified known matches for 68 of the 72 evaluated records.

SCOPE achieved higher macro-AUPRC than the published Squidly 3B ensemble on the paired records from all five original datasets (**Table S30**). Against EasIFA, SCOPE had higher macro-AUPRC on the available NN, HA-superfamily, EF-superfamily and PC subsets, whereas EasIFA had higher macro-AUPRC on EF-fold (**Table S31**).

These original-benchmark results differ from the external CataloDB evaluation, which used different test sequences and CataloDB-specific Squidly weights. Unequal training corpora, model-specific input requirements, unresolved provenance and evaluation coverage limit attribution of the observed differences to model architecture. Results are therefore interpreted under their stated conditions rather than as evidence of universal superiority.

**Table S30.** SCOPE and the published Squidly ensemble.

| **Dataset** | **Model** | **AUPRC** | **Recall** | **F1** |
| --- | --- | --- | --- | --- |
| NN | SCOPE | 0.7362 | 0.8536 | 0.7311 |
| NN | Squidly | 0.4876 | 0.3059 | 0.3749 |
| HA_superfamily | SCOPE | 0.6872 | 0.7779 | 0.6747 |
| HA_superfamily | Squidly | 0.4579 | 0.2576 | 0.3140 |
| EF_superfamily | SCOPE | 0.6617 | 0.6653 | 0.5727 |
| EF_superfamily | Squidly | 0.4763 | 0.2896 | 0.3416 |
| PC | SCOPE | 0.7524 | 0.8694 | 0.7473 |
| PC | Squidly | 0.4601 | 0.2620 | 0.3357 |
| EF_fold | SCOPE | 0.7048 | 0.6922 | 0.6069 |
| EF_fold | Squidly | 0.4724 | 0.2680 | 0.3483 |

**Table S31.** SCOPE and published EasIFA weights.

| **Dataset** | **Model** | **AUPRC** | **AP** | **Recall** | **F1** |
| --- | --- | --- | --- | --- | --- |
| NN | SCOPE | 0.8382 | 0.8634 | 0.9265 | 0.8019 |
| **NN** | EasIFA | 0.6285 | 0.6802 | 0.8480 | 0.6475 |
| **HA_superfamily** | SCOPE | 0.6932 | 0.7333 | 0.7952 | 0.6972 |
| **HA_superfamily** | EasIFA | 0.6228 | 0.6688 | 0.8000 | 0.6092 |
| **EF_superfamily** | SCOPE | 0.8352 | 0.8385 | 0.6000 | 0.7500 |
| **EF_superfamily** | EasIFA | 0.7932 | 0.8017 | 0.6000 | 0.7500 |
| **PC** | SCOPE | 0.8882 | 0.8987 | 0.8731 | 0.8298 |
| **PC** | EasIFA | 0.6609 | 0.7026 | 0.8090 | 0.7312 |
| **EF_fold** | SCOPE | 0.5139 | 0.6389 | 1.0000 | 0.8571 |
| **EF_fold** | EasIFA | 0.9028 | 0.9167 | 1.0000 | 0.4286 |

**CataloDB evaluation and a training restricted BLAST baseline**

We used the original CataloDB release archived at Zenodo 10.5281/zenodo.15541320. The official training and test files contain 5093 and 232 records, respectively, corresponding to 5035 and 231 unique sequences, with no exact sequence overlap between these official partitions. Eligibility required valid labels, a sequence-matched structure and length≤1000. Feature preparation and validation succeeded for 5108 records:4417 training,490 validation and 201 test records.

The archived Squidly CataloDB predictions covered 199 of the eligible test records. EasIFA predictions covered 121 records with available exact-sequence-matched reaction inputs; distinct available reactions were averaged equally without selecting reactions according to test performance. Three-model comparisons therefore used 121 records representing 120 unique sequences. SCOPE-Squidly comparisons additionally used 199 records representing 198 unique sequences. The 201 SCOPE records represent 200 unique sequences.

BLASTP queried the 201 eligible test sequences against only the 4417 records used to train SCOPE. Validation and test sequences were excluded from the reference library. Unaligned and no-hit positions were negative. Seventy-three queries had a qualifying hit. These are binary transferred labels rather than calibrated probabilities; trapezoidal AUPRC and average precision are therefore reported separately. This benchmark evaluates the specified label-transfer strategy, not every possible homology-based method.

All compared predictions were aligned to the same sequence positions and reference labels within each reported subset. The neural-model classification threshold was 0.5, fixed without test-set tuning. We report record-macro, exact-sequence-group-macro and pooled residue metrics separately. For the primary CataloDB macro analysis, record metrics were first averaged within exact-sequence groups and groups then received equal weight. A sensitivity analysis excluded exact sequence matches to distributed EasIFA supervised pretraining and fine-tuning training/validation tables. Of 121 common records,109 had such a known match, leaving 12 records.

On the common CataloDB cohort of 121 records representing 120 equally weighted sequence groups, retrained SCOPE achieved macro-AUPRC and average precision (AP) of 0.6881 and 0.7322, respectively, compared with 0.6080 and 0.6410 for Squidly, 0.5775 and 0.6507 for EasIFA, and 0.5909 and 0.2250 for the training-restricted BLAST baseline. SCOPE achieved higher ranking metrics under these conditions, whereas Squidly achieved higher F1 and EasIFA achieved higher recall (**Table S32**).

Paired sequence-group analyses supported positive AUPRC and AP differences between SCOPE and each comparator on this common cohort. The AUPRC differences relative to Squidly, EasIFA and BLAST were 0.0801, 0.1106 and 0.0972, respectively, with Holm-adjusted P values of 0.003567, 0.02235 and 0.008354. All six ranking comparisons remained significant after correction across the common-cohort comparison family (**Table S33**).

Expanded and sensitivity evaluations used separately defined cohorts. In the SCOPE-Squidly comparison covering 199 records and 198 sequence groups, macro-AUPRC was 0.6976 for SCOPE and 0.6220 for Squidly. Across all 201 input-eligible records representing 200 sequence groups, SCOPE and BLAST achieved macro-AUPRC of 0.6960 and 0.6033 and AP of 0.7367 and 0.2333, respectively. Excluding known exact matches to the available EasIFA supervised development data left 12 records; their results are reported separately (**Table S34**). These cohorts overlap and are not independent replications. BLAST produces binary label-transfer scores, for which trapezoidal AUPRC and AP can differ substantially; the two metrics are therefore reported separately.

SCOPE and SCREEN were also retrained using the same CataloDB training and validation split. Across the 201 input-eligible official test records, record-macro AUPRC was 0.6975 for SCOPE and 0.6873 for SCREEN, while pooled AUPRC was 0.6782 and 0.6741, respectively (**Table S35**).

**Table S32.** Common-cohort sequence-group macro performance.

| **Model** | **AUPRC** | **AP** | **Recall** | **F1** |
| --- | --- | --- | --- | --- |
| SCOPE | 0.6881 | 0.7322 | 0.5674 | 0.5437 |
| Squidly | 0.6080 | 0.6410 | 0.6556 | 0.5881 |
| EasIFA | 0.5775 | 0.6507 | 0.8799 | 0.4579 |
| BLAST | 0.5909 | 0.2250 | 0.2750 | 0.2544 |

**Table S33.** Paired ranking differences on the common cohort.

| **SCOPE minus** | **Metric** | **Difference** | **95% CI** | **Holm P** |
| --- | --- | --- | --- | --- |
| Squidly | AUPRC | 0.0801 | 0.0211 to 0.1367 | 0.003567 |
| Squidly | AP | 0.0912 | 0.0421 to 0.1415 | 0.001145 |
| EasIFA | AUPRC | 0.1106 | 0.0269 to 0.1931 | 0.02235 |
| EasIFA | AP | 0.0815 | 0.0133 to 0.1496 | 0.02235 |
| BLAST | AUPRC | 0.0972 | 0.0298 to 0.1615 | 0.008354 |
| BLAST | AP | 0.5072 | 0.4369 to 0.5775 | 3.47×10⁻¹⁷ |

**Table S34.** Expanded and development-excluded evaluations.

| **Subset** | **Records / sequences** | **Model** | **AUPRC** | **AP** | **Recall** | **F1** |
| --- | --- | --- | --- | --- | --- | --- |
| SCOPE_Squidly | 199/198 | SCOPE | 0.6976 | 0.7385 | 0.5918 | 0.5692 |
| SCOPE_Squidly | 199/198 | Squidly | 0.6220 | 0.6556 | 0.6511 | 0.5976 |
| SCOPE_EasIFA | 121/120 | SCOPE | 0.6881 | 0.7322 | 0.5674 | 0.5437 |
| SCOPE_EasIFA | 121/120 | EasIFA | 0.5775 | 0.6507 | 0.8799 | 0.4579 |
| Development excluded | 12/12 | SCOPE | 0.7750 | 0.8211 | 0.7778 | 0.6556 |
| Development excluded | 12/12 | Squidly | 0.7738 | 0.8135 | 0.8472 | 0.6917 |
| Development excluded | 12/12 | EasIFA | 0.7430 | 0.7837 | 0.9722 | 0.4792 |
| SCOPE_full | 201/200 | SCOPE | 0.6960 | 0.7367 | 0.5908 | 0.5685 |
| All eligible | 201/200 | BLAST | 0.6033 | 0.2333 | 0.2675 | 0.2562 |

**Table S35.** SCOPE and SCREEN retrained on the common split.

| **Model** | **Aggregation** | **AUPRC** | **AP** | **Recall** | **F1** |
| --- | --- | --- | --- | --- | --- |
| SCOPE | Record macro | 0.6975 | 0.7380 | 0.5929 | 0.5702 |
| SCOPE | Pooled | 0.6782 | 0.6800 | 0.6246 | 0.6749 |
| SCREEN | Record macro | 0.6873 | 0.7238 | 0.5647 | 0.5527 |
| SCREEN | Pooled | 0.6741 | 0.6752 | 0.5731 | 0.6678 |

**Restricted evaluation by structural similarity**

The sequence-identity sensitivity analysis is described in “Sensitivity analysis under stricter train-test sequence-identity thresholds”. Here, we report the complementary structural restriction. For each test record, structural similarity was defined as the maximum, over the 1054 development references, of the smaller of query- and reference-normalized TM-scores. Records with a value at most 0.5 were retained. Frozen SCOPE and SCREEN predictions were compared on matched positions; weights were not retrained for this analysis. Metrics are per-record macro averages.

Twenty-two records met the structural criterion. Relative performance varied by dataset, with SCOPE showing higher AUPRC in EF-superfamily and SCREEN in EF-fold, HA-superfamily and NN. The subsets are small and do not establish consistent superiority or generalization to all unseen folds. Results are summarised in **Table S36**; CATH family-disjoint retraining is reported separately in **Table S25**.

**Table S36.** Frozen-model performance on structurally restricted test subsets.

| **Dataset** | **Model** | **N** | **AUPRC** | **Recall** | **F1** |
| --- | --- | --- | --- | --- | --- |
| EF-fold | SCOPE | 4 | 0.4467 | 0.5625 | 0.4375 |
| EF-fold | SCREEN | 4 | 0.4543 | 0.5000 | 0.5000 |
| EF-superfamily | SCOPE | 1 | 0.5982 | 0.5000 | 0.3333 |
| EF-superfamily | SCREEN | 1 | 0.2875 | 0.5000 | 0.5000 |
| HA-superfamily | SCOPE | 15 | 0.2013 | 0.2444 | 0.1768 |
| HA-superfamily | SCREEN | 15 | 0.3232 | 0.2333 | 0.2200 |
| NN | SCOPE | 2 | 0.6190 | 0.7500 | 0.6667 |
| NN | SCREEN | 2 | 0.6667 | 0.5000 | 0.4000 |

Plasmid construction and mutagenesis

The tyrosine ammonia-lyase (Fc-TAL2) ^[20]^, laccase (CotA) ^[21]^, and alkaline phosphatase (rsAP) ^[22]^

used in this study for model validation shared the same gene origins as previously reported, while the diacetylchitobiose deacetylase (TGDac) was derived from the PIG-L deacetylase of *T. gammatolerans* (GenBank WP_048811066.1). Primers were synthesized and DNA sequencing was performed by Tsingke Biotechnology Co., Ltd. (Beijing, China). Recombinant plasmids for each target enzyme were constructed as follows: the codon‑optimized coding sequences were artificially synthesised and cloned into the pET-28a vector or pPICZαA, yielding the corresponding recombinant plasmids. Based on model predictions, a series of mutants were generated by whole-plasmid PCR site-directed mutagenesis. The PCR reactions were carried out using the respective recombinant plasmids as templates and 2× Phanta Max High‑Fidelity PCR Master Mix (Vazyme, Nanjing, China). The mutagenic primer sequences are listed in **Table S8**. The PCR products were digested with *Dpn*I (New England Biolabs) to remove the original methylated template, then transformed into *Escherichia coli* DH5α competent cells. After sequence verification, the mutant plasmids were transformed into *E. coli* BL21(DE3) or *Komagataella phafffi* X33 for subsequent protein expression.

Protein expression, purification and relative activity assay

The expression and purification of Fc-TAL2, CotA, and TGDac were performed following the published protocols ^[23]^. The wild‑type and mutants of alkaline phosphatase (rsAP) were expressed and purified according to the method described in previous report ^[22]^. To eliminate interference from variations in protein expression levels, all samples were normalised to the same protein concentration prior to the measurement of relative enzyme activities. Enzyme activity measurements were performed in three independent experiments.

The activities of Fc‑TAL2 and its mutants were determined as described in reference ^[24]^. The experimental procedures for CotA and its mutants followed reference ^[21]^. The activity of rsAP and its mutants was measured according to reference ^[22]^. The activity assay for TGDac was established and optimised in this study as described below: A 500 mM GlcNAc solution in PBS (pH 8.0) was prepared as the substrate. The reaction system (total volume 500 μL) consisted of 50 μL (10% of the system) of enzyme solution and 450 μL of the GlcNAc substrate solution. The enzyme solution and the substrate solution were prewarmed separately at 50 °C for 5 min. After mixing, the reaction mixture was incubated at 50 °C for 10 min, and then an equal volume of chloroform was added and mixed thoroughly to terminate the reaction. The upper aqueous phase was collected as the stock sample. To simultaneously detect the contents of the substrate and the product, the following derivatization step was performed: The stock sample was diluted 5‑fold with water, and 1 mL of the diluted solution was transferred to a 5 mL centrifuge tube, followed by the sequential addition of 600 μL of 1 mol/L sodium nitrite and 50 μL of glacial acetic acid. The mixture was allowed to react overnight at room temperature. The resulting solution was filtered through a 0.22 μm membrane to obtain the final sample for analysis. Detection was carried out using a differential refractive index (RI) detector. An Aminex HPX-87H column (300 mm × 7.8 mm, 9 μm) was used at a column temperature of 35 °C. The mobile phase was a 5 mmol/L sulfuric acid solution (degassed by ultrasonication for 15 min). Isocratic elution was performed at a flow rate of 0.4 mL/min with an injection volume of 10 μL. The activity of each mutant was expressed as a percentage of the activity of the corresponding wild-type enzyme, which was defined as 100%. Relative activities are reported as mean ± SD from three independent experiments.

**DeepGCN implementation and update equations**

For an enzyme graph with N residue nodes, the initial node feature matrix was denoted as:

$$X\in R^{N\times d}$$

where $d$ is the dimension of the residue-level input feature vector. The adjacency matrix $A$ was constructed according to the three-dimensional distances between residues, with $A_{ij}=1$ if the distance between residue $i$ and $j$ is smaller than a predefined threshold of 6Å, and $A_{ij}=0$ otherwise. The adjacency matrix was then symmetrically normalised as follows:

$$\tilde{A}=D^{-\frac{1}{2}}AD^{-\frac{1}{2}}$$

where $D$ denotes the degree matrix with diagonal elements:

$$D_{ii}=\sum_{j} A_{ij}$$

This normalisation balances the influence of nodes with different degrees and ensures numerically stable feature propagation. Residue-level self-information is preserved through residual connections and by incorporating initial features into subsequent layers.

The input node features are first linearly projected and activated to obtain the initial residue embedding:

$$h^{(0)}=\sigma(XW_{0})$$

where $W_{0}\in R^{d\times h}$ is a learnable weight matrix and $\sigma(\cdot)$denotes the ReLU activation function. For the $l-th$ layer $(l=1,...,L)$, neighborhood aggregation is first performed as

$$h_{i}=\tilde{A}h^{(l-1)}$$

An initial residual term is then introduced by linearly combining the aggregated features with the initial representation:

$$\boldsymbol{r}\boldsymbol{=(}\boldsymbol{1}\boldsymbol{-}\boldsymbol{\alpha}\boldsymbol{)}h_{i}+\alpha h^{(0)}$$

where $\alpha\epsilon(0,1)$ is a hyperparameter controlling the contribution of the initial features. In the variant setting, the aggregated representation and the initial embedding are concatenated along the feature dimension:

$$s=[h_{i}\parallel h^{(0)}]$$

and transformed by a learnable weight matrix:

$$z=sW, W\epsilon R^{2h\times h}$$

To implement a depth-dependent identity mapping, a layer-specific coefficient is defined as:

$$\theta_{l}=min(1,\log(\lambda l+1))$$

where $\lambda$ is a hyperparameter controlling the strength of identity mapping. The residue representation at layer $l$ is finally updated as

$$h^{(l)}=h^{(l-1)}+\theta_{l}z+(1-\theta_{l})\boldsymbol{r}$$

This update rule preserves information from the previous layer, maintains access to the initial residue representation and incorporates newly transformed neighbourhood features at each layer. By reducing over-smoothing during deep graph propagation, the DeepGCN module improves the stability and expressive capacity of residue-level structural learning.

**Evaluation metrics**

Binary classification was performed using the optimal decision threshold $t$, namely

$$\hat{y}_{i}=\left\{ \begin{aligned} 1, p_{i}\geq t \\ 0, p_{i}<t \end{aligned} \right.$$

where $p_{i}$ denotes the predicted probability that a residue $i$ belongs to the catalytic residue class. Based on these predictions, we calculated multiple binary classification metrics to comprehensively evaluate model performance. Accuracy is defined as the proportion of correctly classified residues among all residues:

$$Accuracy=\frac{TP+TN}{TP+TN+FP+FN}$$

where $TP$, $TN$, $FP$and $FN$ denote true positives, true negatives, false positives and false negatives, respectively. Precision measures the proportion of samples correctly classified as positive among those predicted to be positive:

$$Precision=\frac{TP}{TP+FP}$$

Recall measures the proportion of positive samples correctly identified:

$$Recall=\frac{TP}{TP+FN}$$

$F_{1}$ score is the harmonic mean of precision and recall, used to comprehensively evaluate classification performance at a single threshold:

$$F_{1}=\frac{2\times Precision\times Recall}{Precision+Recall}$$

The area under the ROC curve (AUC) is calculated by integrating the receiver operating characteristic (ROC) curve and reflects the model's overall capability to distinguish positive from negative samples across decision thresholds. The Area Under the Precision-Recall Curve (AUPRC) is calculated by integrating the Precision-Recall curve and provides a more sensitive measure of positive-class detection under class imbalance:

$$AUPRC=\int_{0}^{1} Precision(r)d(Recall(r))$$

The Matthew Correlation Coefficient (MCC) provides a robust evaluation by considering true positives, true negatives, false positives, and false negatives in scenarios with imbalanced sample categories. Its formula is as follows:

$$MCC=\frac{TP\times TN-FP\times FN}{\sqrt{(TP+FP)(TP+FN)(TN+FP)(TN+FN)}}$$

Among these metrics, $F_{1}$, Precision, Recall, and MCC assess classification performance at a specific decision threshold, while AUC and AUPRC measure discriminative ability the entire threshold range ^[25]^. Together, these metrics provide a comprehensive evaluation of the model’s predictive performance and generalizability for catalytic residue identification.

**ProtT5 embedding extraction and batching**

To improve computational efficiency and reduce memory overflow, sequences were sorted from longest to shortest and processed in batches. Batch construction was controlled by the maximum number of residues ($N_{res}$), the maximum number of sequences ($N_{seq}$) and the maximum sequence length ($L_{max}$). A forward propagation was performed when any of these conditions were met: the total number of residues reaches $N_{res}$, the number of sequences reaches $N_{seq}$, the length of any single sequence exceeds $L_{max}$, or all sequences have been processed. The model then returns the hidden state of its final layer:

$$H=ProtT5(X)\in R^{B\times L\times1024}$$

where $B$ denotes the batch size and $L$ represents the maximum sequence length within the batch. After removing padding according to the actual length of each sequence, residue-level embeddings are obtained for each enzyme. This yields a feature matrix of size $L\times1024$, which is stored in HDF5 format.

Multimodal residue feature construction

To support catalytic-residue prediction, each residue node was encoded using a multimodal feature vector that integrated evolutionary, structural, and physicochemical information. These features were aligned at the residue level to ensure positional consistency across modalities.

Evolutionary features were extracted using PSI-BLAST and HHblits ^[26]^. PSI-BLAST was run for three iterative searches against the UniRef90 database to generate position-specific scoring matrices (PSSMs) for each protein sequence. For each residue, the 20 raw PSSM scores corresponding to the standard amino acids were extracted and normalised using a sigmoid function to map values to the range [0,1], improving numerical stability while preserving relative evolutionary preferences. This yielded a 20-dimensional PSSM evolutionary feature vector for each residue. Simultaneously, HHblits was utilised to construct a Hidden Markov Model (HMM) profiles from the UniClust30 database to capture deep evolutionary patterns ^[27, 28]^. Residue-wise emission and transition features were extracted from the HHblits output and normalised using min-max scaling, generating a 30-dimensional HMM evolutionary feature vector for each residue.

Structural features were derived from three-dimensional protein structures and aggregated from atoms to residues. For each atom in the PDB file, we parsed its spatial coordinates, element type, B factor, atomic mass, and side-chain status ^[29, 30]^. Additional predefined atomic physicochemical properties were assigned, including charge state, hydrogen-bond donor or acceptor capability, aromatic-ring participation and van der Waals radius ^[31]^. These properties were normalised and combined into a seven-dimensional atomic feature vector. To obtain residue-level structural descriptors, features from all non-hydrogen atoms within the same residue were averaged, producing a seven-dimensional atomic structural feature vector for each residue.

Intrinsic amino acid features were represented using a fixed seven-dimensional lookup encoding inherited from the SCREEN implementation. Each standard amino acid was assigned its corresponding vector in the original channel order. The complete 20$\text{×}$7 numerical mapping, channel descriptions, and implementation-specific exceptions are provided in **Seven-dimensional amino acid encoding**. The stored values were used directly, without protein-specific renormalization or regeneration from molecular descriptors, to preserve consistency with the features used during model training.

After feature extraction, all features were aligned by sequence position to ensure residue-level correspondence across modalities. When atomic features were unavailable because of missing structural regions, zero vectors were used to maintain dimensional consistency. The PSSM features (20 dimensions), HMM features (30 dimensions), atomic structural features (7 dimensions), and amino acid physicochemical descriptors (7 dimensions) were then concatenated to form a 64-dimensional feature vector for each residue node. This multimodal representation combined evolutionary conservation, local three-dimensional structural information and intrinsic residue properties as input for downstream DeepGCN-based catalytic residue prediction.

Edge representations

Graph edges were defined to encode the spatial proximity between residues and capture geometric interaction patterns through distance-based representations. For each enzyme, three-dimensional residue coordinates were extracted from the corresponding PDB structure, and the α-carbon atom of each residue was used as its representative spatial position. The α-carbon atom was selected because it is present in all standard amino acids and provides a stable reference point for residue-level geometric representation. Pairwise Euclidean distances were calculated between all residue pairs to generate a residue-level distance matrix. This matrix was then converted into a contact graph by connecting two residues with an undirected edge when their α-carbon distance was less than or equal to 6Å ^[32]^. This threshold was selected to capture local residue neighbourhoods that are likely to reflect direct or indirect structural coupling, including hydrogen bonding, hydrophobic packing and electrostatic interactions. Compared with smaller thresholds, the 6Å cutoff reduces excessive graph sparsity and preserves important local structural connections. Compared with larger thresholds, it reduces noisy connections between residues that are spatially distant and less likely to participate in the same local catalytic environment (Table S26). Thus, the edge representation encodes the local three-dimensional topology of each enzyme and provides the structural basis for graph-based message passing in SCOPE.

Graph Construction

The thresholding operation converted the continuous residue-residue distance matrix into a binary adjacency matrix, in which an entry of 1 indicates a spatial connection between a residue pair and an entry of 0 indicates no connection. Symmetric normalisation was then applied to preserve the undirected structure of the graph while reasonably scaling feature propagation across nodes with different degrees. Isolated nodes with zero degree were handled separately to avoid numerical instability during normalisation and to minimise their influence on graph-based message passing.

Through distance computation, threshold-based filtering, adjacency matrix construction and symmetric normalisation, each enzyme was represented as a residue-level graph with structural coherence, numerical stability and biological relevance. This graph served as the core input to the downstream graph neural network, enabling SCOPE to aggregate information within spatial neighbourhoods and progressively model higher-order structural interactions relevant to catalytic-residue prediction.

**Supplemental References**

[1]	Torrance, J. W., Holliday, G. L., Mitchell, J. B. O. & Thornton, J. M. The geometry of interactions between catalytic residues and their substrates. *Journal of Molecular Biology* **369**, 1140-1152 (2007).

[2]	Jack, B. R., Meyer, A. G., Julian, E., Wilke, C. O. & Petsko, G. A. Functional Sites Induce Long-Range Evolutionary Constraints in Enzymes. *Plos Biology* **14**, e1002452 (2016).

[3]	Dayhoff, M. A model of evolutionary change in proteins. *Atlas of Protein Sequence & Structure* **5**, 345-352 (1978).

[4]	Kyte, J. & Doolittle, R. F. A Simple Method for Displaying the Hydropathic Character of a Protein. *Journal of Molecular Biology* **157**, 105-132 (1982).

[5]	Kristin*, et al.* Crystal structure of thymidylate synthase A from Bacillus subtilis. *Protein Science*, (1999).

[6]	Cunane, L. M.*, et al.* Structures of the flavocytochrome p-cresol methylhydroxylase and its enzyme-substrate complex: gated substrate entry and proton relays support the proposed catalytic mechanism. *Journal of Molecular Biology* **295**, 357-374 (2000).

[7]	Bompardgilles, C.*, et al.* Crystal structure of a D-aminopeptidase from Ochrobactrum anthropi, a new member of the 'penicillin-recognizing enzyme' family. *Structure* **8**, 971-980 (2000).

[8]	Saadat, D. & Harrison, D. The crystal structure of methylglyoxal synthase from Escherichia coli. *Structure (London, England : 1993)* **7**, 309-317 (1999).

[9]	Jabri, E. & Karplus, P. A. Structures of the Klebsiella aerogenes urease apoenzyme and two active-site mutants. *Biochemistry* **35**, 10616-10626 (1996).

[10]	Fox, K. M., Maley, F., Garibian, A., Changchien, L. M. & Roey, P. V. Crystal structure of thymidylate synthase A from Bacillus subtilis. *Protein Science* **8**, 538-544 (1999).

[11]	Ying, R., Bourgeois, D., You, J., Zitnik, M. & Leskovec, J. GNNExplainer: Generating Explanations for Graph Neural Networks. *Adv Neural Inf Process Syst* **32**, 9240-9251 (2019).

[12]	Ribeiro, A. J. M., Tyzack, J. D., Borkakoti, N., Holliday, G. L. & Thornton, J. M. A global analysis of function and conservation of catalytic residues in enzymes. *J Biol Chem* **295**, 314-324 <https://doi.org/10.1074/jbc.REV119.006289> (2020).

[13]	Turnbull, A. P.*, et al.* Analysis of the structure, substrate specificity, and mechanism of squash glycerol-3-phosphate (1)-acyltransferase. *Structure* **9**, 347-353 (2001).

[14]	Wang, J., Hartling, J. A. & Flanagan, J. M. The structure of ClpP at 2.3 A resolution suggests a model for ATP-dependent proteolysis. *Cell* **91**, 447-456 (1997).

[15]	Romão, M. J.*, et al.* Crystal structure analysis, refinement and enzymatic reaction mechanism of N-carbamoylsarcosine amidohydrolase from Arthrobacter sp. at 2.0 A resolution. *J Mol Biol* **226**, 1111-1130 <https://doi.org/10.1016/0022-2836(92)91056-u> (1992).

[16]	Newmann, M.*, et al.* X-ray analyses of aspartic proteinases. V. Structure and refinement at 2.0 A resolution of the aspartic proteinase from Mucor pusillus. *Journal of Molecular Biology* **230**, 260-283 (1993).

[17]	Dreyer, M. K. & Schulz, G. E. Refined high-resolution structure of the metal-ion dependent L-fuculose-1-phosphate aldolase (class II) from Escherichia coli. *Acta Crystallographica* **52**, 1082-1091 (2010).

[18]	Bond, C. S., White, M. F. & Hunter, W. N. High Resolution Structure of the Phosphohistidine-activated Form of Escherichia coli Cofactor-dependent Phosphoglycerate Mutase. *Journal of Biological Chemistry* **276**, 3247-3253 (2001).

[19]	Webb, E. C. Enzyme nomenclature. Recommendations 1984. *European Journal of Biochemistry* **187**, 263-281 (1990).

[20]	Wang, J., Zhang, N., Huang, Y., Li, S. & Zhang, G. Simple and efficient enzymatic procedure for p-coumaric acid synthesis: Complete bioconversion and biocatalyst recycling under alkaline condition. *Biochemical Engineering Journal* **188**, 108693 <https://doi.org/10.1016/j.bej.2022.108693> (2022).

[21]	Sun, X.*, et al.* Engineering Bacterial Laccase with Improved Catalytic Activity and Thermostability by Rational Design. *Applied Biochemistry and Biotechnology* **197**, 4685-4701 <https://doi.org/10.1007/s12010-025-05240-1> (2025).

[22]	Xie, Y.*, et al.* A highly efficient alkaline phosphatase for dephosphorylation of macromolecular substrates. *Int J Biol Macromol* **352**, 151134 <https://doi.org/10.1016/j.ijbiomac.2026.151134> (2026).

[23]	Pan, T.*, et al.* TraSPIN: inferring transporter specificity through language model-guided graph learning. *Genome Biology*, <https://doi.org/10.1186/s13059-026-04309-2> (2026).

[24]	HUANG, Y., JIANG, X., CHEN, W., ZHANG, G. & WANG, Q. Characterization of highly active tyrosine ammonia lyase and its application in biosynthesis of p-coumaric acid. *Chinese Journal of Biotechnology* **38**, 4553-4566 <https://doi.org/10.13345/j.cjb.220182> (2022).

[25]	Hao, Y., Guo, X., Ran, Z., Bi, Y. & Li, F. LncTracker: A Unified Multi-Channel Framework for Multi-Label lncRNA Localization. *IEEE Journal of Biomedical and Health Informatics* **30**.

[26]	Altschul, S. F.*, et al.* Gapped BLAST and PSI-BLAST: a new generation of protein database search programs. *Nucleic Acids Res* **25**, 3389-3402 <https://doi.org/10.1093/nar/25.17.3389> (1997).

[27]	Potter, S. C.*, et al.* HMMER web server: 2018 update. *Nucleic Acids Res* **46**, W200-w204 <https://doi.org/10.1093/nar/gky448> (2018).

[28]	Mirdita, M.*, et al.* Uniclust databases of clustered and deeply annotated protein sequences and alignments. *Nucleic Acids Res* **45**, D170-d176 <https://doi.org/10.1093/nar/gkw1081> (2017).

[29]	Bernstein, F. C.*, et al.* The Protein Data Bank: a computer-based archival file for macromolecular structures. *J Mol Biol* **112**, 535-542 <https://doi.org/10.1016/s0022-2836(77)80200-3> (1977).

[30]	Ran, Z., Wang, C., Sun, H., Pan, S. & Li, F. Characterizing Secretion System Effector Proteins With Structure-Aware Graph Neural Networks and Pre-Trained Language Models. *Journal on Biomedical and Health Informatics (J-BHI)* **28**, 9 (2024).

[31]	Silva, J. C. F.*, et al.* InteracTor: Feature engineering and explainable AI for profiling protein structure-interaction-function relationships. *PLoS Computational Biology* **21**, (2025).

[32]	Plaxco, K. W., Simons, K. T. & Baker, D. Contact order, transition state placement and the refolding rates of single domain proteins. *J Mol Biol* **277**, 985-994 <https://doi.org/10.1006/jmbi.1998.1645> (1998).
